# Frustration in the Protein-Protein interface Plays a Central Role in the Cooperativity of PROTAC Ternary Complexes

**DOI:** 10.1101/2024.12.03.626645

**Authors:** Ning Ma, Supriyo Bhattacharya, Sanychen Muk, Zuzana Jandova, Philipp S. Schmalhorst, Soumadwip Ghosh, Keith Le, Emelyne Diers, Nicole Trainor, William Farnaby, Michael J. Roy, Christiane Kofink, Peter Greb, Harald Weinstabl, Alessio Ciulli, Gerd Bader, Kyra Sankar, Andreas Bergner, Nagarajan Vaidehi

## Abstract

Targeted protein degradation of a protein of interest (POI) by **Pro**teolysis **Ta**rgeting **C**himeras (PROTACs) is an attractive approach for dealing with formerly undruggable protein targets. PROTACs are heterobifunctional molecules that connect a POI-binding and an E3-ligase (E3) binding motif with a linker. The simultaneous binding and formation of a ternary POI::PROTAC::E3 complex (TC) induces proximity between the POI and the E3, allowing for POI-ubiquitination and subsequent induction of proteasomal degradation. Despite the availability of many three-dimensional structures of TCs, unveiling the structure-function relationships for the design of PROTACs remains a challenge. This is because the TCs can be dynamic with a complex conformational landscape that individual crystal structures may not capture. In this work, we used SMARCA2 as the POI and VHL as the E3-ligase and solved the X-ray crystal structures of the respective ternary complexes with four different PROTACs. Molecular dynamics (MD) simulations were used to show that the SMARCA2-VHL interface is flexible with multiple energy minima. The protein-protein (POI-E3) interactions are largely formed by residues located in structurally disordered loops in both VHL and SMARCA2. The residue pairs in the SMARCA2-VHL interface are ‘frustrated’, i.e., adopt a suboptimal energetic state. The number of frustrated residue pairs averaged over the MD ensemble shows a positive correlation with the experimentally determined cooperativity of the PROTACs. This indicates that protein-protein interface frustration can play an important role in PROTAC function. The TC ensembles of VHL, SMARCA2 and 11 different PROTACs were modeled by comparative modeling followed by MD. The frustration was subsequently calculated from the MD trajectories and correlated with the cooperativity. We found that identification of the dynamic protein-protein contacts and frustrated residue pairs in the interface can provide a rational framework for the structure-based design of PROTACs.

## Introduction

Proteolysis-targeting chimeras (PROTACs) are an emerging class of therapeutics that have opened opportunities for targeting a wider range of proteins, as they enable an alternative mode of action when compared to conventional small molecule drugs, e.g. inhibitors^1^. PROTACs are heterobifunctional molecules consisting of two binding motifs each of which binds to the POI and an E3 ubiquitin ligase (E3) and are joined by a linker. Upon formation of a ternary complex (TC) composed of the POI, the PROTAC and the E3 ligase (POI::PROTAC::E3), the E3 catalyzes the transfer of ubiquitin from the E2 ubiquitin conjugating ligase to the POI. The ubiquitinylated POI is then subjected to the ubiquitin– proteasomal degradation machinery, leading to the removal of the POI from the cell^2^. Unlike conventional protein inhibitors that rely on an occupancy-driven mode of action, PROTACs operate through a different mechanism where only a brief interaction is needed to trigger the target protein’s degradation, essentially acting as a "catalytic" event rather than relying on constant occupancy. Very often, the ternary complex formation by a PROTAC is mediated through proximity-induced formation of *de novo* interactions within the complex^3^. The tendency of a PROTAC to stabilize such a ternary complex can be quantified using the cooperativity value α, i.e., the binding affinity ratio of a PROTAC in the binary complex (PROTAC::POI, PROTAC::E3) versus the ternary complex. The cooperativity is positive (α > 1) when the binding affinity to the ternary complex is greater than the affinities to the binary complexes, thus stabilizing the ternary complex. The cooperativity is negative (α < 1) when the binding affinity of the PROTAC to the ternary complex is lower than to the binary complexes, hence destabilizing the ternary complex. A non-cooperative cooperativity (α = 1) means no change in the binding affinity for the ternary complex when compared to the binary complexes^4,5^.

Although the three-dimensional structures of TCs inform us about the nature of the POI-E3 interface^6–9^, structure-based design of PROTACs still remains challenging. This is in part because some ternary complexes are highly dynamic^10–13^, and the crystal structure of an individual ternary complex represents only one of, in some cases, many energy minima in the energy landscape of the TC conformational ensemble. There has been a lack of computational metrics to assess cooperativity of PROTACs prior to synthesis due to the highly dynamic energy landscape of the TCs.

Computational methods play a critical role in understanding the behavior of the PROTAC-mediated TCs. Multiple methods for the modelling or prediction of ternary complex structures have been published^14,15,16^, many of which are based on protein-protein docking. A recent study used weighted ensemble molecular dynamics (MD) simulations to model the formation of the ternary complex ensembles^17^. However, no descriptors derived from the PROTAC ternary complex ensembles have yet been discovered that correlate with the cooperativity, even when the 3D-structures of the ternary complexes are known. Often, the protein-protein interface (PPI) residues of the POI and the E3 ligase are in unstructured or disordered regions, and hence are highly dynamic^10,11,18^. Furthermore, PROTAC mediated PPIs are not driven through coevolution of interface residues which, in naturally occurring PPIs, is often the basis of the PPI stabilization. Given the lack of correlation between calculated binding affinities of PROTACS and cooperativity^17^, we looked for structural descriptors that characterize the PPIs in unstructured regions and their role in regulating the protein activity. Previous studies have shown that distinct residue positions which play a critical functional role are often in a suboptimal energetic state before being involved in intramolecular or intermolecular protein interaction^12^. The presence of such locally perturbed thermodynamic states in proteins is referred to as ‘frustration’^9,15^.

Frustration was developed as a concept to quantify the conformational variations and heterogeneity often seen in dynamic PPIs formed through disordered regions^19,20^. Since the TCs are highly dynamic, we hypothesized that the frustration of the interactions at the POI-E3 ligase interface could be a relevant factor in determining the cooperativity of the ternary complex.

We investigated a series of PROTACs that mediate the degradation of SMARCA2 as the POI through recruitment of the E3-ligase VHL, to assess whether frustration could predict ternary complex stability. The helicases SMARCA2 and SMARCA4 are close paralogs and crucial components of the BAF chromatin remodeling complex. Based on a synthetic lethality concept, SMARCA2 has been proposed as a drug target for certain tumors exhibiting SMARCA4-deficiency (e.g. non-small cell lung cancer (NSCLC))^21^. The von Hippel-Lindau tumor suppressor (VHL) is an E3 ubiquitin ligase in which small molecule binders with suitable exit vectors for PROTAC design have been described before^22^. Farnaby, Koegl et al. utilized a SMARCA bromodomain (BD) binder reported by Genentech (GEN-1)^23^ to synthesize the first VHL-based SMARCA2/4-degrading PROTACs and solved three ternary complex crystal structures to guide PROTAC design^8^. Later, Kofink, Trainor, Mair et al. discovered alternative SMARCA^BD^ binders with improved physicochemical properties. Guided by a further three TC crystal structures, SMARCA2-selective PROTACs with oral bioavailability could be designed^9^.

In this study, we solved the X-ray crystal structures of SMARCA2-VHL complexes with four different PROTACs to a resolution ranging from 2.2 Å to 3.74 Å. Starting from the crystal structures of five TCs with SMARCA2 as the POI and VHL as the E3 ligase, we performed all-atom MD simulations to study the TC dynamics. We calculated the frustration of all residue pairs in the POI:E3 interface and demonstrated that frustration correlates with the cooperativity. We further modelled the TCs of 11 GEN-1 based PROTACs using comparative modeling and calculated the frustration in the modelled SMARCA2-VHL PPIs. We found that the average number of frustrated residue pairs in the SMARCA2-VHL interface is higher for PROTACs with high cooperativity than those with low cooperativity. Thus, we speculate that the frustration can be used as a measure for cooperativity for prioritization of PROTACS prior to synthesis, for cases where the POI-E3 pairing supports cooperativity as a relevant optimization parameter.

## Results

### Synthesis of PROTACs for SMARCA2-VHL complexes and measurement of cooperativity

We investigated 16 PROTACs (shown in Fig. 1A and Fig. S1) in this work and measured their cooperativity (α). Eleven of these PROTACs, P6 to P20, are based on the original GEN-1 SMARCA^BD^ binder^23^, while P1 to P5 incorporate alternative SMARCA binder versions^9^. All PROTACs contain the same VHL binder, VH101 (compound 10)^22^, except for P3 in which the VH101 fluorine was replaced by a dimethyl amino function^9^. A phenolic hydroxyl group on VH101 serves as exit vector to the linker, except for P2, P3 and P18, which use the benzylic position instead. P19 and P20 contain the *cis*-diastereomer of the hydroxy-pyrrolidine moiety in VH101, rendering the binder inactive to VHL^8^. Seven PROTACs, P1, P3, P4, P9, P10, P11 and P18, are novel disclosures and their synthesis is described in the Supporting Information.

**Figure 1.**
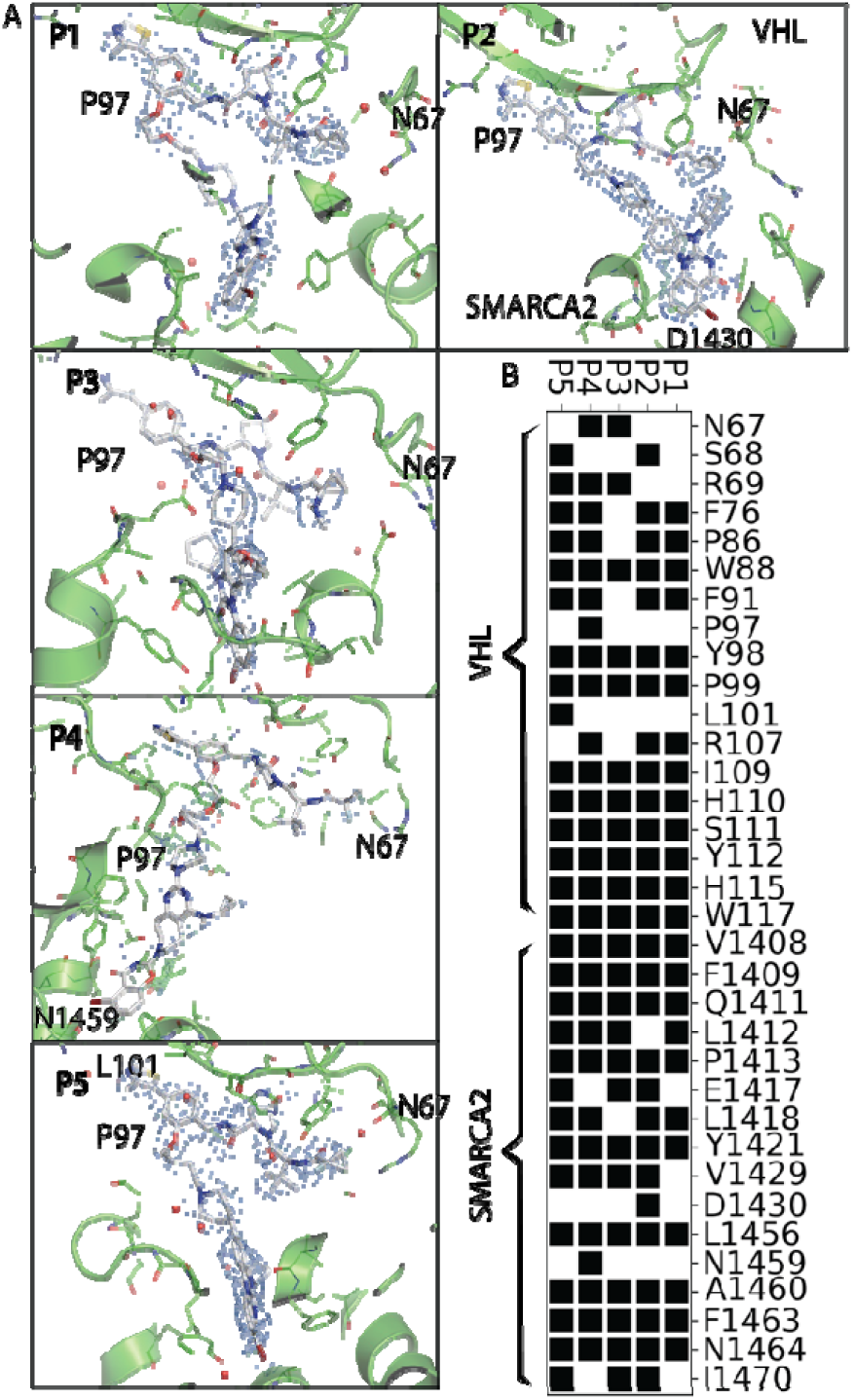
Ternary complex crystal structures of SMARCA2^BD^, VCB and PROTACs P1 to P5. A) The VHL-binding part of the PROTACs is oriented in similar positions. SMARCA2^BD^ and VHL are shown in green, PROTACs in white. The electron density map of the refine 2f_o_f_c_ map i depicted at a contour level of 1.0 σ for the PROTACs. B) Protein residues that interact with PROTAC in the five crystal structures include VHL residues with indices below 1000 and SMARCA2 residues with indices above 1000.

A previously developed time-resolved fluorescence resonance energy transfer (TR-FRET) competition assay was used to assess the cooperativity of PROTAC binding to SMARCA2 and VHL^8^. Briefly, a biotinylated SMARCA2 probe^23^ was incubated with the His_6_- tagged SMARCA2^BD^ protein and the proximity of both components was measured from a FRET donor/acceptor pair tagged to streptavidin or an anti-His antibody, respectively. The loss of the proximity signal, after addition of a concentration series of each PROTAC in the presence or absence of saturating concentrations of VCB (a pre-formed complex of VHL, Elongin-C, and Elongin-B), yielded the ternary and binary IC_50_, respectively. The cooperativity (α) was defined as the ratio: IC_50_(binary) / IC_50_(ternary), so that the affinity gains of a PROTAC to SMARCA2 after addition of VCB, i.e., a lower ternary IC_50_, lead to higher α values. Binary and ternary IC_50_ values, as well as α values for all 16 PROTACs, are reported in Table S1.

### Crystal structures of five PROTAC ternary complexes

For two of the 16 PROTACs, P2 and P13, ternary complex crystal structures have previously been published (PDB codes 7Z77^9^ and 6HAY ^8^, respectively). We solved the crystal structures of the PROTAC ternary complexes: P1, P3, P4 and P5, to enable comparison of their binding mode with existing crystal structures (Fig. 1A and Tables S3 to S6). In all cases, the SMARCA binders bound in the acetyl-lysine binding site as previously described, i.e., with the key interactions of the quinazolinone (P4: benzoxazinone) core to Leu1456 and Asn1464 of SMARCA2^BD^ ^9^. The VHL binder was consistently found to bind as described previously^22^. The interactions between SMARCA2^BD^ and VHL varied substantially across the PROTACs, as detailed later in the manuscript.

### Comparison of the crystal structures of SMARCA2-VHL PROTAC ternary complexes show that the ternary complexes are dynamic

The cooperativity of the five PROTACs, P1 to P5, ranges from 0.2 to 93 (see Fig. 2A, table S1). Structural alignment of the five TC crystal structures onto the VHL domain shows the high flexibility of the ternary complexes: SMARCA2 and VHL adopt very different conformations relative to each other with different PROTACs (Fig. 2B). These differences can stem both from the differences in the SMARCA2 binding motif and/or the linker lengths of the PROTACs. The root mean square deviation (RMSD) in the coordinates of the backbone atoms of the SMARCA2 domain ranges from 2.8 Å to 15.6 Å when aligned by the VHL domain of the P1 ternary complex crystal structure. The SMARCA2 domain orientations vary even between the two low cooperativity PROTACs (P4, α =1.3 and P5, α =0.2).

**Figure 2:**
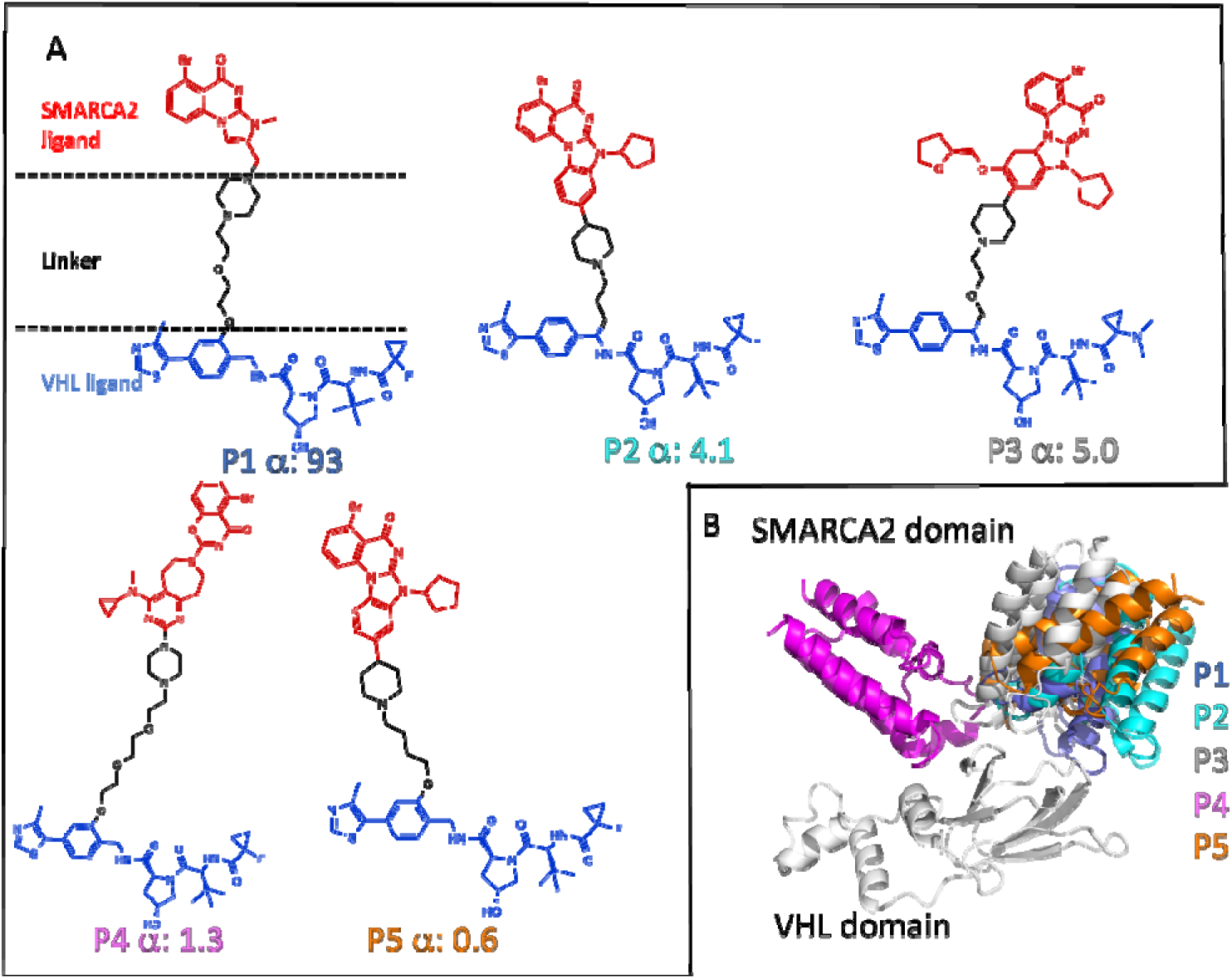
(A) 2D Chemical structures of the five PROTACs ordered by decreasing cooperativity. (B) Crystal structures of the five PROTAC-bound ternary complexes aligned onto their VHL domains.

### MD simulations show that each PROTAC ternary complex samples multiple conformational states

Starting from the respective crystal structures we performed all-atom MD simulations with the ternary complexes of the five PROTACs shown in Fig 2A, for 3000 ns each using AMBER18 MD simulation package^24^ (details in the Methods section). To analyze and visualize the dominant motions, Principal Component Analysis (PCA) was performed on the combined trajectories of all five complexes (see Methods for details). The top three weighted principal components capture nearly 80% of the variance (as shown in Fig. S2 of the Supporting Information). Principal component 1 (PC1) represents a rotation of the SMARCA2 domain with respect to the VHL domain, while PC2 represents a swinging motion of the SMARCA2 with respect to VHL (Fig. 3A, movies 1 and 2 in the Supporting Information). Projection of the snapshots from MD simulations of all the ternary complexes (Fig. 3B and Fig. S3) in the PC1 and PC2 coordinates shows that the high cooperativity PROTAC P1 complex samples a relatively compact conformational ensemble compared to the broader ensembles of the low cooperativity PROTAC complexes, for example P4 and P5. The respective crystal structures thus only represent single low energy snapshots in this wide conformational space as seen in Fig. 3B.

**Figure 3:**
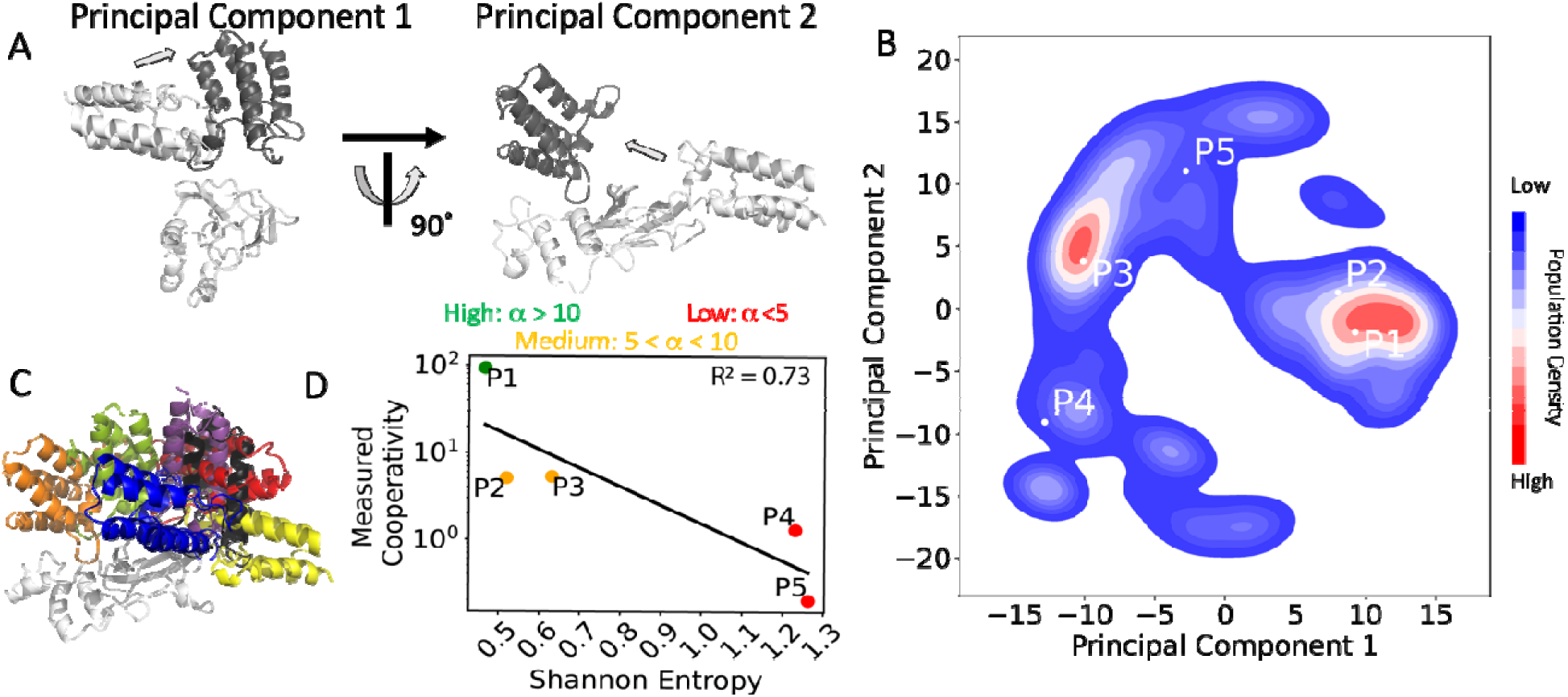
(A) Domain motions represented by the Principal Component 1 and 2 (PC1 and PC2) resulting from PC analysis of the MD trajectories of all the five PROTAC ternary complexes. (B) Projections of conformations from the MD simulations onto the PC1-PC2 space shown in a contour plot; the respective crystal structures are marked by white dots and labeled P1-P5. (C) representative conformations from the highest populated cluster of each from the PROTAC bound TC MD simulations overlaid, after aligning their VHL domains. (D) Scatter plot of Shannon entropy (x-axis) versus the experimentally determined cooperativity values (y-axis) are shown as dots. PROTACs with α > 10 are colored in green, those with 5 < α < 10 in orange, and α < 5 cases are colored in red.

We clustered the conformations from the MD simulation trajectories of each ternary complex by PC1 and PC2 and Table S2 shows the number of conformation clusters and their relative population for all five complexes. Each TC shows multiple conformation clusters indicative of their dynamic nature. The PROTACs with high and medium cooperativity: P1, P2 and P3 show fewer conformational clusters when compared to the PROTACs with low and negative cooperativity: P4 and P5, where the ensemble population is distributed over a larger number of clusters. This indicates that PROTACs with low cooperativity impart more flexibility to the ternary complex than PROTACs with high cooperativity. Fig. 3C shows the alignment of representative structures extracted from the most populated conformational cluster (see Methods) for all five PROTACs. The positional spread of the SMARCA2 domain in relation to VHL is wider than observed when comparing only crystal structures (Fig. 2B). We then calculated the Shannon entropy using the conformation cluster population distributions (see Methods) for the five PROTAC complexes as a measure of their conformational diversity. As seen in Fig. 3D, the Shannon entropy shows weak but significant (p value: 0.036)^25^ inverse correlation with cooperativity. The two PROTACs with low cooperativity, P4 and P5, show significantly higher Shannon entropy than the PROTACs, P1-P3, with high and medium cooperativity.

### The Spatiotemporal residue contact heat map from MD simulations reflects the cooperativity of the PROTACs

In the previous section, we analyzed the conformational diversity of TCs with five different PROTACs using MD simulations. To understand how PROTAC flexibility could have contributed to these differences, we performed backbone RMSD clustering on the SMARCA2-VHL complex conformations from the MD simulations and extracted one representative structure from each cluster (Fig. 4A and 4B). As seen in Fig. 4A, PROTAC P1 with high cooperativity showed closely related PROTAC conformations, in contrast to PROTAC P5 with low cooperativity (Fig. 4B), which showed multiple disparate conformations. The higher flexibility of the PROTACs with low cooperativity could be due to the linker structure (Fig. S5), which in turn could lead to increased flexibility in the PPI. To characterize the PROTAC flexibility further, we identified the VHL and SMARCA2 residues that are in contact with the PROTACs in the crystal structures. As seen in Fig. 4C, the number of PROTAC-protein contacts gleaned from the crystal structures are similar for all five PROTACs and do not distinguish between low and high cooperativity PROTACs. On the other hand, the contact frequency heatmap between the PROTAC and VHL or SMARCA2, as derived from MD simulation trajectories (shown in Figs. 4C and 4D), shows that the high and medium cooperativity PROTACs (P1 to P3) have a higher number of persistent PROTAC:protein contacts (P1:21, P2:23, P3:24) when compared to the low cooperativity PROTACs (P4:15, P5:19). In addition, the PROTAC contacts with SMARCA2 are varied for the different PROTACs (Fig. 4D). These frequency heatmaps reflect the dynamic PPI mediated by the PROTACs. The interaction fingerprints derived from MD simulations are therefore more indicative of PROTAC cooperativity when compared to fingerprints derived from crystal structures (Fig. 4D).

**Figure 4:**
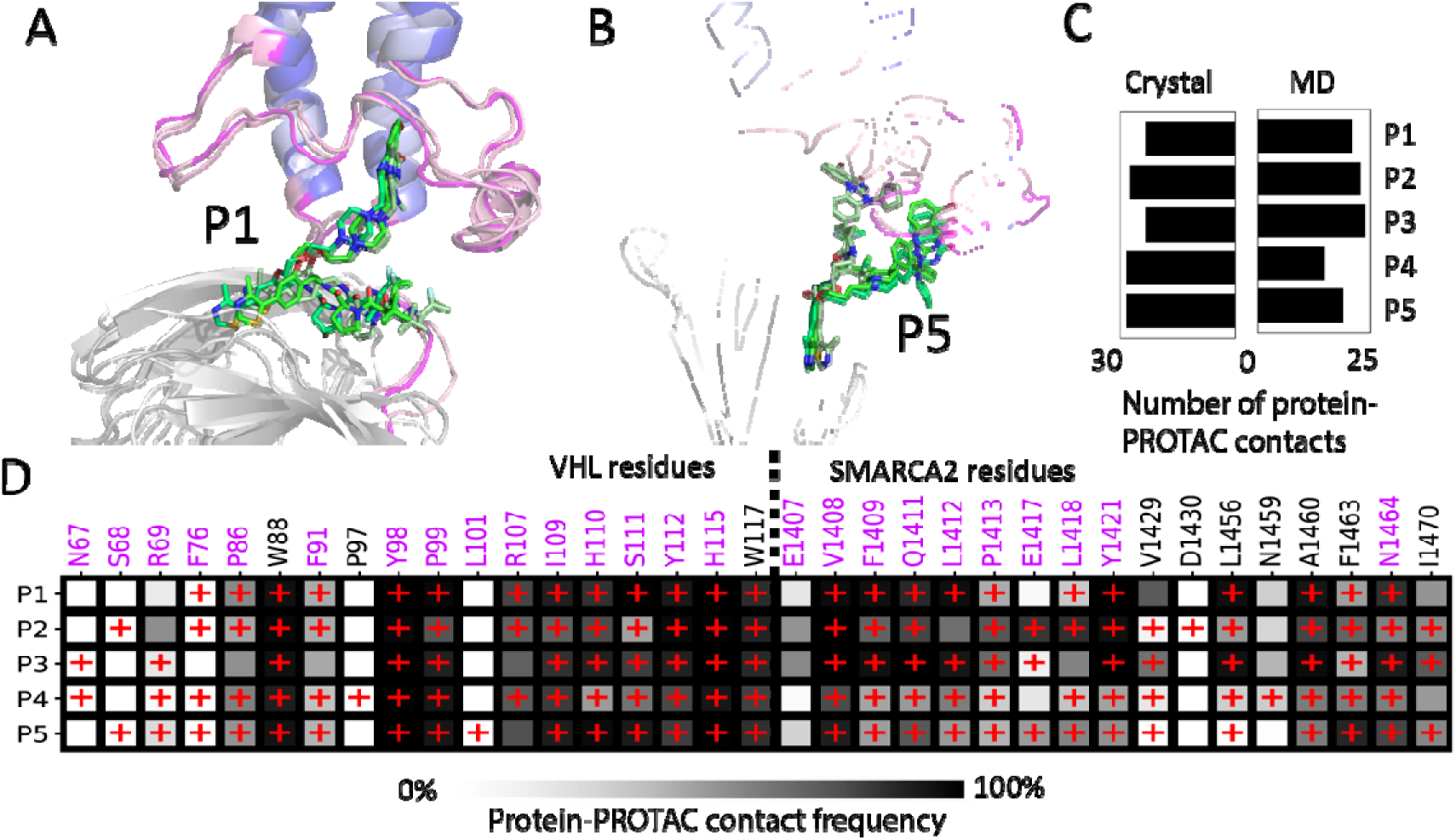
A) The representative structures of the top three most populated TC conformation clusters of P1, superimposed onto the VHL domain. The three conformations are colored from dark to light color, SMARCA2 colored in blue, VHL colored in grey, PROTAC colored in green. The unstructured loop regions are colored in pink. B) The representative structures of the three most populated clusters of P5. C) Total number of residue contacts in PROTAC-VHL and PROTAC-SMARCA2 interfaces, as observed in crystal structure and MD simulations (contact formed for >50% of simulation time). D) The contact heatmap between PROTAC and proteins (POI and E3) obtained from the MD ensembles. Red “+” symbols indicate contacts formed in crystal structures. Residue numbers for VHL are below 1000, and for SMARCA2 are above 1000. Residues located in the unstructured loop regions are shown in magenta labels.

### Residue contacts in the SMARCA2-VHL interface in PROTAC ternary complexes are dynamic and located in unstructured regions

Previous structural and biophysical studies have shown that the contact surface area between the POI and E3 ligase is critical to the cooperativity and can impact degradation properties of PROTACs^26,27^. We analyzed the pairwise residue interactions in the VHL-SMARCA2 interface, both in the crystal structures and in the MD trajectories of the five PROTACs (Fig. 5A). Comparison of the contact heatmaps in Figs. 5A and Fig. 4D, shows that most of the protein-protein interface contacts are mediated by the PROTAC, but also a few direct protein-protein contacts exist. The number of such direct contacts in the crystal structures doe not allow discrimination between the high cooperativity PROTAC P1 from the low cooperativity PROTAC P4, although the negatively cooperative PROTAC P5 shows a negligible number of SMARCA2-VHL contacts. Interestingly, during the MD simulations, additional contacts were formed and many of the contacts observed in the crystal structures were weakened. The frequency heatmap calculated from the MD simulation trajectories shows that the high and medium cooperativity PROTACs exhibit more direct PPI contacts than the low cooperativity PROTACs P4 and P5 (Fig. 5B). This loss of initial residue contacts in the SMARCA2-VHL interface can be attributed to the high flexibility of the ternary complexes. In contrast, the PROTAC-mediated SMARCA2-VHL contacts are maintained in the MD simulations demonstrating the critical role of the PROTAC in stabilizing the ternary complex. Other MD simulation studies on PROTAC ternary complexes have also shown this behavior^17,28^. We also observed that the SMARCA2-VHL interface contacts are assembled from unstructured loop regions in both VHL and SMARCA2 (POI), as shown in Figs. 5C and 5D. In summary, th number of PPI residue contacts in the crystal structures does not correlate with the cooperativity. Next, we investigated PPI properties with a focus on disordered regions, these are often in suboptimal, or ‘frustrated’, energy states^13,29^. The SMARCA2-VHL interface is dynamic and is formed by residue contacts built from the unstructured loop regions as shown in Figs. 5A, 5C and 5D. We calculated the frustration of these interface residues to examine whether or not frustration can distinguish between high and low cooperativity PROTACs.

**Figure 5:**
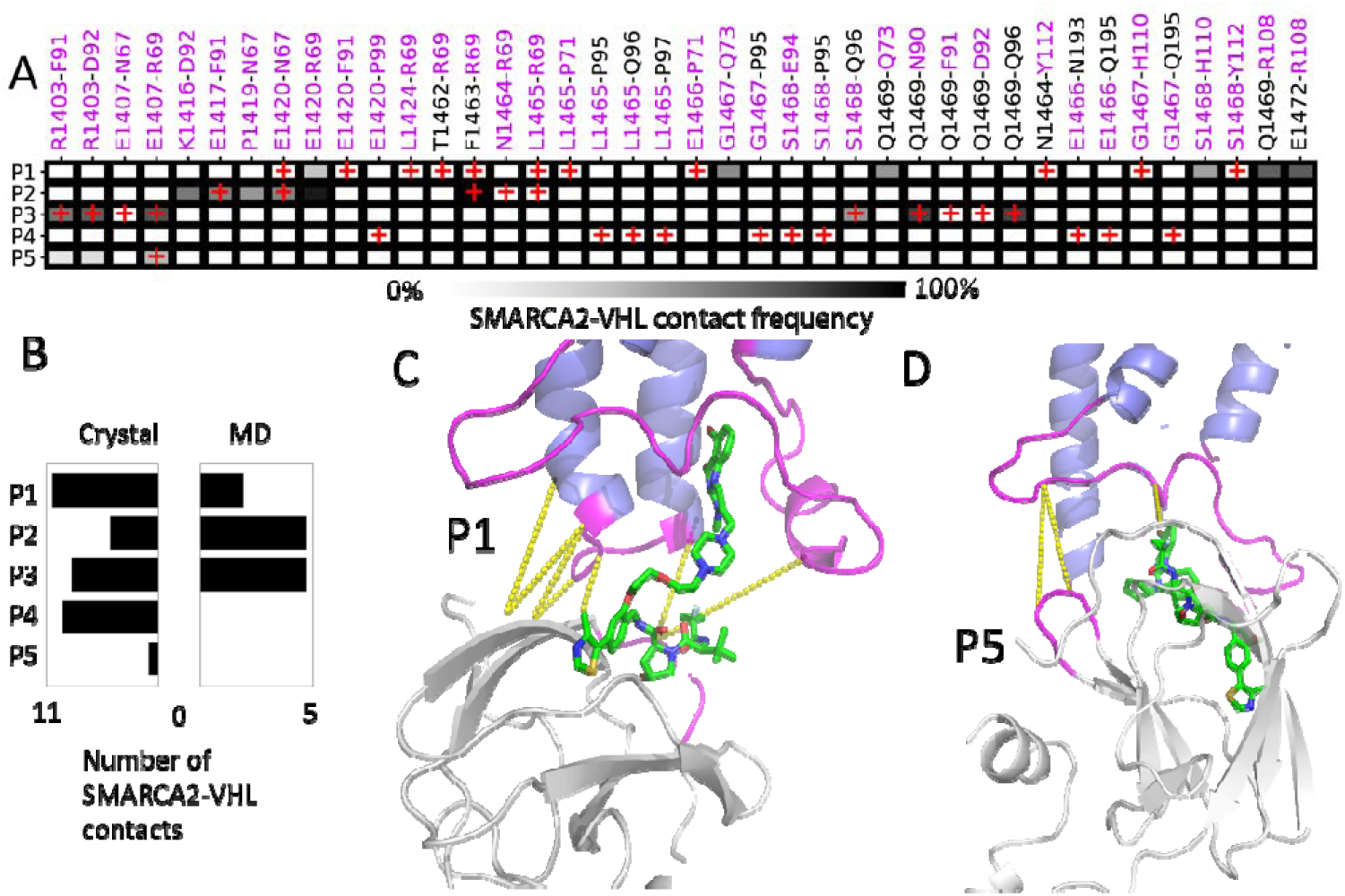
A) Heatmap of residue contacts made between SMARCA2 and VHL domains, from both crystal structures and MD simulations (contact present if formed for >50% simulation time). Red “+” signs indicate residue contacts observed in the crystal structures. Residue numbers for VHL are below 1000, and for SMARCA2 are above 1000. Residues located in the unstructured loop regions are labeled in magenta. B) The total number of contacts formed between residues in the SMARCA2 and VHL domains in the crystal structure and MD simulations. D) The residue pairs with more than 50% contact frequency in P1 (C) and P5 (D) are shown as yellow dashed lines. SMARCA2 is colored in blue, VHL is colored in grey, the loop regions of the SMARCA2-VHL interface are colored in magenta.

### High cooperativity PROTACs show a greater number of frustrated residue contacts in the protein-protein interface of the ternary complex

As described in the previous section, part of the SMARCA2-VHL interface interactions involves disordered loops in SMARCA2, ranging from residues N1396 to F1431 and from L1465 to S1468. Residues in unstructured loops contribute to the SMARCA2-VHL interface contacts during MD simulations (Fig. 5A). As discussed by Marton et al. ^30^, disordered regions sample a wider range of conformations and thus play an important role in protein function. Chen et al. ^12^ have shown that functionally critical structural regions in a protein often have patches of ‘frustrated’ residues that adopt suboptimal energetic states. Based on this concept, Gonzalo-Parra et al. developed the ‘Frustratometer’ toolkit^31^ to calculate the frustration of residue contacts within proteins or between interacting protein domains. The Frustratometer toolkit is based on the coarse-grained forcefield AWESEM^32^. Briefly, in the Frustratometer procedure, each amino acid pair is mutated *in silico* to all other amino acids and the energy distribution for all the mutated pairs (decoy pairs) is calculated with respect to the energy of the wild type pair. The wildtype residue pair energy is then converted into a z-score (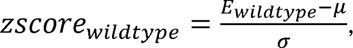 where μ and σ are the mean and standard deviation of the decoy energy distribution, respectively). The z-score is a measure of the wildtype contact energy relative to all other amino acid substitutions at these residue positions, and hence an estimation as to whether a residue is in an energetic optimal or suboptimal state. Each residue pair was classified as highly frustrated, neutral, or minimally frustrated based on the criteria employed in Frustratometer V2 (a z-score of mutational frustration > 0.78 is minimally frustrated, < -1 is highly frustrated and between 0.78 and -1 is neutral). More details on the calculation of frustration are given in the Methods section.

For each PROTAC ternary complex shown in Fig. 2, we used the Frustratometer^31^ on the MD trajectories and calculated the number of frustrated residue pairs in the SMARCA2-VHL interface for each MD snapshot. As shown in Fig. 6A, the ratio of the average number of frustrated residue pairs to the total number of contacts (between SMARCA2-VHL and between VHL or SMARCA2-PROTAC) in the interface, for each PROTAC TC correlates with measured cooperativity. Since frustrated residue pairs are often located at active sites and protein interfaces^12^, we speculate that the frustrated interface residues could, could show a high energy transient state that is indicative of catalytically active conformations and hence PROTAC cooperativity.

**Figure 6:**
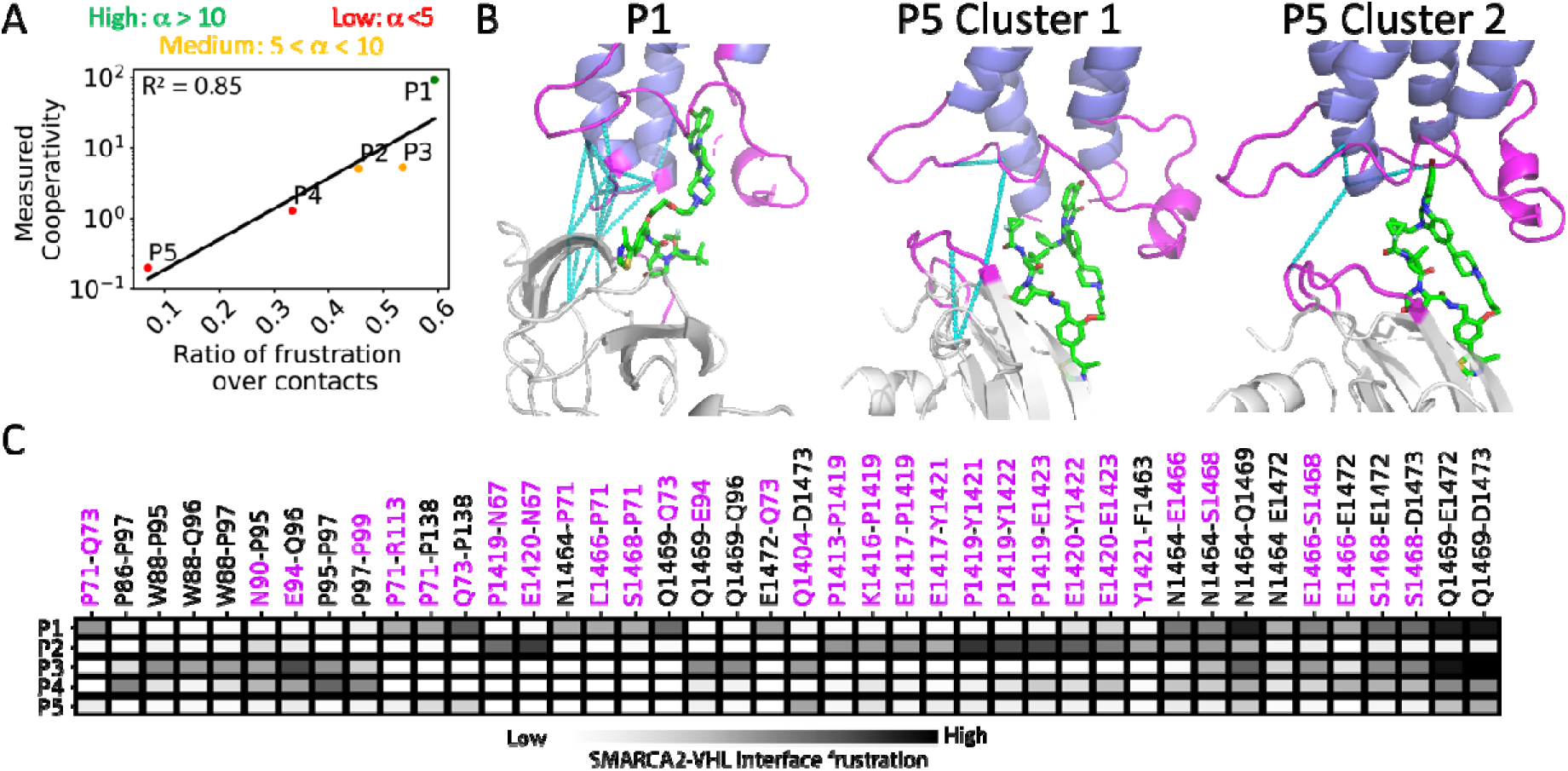
Frustration in SMARCA2-VHL-PROTAC TC. A) The ratio of the average number of highly frustrated residue pairs in SMARCA2-VHL interface and the average number of SMACAR2-VHL contacts plus VHL-PROTAC or SMARCA2-PROTAC contacts, compared to the cooperativity value. PROTACs with cooperativity greater than 10 are colored in green, between 5 and 10 are colored as orange, below 5 are colored as red. B) The highly frustrated residue pairs from the most populated cluster of PROTAC P1 and the top 2 populated clusters of P5 are shown as cyan dash lines. SMARCA2 is colored in blue, VHL in white. The unstructured loop region of SMARCA2-VHL interface is colored in magenta. C) Heatmap of frustration level for residue pairs located in the SMARCA2-VHL interface. The frustration level was calculated by averaging the frustration scores of all frames from the MD simulation. Residues located in the unstructured loop region are labeled in magenta.

Most frustrated residues are in the loop regions of either VHL or SMARCA2 as shown in Figs. 6B for high (P1) and low (P5) cooperativity PROTACs. This is also shown in the frustration heat map for the SMARCA2-VHL interface residue pairs (Fig. 6C). Movies S3 and S4 showing the three-dimensional view of this region are given in the Supporting Information. Using double mutant cycles, Jemth et al. ^33^ have shown that the interactions in a protein complex involving unstructured domains, especially in promiscuous protein complexes, are likely to be energetically suboptimal and exhibit some strain due to unfavorable interactions. This is what we observe here in the PROTAC ternary complexes. In most MD simulation runs, the number of frustrated residues in the SMARCA2 protein, is higher than in VHL (Fig. S4). Interestingly, Proline is present in 23% of highly frustrated pairs in all five simulations. Both glutamine and glutamic acid are present in 21% of highly frustrated pairs, and asparagine in 11% of the frustrated residue pairs in the interface. These frequently occurring residues imply that the chemical nature of amino acid may also play a role in regulating the frustration level of PPIs, and it has been shown in literature that certain residues may be more critical in forming a catalytically effective interface ^30,34^. In summary, we show that, for the SMARCA2/VHL system under study, the frustrated residue pairs in the PPI of PROTAC ternary complexes can successfully distinguish high from low cooperativity PROTACs, and we will use the same high/low terminology in future discussion. It is the focus of future research to establish as to whether this can be utilized for all POI-E3 pairings in which cooperativity is a relevant optimization parameter.

### Frustration involving the POI-E3 interface residues can be used to assess the cooperativity of PROTACs prior to synthesis

One goal of this work was to assess the utility of the frustration in the PPI for predicting the cooperativity of PROTACs, and ideally to assist synthesis prioritization. As a further validation of the frustration measure, we studied 11 GEN-1 based PROTACs (shown in Fig. S1) that have no crystal structures except P13 (pdb ID: 6HAY). Using the methods detailed in the Methods section, we generated ternary complex structural models using a now publicly available PROTAC (P2) ternary complex template structure (pdb ID: 6HAX)^8^. We speculated that TC conformations, present in high-cooperativity PROTACs such as the one in 6HAX structure, would lead to more pronounced differences in the number of frustrated residue pairs. The 11 PROTACs feature the same VHL binder as the PROTACs P1, P2, P4 and P5, except for P19 and P20, which contain a non-binding distomer of VH101. The SMARCA2 binder in these PROTACs, GEN-1, binds into the same pocket and provides a similar exit vector.^21^ Starting from the structural models of the 11 PROTAC TCs, we performed MD simulations using the sam protocol as applied to the crystal structures (see Methods). Using the MD simulation trajectories for each TC model, we calculated the average number of frustrated residue pairs located at the interface between SMARCA2 and VHL. To strengthen the error estimation, we increased the MD simulation sample size by generating 10 different conformation ensembles using bootstrapping methods (see Methods for more details). As shown in Fig. 7, the average number of frustrated residue pairs shows the same trend as the cooperativity, with the higher cooperativity PROTACs being more likely to show higher frustration than weak cooperativity PROTACs. Although there are outliers, the trend from the docked models concurs with the frustration pattern observed among the high, medium and low cooperativity PROTACs in the MD simulations starting from the crystal structures. This indicates that frustration can be used to predict cooperativity and hence to prioritize PROTACs for synthesis and experimental testing.

**Figure 7:**
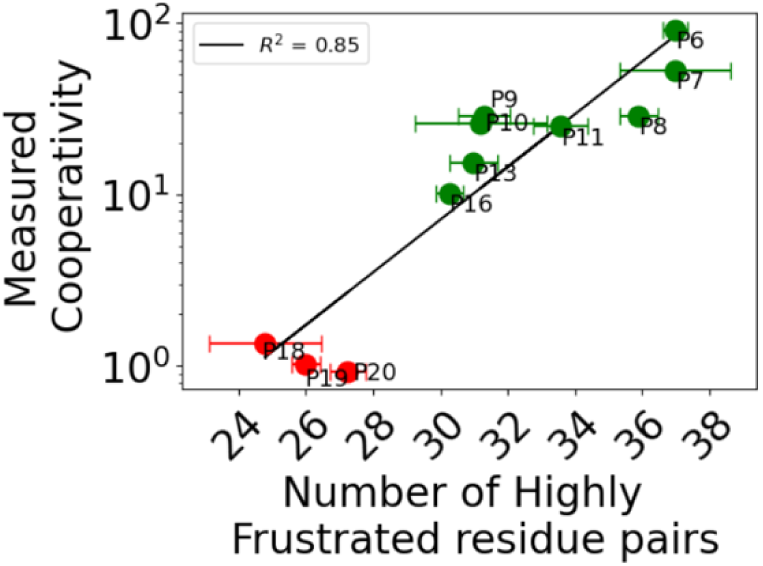
The average number of highly frustrated pairs (along with their standard deviations shown as line) from MD simulations starting from the docked models of 11 PROTACs are shown along with their cooperativity (shown as dot). PROTACs with cooperativity greater than 10 are colored in green, between 5 and 10 are colored orange, below 5 are colored red.

## Discussion

Despite some notable successes, the design of PROTACs can be challenging, since conventional structure-based drug design principles using single crystal structures do not represent the highly dynamic nature of some ternary complexes formed with PROTACs. PROTAC properties are governed by the conformational ensemble and the interacting interfaces in the ternary complexes. Comparison of the five SMARCA2::PROTAC::VHL crystal structures of these TCs reveals large domain motions. We derived the free energy surfaces of these from MD simulations, and identified multiple free energy minima, representing distinct TC conformations. We also observed that the TCs formed with low-cooperativity PROTACs are more flexible than those with high-cooperativity PROTACs. The spread of the relative orientations in the SMARCA2 domain is wider in the MD simulations than observed when comparing crystal structures only. The orientation of the E3 ligase domains relative to the POI have been shown to play a critical role in substrate selectivity by PROTACs ^35^. Such dynamic behavior of TCs with other POIs or E3 ligases, such as cereblon, have been shown by other MD simulation studies^17,36^. We observed that the number of residue contacts formed during MD simulations in the protein-protein interacting interface is larger in high-cooperativity PROTACs when compared to low-cooperativity PROTACs (Fig S4 and Fig 5B). Such differences have also been seen in cereblon-based PROTACs paired with other POIs ^27^.

Four new crystal structures along with the MD simulations also showed that, for different PROTACs, the SMARCA2-VHL interface is formed by residues that are mostly located in the unstructured regions of the two proteins. Frustration is a measure of the degree of suboptimal energy states of protein residues, which are often found in the unstructured regions of a protein. Frustration in protein-protein interfaces arises when residues at the interface cannot adopt optimal conformations or achieve dense packing due to conflicting interactions ^29^. While it is generally understood that frustration decreases the stability of a protein, the relationship between frustration and the functional activity of a protein remains vague. It is known that in some enzymes, frustration in the active site can promote catalysis by destabilizing the reactants, products and also facilitate the transition state^37^. Here we observe that the number of highly frustrated residue pairs in the SMARCA2-VHL interface of the ternary complex is higher for high-cooperativity than for low-cooperativity PROTACs. The frustrated residue pairs have a higher occurrence of prolines, glutamine and glutamic acid residues compared to other amino acid residues. This is consistent with the finding that these residues are also more frequently found in intrinsically disordered regions of proteins ^34^. Therefore, we hypothesize that frustration in the POI-E3 interface can be utilized for characterizing the nature of the TCs and for describing cooperativity. We have provided initial evidence that the number of frustrated residue pairs can be used as a measure to distinguish high from low cooperativity PROTACs, based on the TC models that were generated by comparative modelling in combination with MD simulations. In the future, we aim to use the specific frustrated residue pairs in the PPI of TCs for rational design of PROTACs..

## Methods

### MD simulation protocol

The VHL::P2::SMARCA2^BD^ ternary complex crystal structure has previously been published and is available in the protein databank (PDB ID: 7Z77)^9^. Each PROTAC ternary complex was initially prepared using the Maestro Protein Preparation Wizard module^38^ (Schrödinger Release 2023-1: Maestro, Schrödinger, LLC, New York, NY, 2021). *H*istidine protonation states were determined under neutral pH conditions. PROTAC forcefield parameters were derived using the Antechamber module from AmberTools^24^, while the partial charges were calculated using the Jaguar module in Maestro^39^. The protein atoms were parameterized using the FF99SB-ILDN force field^40^. The PROTAC TC complex was solvated in a truncated octahedron SPC^41^ water box with at least 14 Å margin from the protein surface using tleap from the AmberTools 18 package^24^. The simulation box was neutralized using adequate number of Na^+^/Cl^-^ ions. The simulation box went through a two-step minimization procedure. In the first step, a 2000-step steepest descent minimization was conducted, with a 10 kcal/mol constraint applied to all atoms, excluding water. During the second minimization phase, all atoms were permitted to move, and an additional 2000-step steepest descent minimization was carried out. Following minimization, an equilibration simulation was executed. The initial equilibration step involved a brief 0.5 ns NVT heating phase, raising the temperature from 0K to 300K with a 10 kcal/mol restraint applied to the protein and PROTAC’s heavy atoms. The system underwent a seven-step NPT equilibration to gradually release restraints at 1 atmosphere pressure and 300K temperature. Restraints were gradually reduced from 10 kcal/mol to 5 kcal/mol, then 4, 3, 2, 1 with each step lasting for 5 ns. Finally, a 10 ns equilibration without restraints was conducted at the end. The final snapshot of the equilibration phase was used as the initial conformation for production runs. Six random velocities were employed to generate six independent production runs. Each production run lasted for 500 ns, and the complete production trajectories (30,000 snapshots in total) were used for subsequent analysis. Throughout the simulations, the temperature and pressure were controlled by the Nosé-Hoover method ^42,43^. The particle mesh Ewald method was utilized to compute the electrostatic interactions^44^, while the non-bound interactions were determined using a 12 Å cutoff.

### Principal Component Analysis and Shannon entropy calculation

We conducted principal component analysis (PCA) on the combined production trajectories, considering all Cα atom positions for the RMSD calculation in the PCA, and projected the resulting principal components (PCs). The first two principal components constituted approximately 80% of the total eigenvector variance (Figure S1). The gmx anaeig module^45^ was employed to examine the protein motions represented by the first two principal components. The two TC conformations with maximal PC1 and PC2 values were aligned onto the VHL domain to assess the conformational changes in the SMARCA2 domain (Figure 2A). The combined sampling points were projected onto the first two PC axes and displayed as a contour map (Figure 2B). The individual contour map for each PROTAC is displayed in Figure S2. The combined structural ensemble of all five PROTACs was clustered in the top two PC dimensions using the agglomerative clustering algorithm, generating seven clusters (yielded the best Silhouette Coefficient). The conformations corresponding to the centroids of all seven clusters were aligned by their VHL domains to compare the orientations of the SMARCA2 domains (Figure 2C). For each PROTAC, we calculated the Shannon entropy as -∑ P_i_ ln (P_i_), where *P_i_* denotes the fraction of conformations that fall into cluster i (Figure 2D).

### RMSD based clustering and contact analysis

Each TC ensemble was clustered based on the RMSD among the Cα atom positions using the program CPPTRAJ ^46^. We selected a cutoff between 1.5 to 2.5 Å such that the top three clusters accounted for approximately 60% of the overall population. The representative structures of the top three populated clusters were extracted and aligned onto their VHL domain to display the conformational differences in the SMARCA2 domain (Figure 3A). The RMSD values among the top three representative structures were calculated as follows: P1: 0.5∼0.8 Å, P2: 0.5∼0.8 Å, P3: 0.5∼0.8 Å, P4: 1.4∼4.5 Å, P5: 0.4∼2.7 Å. We computed the contact frequencies between the proteins and the PROTAC, as well as between SMARCA2 and VHL, for both the crystal structures and the simulation trajectories. The contact analysis was carried out using the program get_contact (https://getcontacts.github.io/), considering every atom and all types of contacts in the interface^16,34,35^.

### Frustration calculation using the MD conformations

Following the seven clusters obtained from the prior PCA, we conducted frustration calculations for each cluster from every PROTAC, using Frustratometer^31^. For each cluster, we initially identified the SMARCA2-VHL interface residue pairs from every snapshot by locating all pairs with a minimum distance of less than 4.5 Å between heavy atoms. We then calculated the frustration levels for these identified pairs in each MD snapshot. Each pair was classified as highly frustrated, neutral, or minimally frustrated based on the criteria employed in Frustratometer V2 (a z-score of mutational frustration > 0.78 is minimally frustrated, < -1 is highly frustrated). For each cluster, we counted the number of residue pairs that were highly frustrated in over 40% of the cluster population. To compare the level of frustration among different PROTAC simulations, we calculated the average number of highly frustrated residue pairs over all seven clusters, weighted by the cluster populations.

### Comparative modeling, simulation, analysis of PROTAC ternary complexes without crystal structures

To investigate the 11 GEN-1 based PROTACs, we utilized the publicly available SMARCA2^BD^: PROTAC 2: VHL crystal structure (PDB ID: 6HAX)^8^ as template to construct ternary complex models, as 6HAX displays a similar SMARCA2-VHL conformation to that found in the P1 and P2 TCs. This was one of the two crystal structures of SMARCA2-VHL complexes available at the start of this project. The purpose of generating these ternary structural models was not to predict the complex structure that each of the 11 PROTACs would form. Instead, our goal was to place each of the 11 PROTACs in a known high cooperativity PROTAC ternary model conformation and perform short MD simulations to calculate the frustration in the protein-protein interface.

We constructed the 3D coordinates of each of the 11 PROTACs based on the P2 in the 6HAX conformation as reference using Maestro and these were subjected to geometry optimization using Hartree-Fock (HF) quantum mechanical calculations and the STO-3G basis set as implemented in the Jaguar module of Maestro^39^. Then these structures were aligned with PROTAC 2 in the 6HAX ternary complex formed by SMARCA2 and VHL using the pair-fit utility in PyMOL 3.0^47–49^. Subsequently, the local clashes between the protein side chain atoms and the newly inserted PROTACs were resolved using the side chain replacement feature in Maestro’s PRIME module^50,51^. Partial charges calculated from the HF calculations were used in the molecular topology of PROTACs created by Antechamber for MD simulations without any modifications.

Each TC was prepared and simulated using the same protocol as described in the previous sections. For each system, we conducted three 100 ns-long simulations starting with random velocities. After the production runs, we performed RMSD clustering and extracted frames with a SMARCA2-VHL interface RMSD below 1.5 Å with the initial structure, for a frustration calculation using Frustratometer. This step is to ensure that the frustration metric can be compared across all 11 PROTACs sharing a consistent ensemble of PPIs.

To account for sampling variability in the frustration calculations, we implemented a bootstrapping approach. For each PROTAC, from the subset of conformations with RMSD < 1.5Å, we randomly selected 100 frames, repeating this process 10 times. The number of highly frustrated pairs was calculated from each of the 10 trials, and the average and standard deviation were estimated to assess the level of robustness in our sampling protocol.

### Chemical synthesis

A list of final compounds is compiled in Supplementary Table 2 and Supplementary Fig. 4. Full details of synthetic procedures for P1, P3, P4, P9, P10, P11, and P18 (including supplementary schemes S1-S9) with NMR and HPLC spectra of final compounds are provided as Supplementary Methods. The syntheses of P2 and P5 have been described in reference^9^. (compounds 6 and 23 therein respectively), the syntheses of P6, P7, and P16 have been described in reference^52^. the syntheses of P8, P13, P19, and P20 have been described in reference^8^ (ACBI1, PROTAC 1, *cis*-ACBI1 and *cis*-PROTAC 2 therein respectively).

### Protein production and crystallography

Protein production for SMARCA2^BD^ and the VCB complex was done as previously described^8,9^. For P1 and P3 ternary complex crystallization, SMARCA2^BD^ and VCB complex were incubated with 1 mM of the ligand in a 1:1:1 ratio overnight and crystallized in 96-well sitting drop plates. P1 (BI01802957) was crystallized with a reservoir solution of 0.1 M K_2_HPO_4_, 16% PEG 8000 and 0.2 M sodium chloride. For P3 (BI01802953), the reservoir solution was 0.1 M Bis/Tris propane pH 7.5, 32% PEG 3350 and 0.1 M tri-sodium citrate. Datasets were measured at SLS beamline X10SA and processed with autoPROC^53^. Resolution cutoffs were determined using the default parameters of autoPROC. The structures were solved by molecular replacement using Phaser^54^ with PDB: 7Z76^9^ as a search model. The final model was prepared in iterative cycles of model building with COOT^55^ and refinement with autoBUSTER (Global Phasing Ltd). Data collection and refinement statistics are provided in Supplementary table 3 and Supplementary table 4, respectively. For the P4 (BI01701345) ternary complex: SMARCA2^BD^, VCB complex and P4 were mixed in a 1:1:1 molar ratio, concentrated to 10 mg/mL in a buffer containing 0.1 M HEPES pH 7.5, 100 mM sodium chloride, 1 mM TCEP, 2% DMSO, incubated at room temperature. Drops were prepared by mixing 1 μL of the ternary complex with 1 μL of well solution and crystallized at 18 °C using the hanging-drop vapor diffusion method. Crystals were obtained in 0.1 M HEPES pH 7.0, 8% ethylene glycol, 14% PEG 8K. Harvested crystals were flash cooled in liquid nitrogen following gradual equilibration into cryoprotectant solution consisting of 20% PEG 8K, 0.1 M HEPES pH 7.0, 20% ethylene glycol. Diffraction data were collected at Diamond Light Source beamline I24 (λ = 0.9686 A) using a Pilatus3 6 M detector and processed using XDS. The crystals belonged to space group P21 with unit cell parameters a = 48.7, b = 89.8, c = 64.2 A and α = 90.0°, β = 96.9°, γ = 90.0° and contained one copy of the ternary complex per asymmetric unit. The structure was solved by molecular replacement using PHASER with VCB coordinates derived from the VCB: MZ1: Brd4^BD2^ complex (PDB: 5T35) and SMARCA2^BD^ (PDB: 4QY4) as search models. Subsequent iterative model building, and refinement, was done according to standard protocols using CCP4, COOT, and autoBUSTER (Global Phasing Ltd). For the VCB:P5(BI01702722):SMARCA2^BD^ ternary complex: VCB, P5 and SMARCA2^BD^ were mixed in a 1:1:1 stoichiometric ratio in 20 mM HEPES, pH 7.5, 100 mM sodium chloride, 1 mM TCEP, 2% DMSO, incubated at room temperature and concentrated to a final concentration of approximately 16 mg/ml. Drops were prepared by mixing 1 μL of the ternary complex with 1 μL of well solution and crystallized at 18 °C using the hanging-drop vapor diffusion method. Crystals were obtained in 28% PEG 3350, 0.2 M potassium thiocyanate, 0.1 M Bis-Tris Propane pH 7.5. Harvested crystals were flash cooled in liquid nitrogen following gradual equilibration into cryoprotectant solution consisting of 40% PEG 3350, 0.2 M potassium thiocyanate, 0.1 M Bis-Tris Propane pH 7.5. Diffraction data were collected at Diamond Light Source beamline I24 (λ = 0.9686 A) using a Pilatus3 6 M detector and processed using XDS. The crystals belonged to space group P21 with unit cell parameters a = 46.6, b = 81.6, c = 58.1 A and α = 90°, β = 97.8°, γ = 90.0° and contained one copy of the ternary complex per asymmetric unit. The structure was solved by molecular replacement using PHASER with VCB coordinates derived from the VCB: MZ1: Brd4^BD2^ complex (PDB: 5T35) and SMARCA2^BD^ (PDB: 4QY4) as search models. Subsequent iterative model building, and refinement, was done according to standard protocols using CCP4, COOT, and autoBUSTER (Global Phasing Ltd).

## Abbreviations

PROTAC: **Pro**teolysis **Ta**rgeting **C**himeras
POI: Protein of interest
E3: E3 ubiquitin ligase
MD: Molecular Dynamics
RMSD: root mean square deviation
PC: Principal Component
PCA: Principal Component Analysis
α: Cooperativity;
TC: ternary complex (formed by POI, E3 and PROTAC)
VHL: von Hippel-Lindau disease tumor suppressor
BD: bromo domain
SMARCA: SWI/SNF related BAF chromatin remodeling complex subunit ATPase
PPI: protein-protein interface
VCB: VHL-Elongin BC-CUL2

## Data availability

Coordinates and structure factors for the P1 (BI01802957), P3 (BI01802953), P4 (BI01701345), and P5 (BI01702722) ternary complexes with SMARCA2^BD^ and the VCB complex have been deposited at the PROTEIN DATA BANK with accession codes XXXX, YYYY, ZZZZ, and AAAA, respectively.

## Acknowledgements

The research conducted in the Ciulli laboratory reported in this study has received funding from Boehringer Ingelheim. Biophysics and drug discovery activities at Dundee were supported by Wellcome Trust strategic awards to Dundee (100476/Z/12/Z and 094090/Z/10/Z, respectively). We thank Karen J. Bergner for her invaluable help in carefully revising the manuscript.

## Competing interests

A.C. is a scientific founder and shareholder of Amphista Therapeutics, a company that is developing targeted protein degradation therapeutic platforms. The Ciulli laboratory receives or has received sponsored research support from Almirall, Amgen, Amphista Therapeutics, Boehringer Ingelheim, Eisai, Merck KaaG, Nurix Therapeutics, Ono Pharmaceutical, and Tocris-Biotechne. Z.J., P.S., C.K.,P.G.,H.W.,G.B. and A.B. are current employees of Boehringer Ingelheim. E.D. is now an employee of Astra Zeneca. All other authors declare that they have no competing interests.

## Supporting Information

### Supplementary Figure

**Figure S1.**
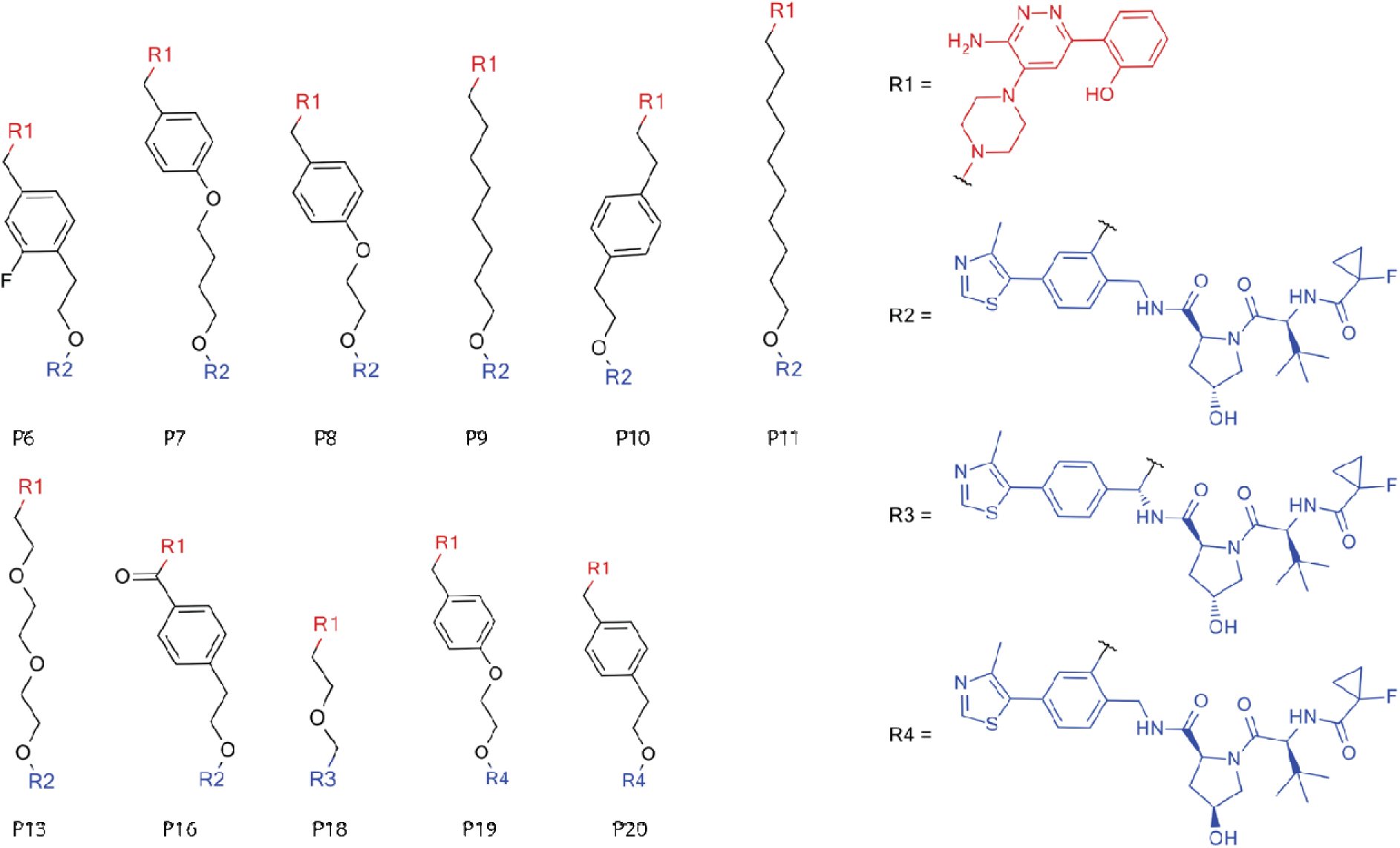
Eleven GEN-1 based PROTACs investigated in this work. P9, P10, P11 and P18 have not been described previously, their synthesis is detailed below.

**Figure S2:**
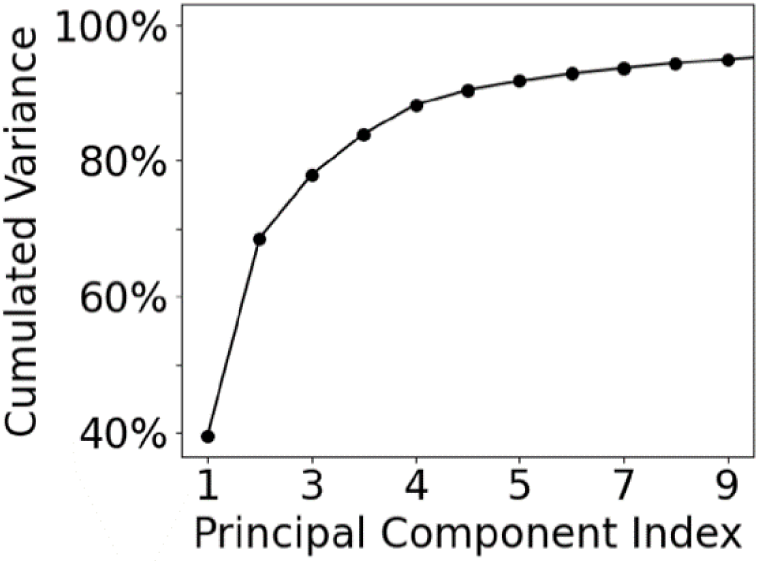
The cumulated variance in the Principal Components for the top 10 components. The Principal Component Analysis was performed on the aggregated trajectories of the MD simulations starting from the five crystal structures. This shows that the top 3 PCs recapitulate about 80% of the domain motion of the ternary complexes.

**Figure S3.**
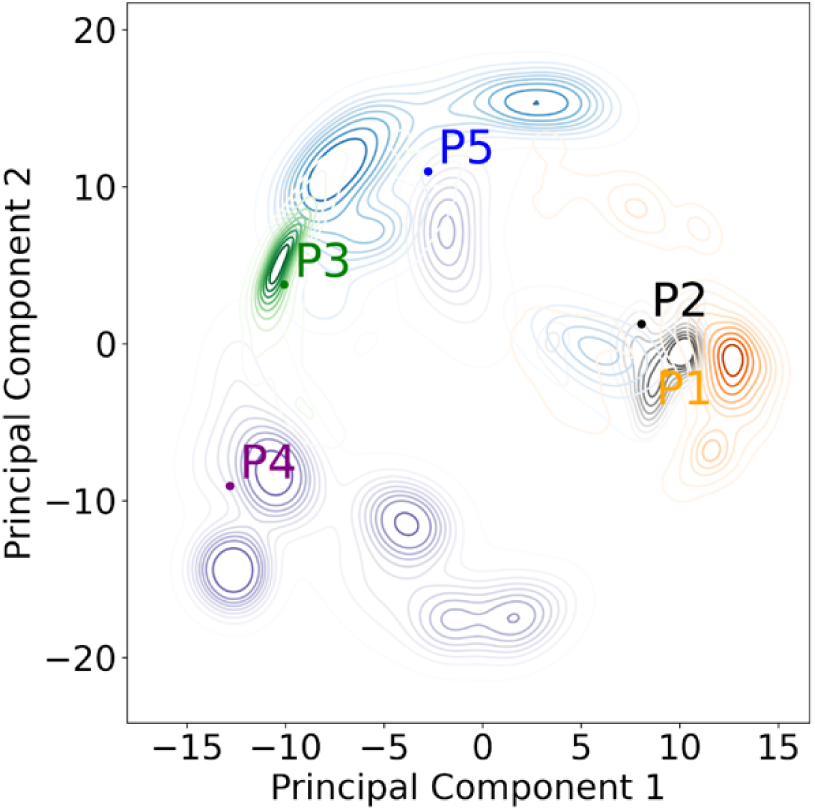
Individual contour maps for all five PROTACs, projecting onto a common PC landscape.

**Figure S4.**
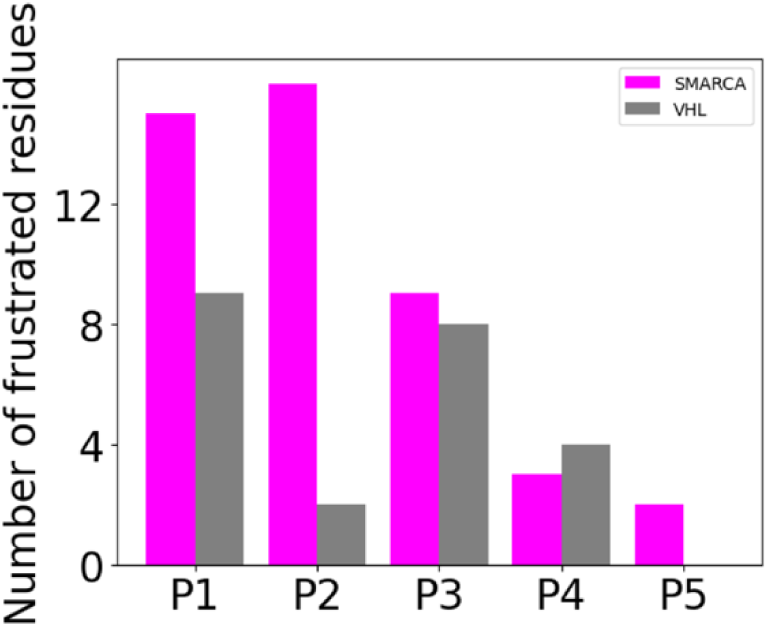
Number of SMARCA2 or VHL residues involved in the frustrated pairs located in the SMARCA2-VHL interface. SMARCA2 has greater number of frustrated pairs than VHL in all PROTACs except P4.

**Figure S5:**
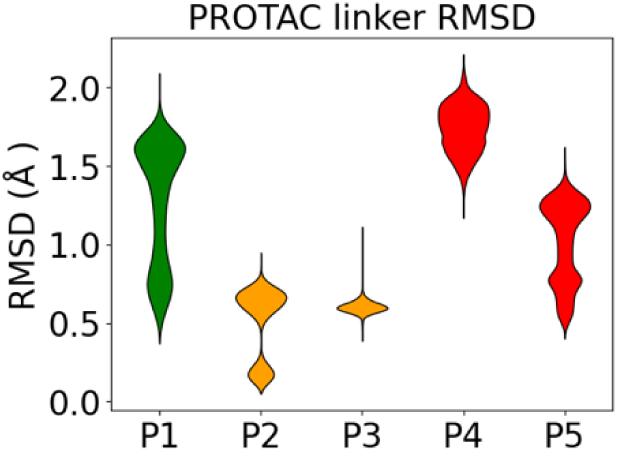
the RMSD of linkers in five PROTACs

**Table S1:**
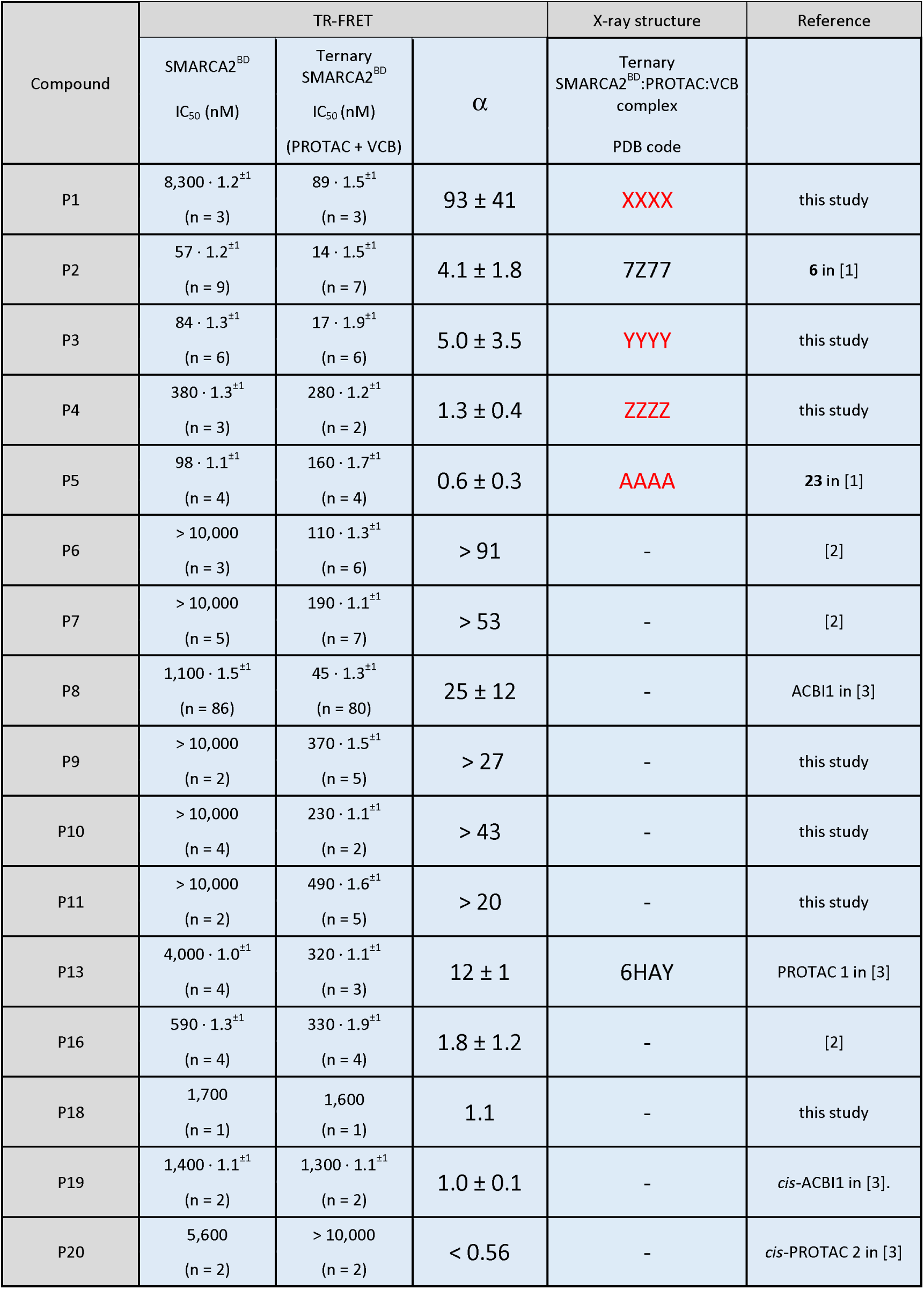
Measurement of PROTAC cooperativity (α) and ternary complex affinity (ternary IC_50_) via TR-FRET competition for SMARCA2^BD^ ± VCB. Experimental details are given in [1]. PROTAC cooperativity (Cp) is calculated as ratio between TR-FRET IC_50_ / TR-FRET IC_50_ +VCB values. IC_50_ data represent geometric means with errors stated as geometric standard deviation and number of experimental replicates (n) given in parentheses. Cooperativity uncertainties were calculated from IC_50_ geometric standard deviations using standard error propagation rules.

| Compound | TR-FRET |  |  | X-ray structure | Reference |
| --- | --- | --- | --- | --- | --- |
| | SMARCA2<sup>BD</sup><br>IC<sub>50</sub> (nM) | Ternary<br>SMARCA2<sup>BD</sup><br>IC<sub>50</sub> (nM)<br>(PROTAC + VCB) | $\alpha$ | Ternary<br>SMARCA2<sup>BD</sup>:PROTAC:VCB<br>complex<br>PDB code | |
| P1 | 8,300 · 1.2<sup>±1</sup><br>(n = 3) | 89 · 1.5<sup>±1</sup><br>(n = 3) | 93 ± 41 | XXXX | this study |
| P2 | 57 · 1.2<sup>±1</sup><br>(n = 9) | 14 · 1.5<sup>±1</sup><br>(n = 7) | 4.1 ± 1.8 | 7Z77 | 6 in [1] |
| P3 | 84 · 1.3<sup>±1</sup><br>(n = 6) | 17 · 1.9<sup>±1</sup><br>(n = 6) | 5.0 ± 3.5 | YYYY | this study |
| P4 | 380 · 1.3<sup>±1</sup><br>(n = 3) | 280 · 1.2<sup>±1</sup><br>(n = 2) | 1.3 ± 0.4 | ZZZZ | this study |
| P5 | 98 · 1.1<sup>±1</sup><br>(n = 4) | 160 · 1.7<sup>±1</sup><br>(n = 4) | 0.6 ± 0.3 | AAAA | 23 in [1] |
| P6 | > 10,000<br>(n = 3) | 110 · 1.3<sup>±1</sup><br>(n = 6) | > 91 | - | [2] |
| P7 | > 10,000<br>(n = 5) | 190 · 1.1<sup>±1</sup><br>(n = 7) | > 53 | - | [2] |
| P8 | 1,100 · 1.5<sup>±1</sup><br>(n = 86) | 45 · 1.3<sup>±1</sup><br>(n = 80) | 25 ± 12 | - | ACBI1 in [3] |
| P9 | > 10,000<br>(n = 2) | 370 · 1.5<sup>±1</sup><br>(n = 5) | > 27 | - | this study |
| P10 | > 10,000<br>(n = 4) | 230 · 1.1<sup>±1</sup><br>(n = 2) | > 43 | - | this study |
| P11 | > 10,000<br>(n = 2) | 490 · 1.6<sup>±1</sup><br>(n = 5) | > 20 | - | this study |
| P13 | 4,000 · 1.0<sup>±1</sup><br>(n = 4) | 320 · 1.1<sup>±1</sup><br>(n = 3) | 12 ± 1 | 6HAY | PROTAC 1 in [3] |
| P16 | 590 · 1.3<sup>±1</sup><br>(n = 4) | 330 · 1.9<sup>±1</sup><br>(n = 4) | 1.8 ± 1.2 | - | [2] |
| P18 | 1,700<br>(n = 1) | 1,600<br>(n = 1) | 1.1 | - | this study |
| P19 | 1,400 · 1.1<sup>±1</sup><br>(n = 2) | 1,300 · 1.1<sup>±1</sup><br>(n = 2) | 1.0 ± 0.1 | - | cis-ACBI1 in [3]. |
| P20 | 5,600<br>(n = 2) | > 10,000<br>(n = 2) | < 0.56 | - | cis-PROTAC 2 in [3] |

**Table S2:** The population percentage of each cluster for all the five PROTAC simulations. These population percentages were used to calculate the Shannon entropy for each PROTAC.

|  | P1 | P2 | P3 | P4 | P5 |
| --- | --- | --- | --- | --- | --- |
| Cluster 1 | 0% | 0% | 7% | 42% | 0% |
| Cluster 2 | 1% | 0% | 7% | 15% | 39% |
| Cluster 3 | 22% | 60% | 0% | 2% | 12% |
| Cluster 4 | 0% | 0% | 85% | 40% | 19% |
| Cluster 5 | 0% | 40% | 0% | 0% | 0% |
| Cluster 6 | 1% | 0% | 1% | 1% | 27% |
| Cluster 7 | 76% | 0% | 0% | 0% | 3% |

**Table S3:**
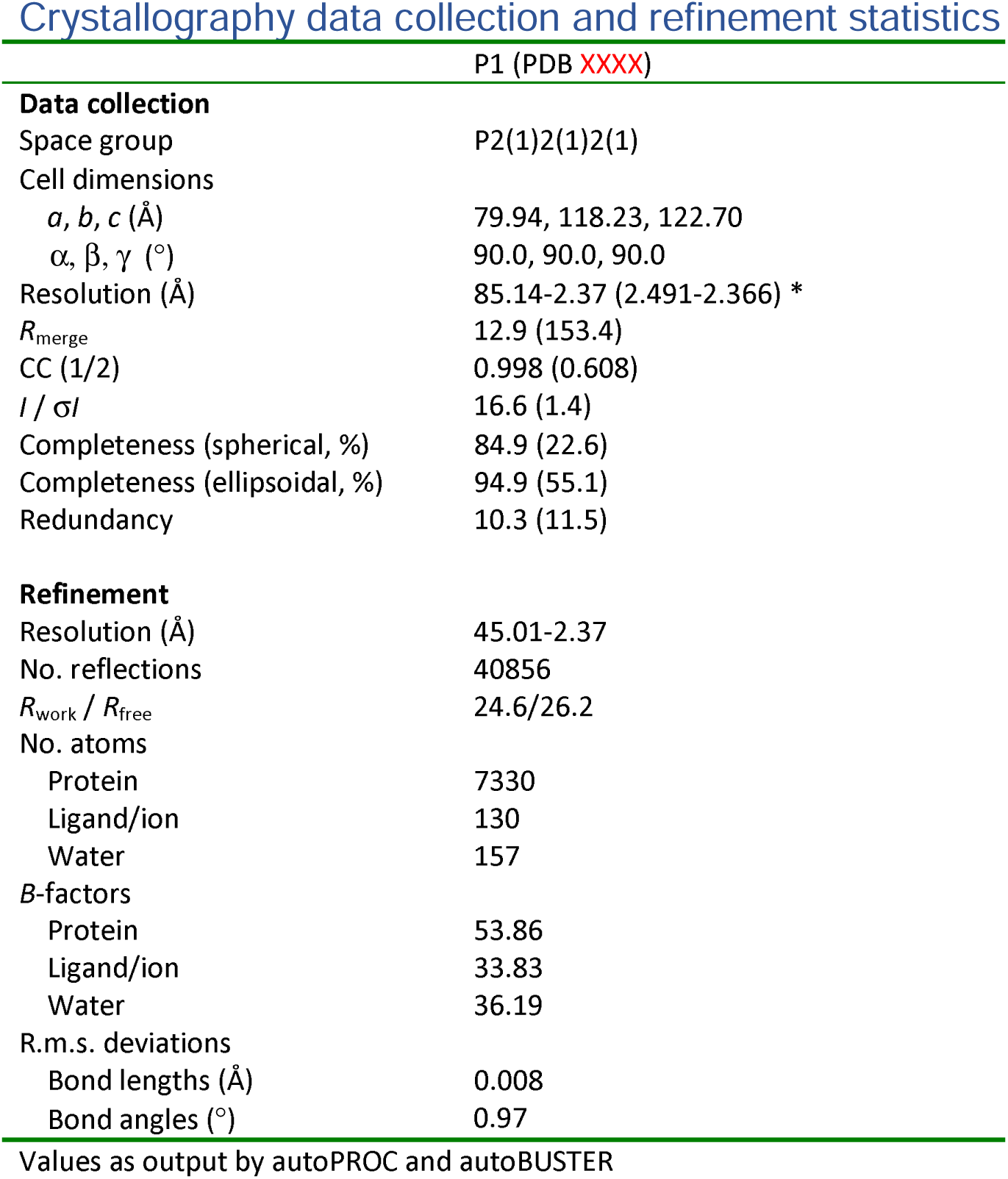
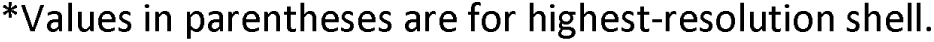
Data collection and refinement statistics for VCB : P1 : SMARCA2^BD^ ternary complex co-crystal structure

**Table S4:**
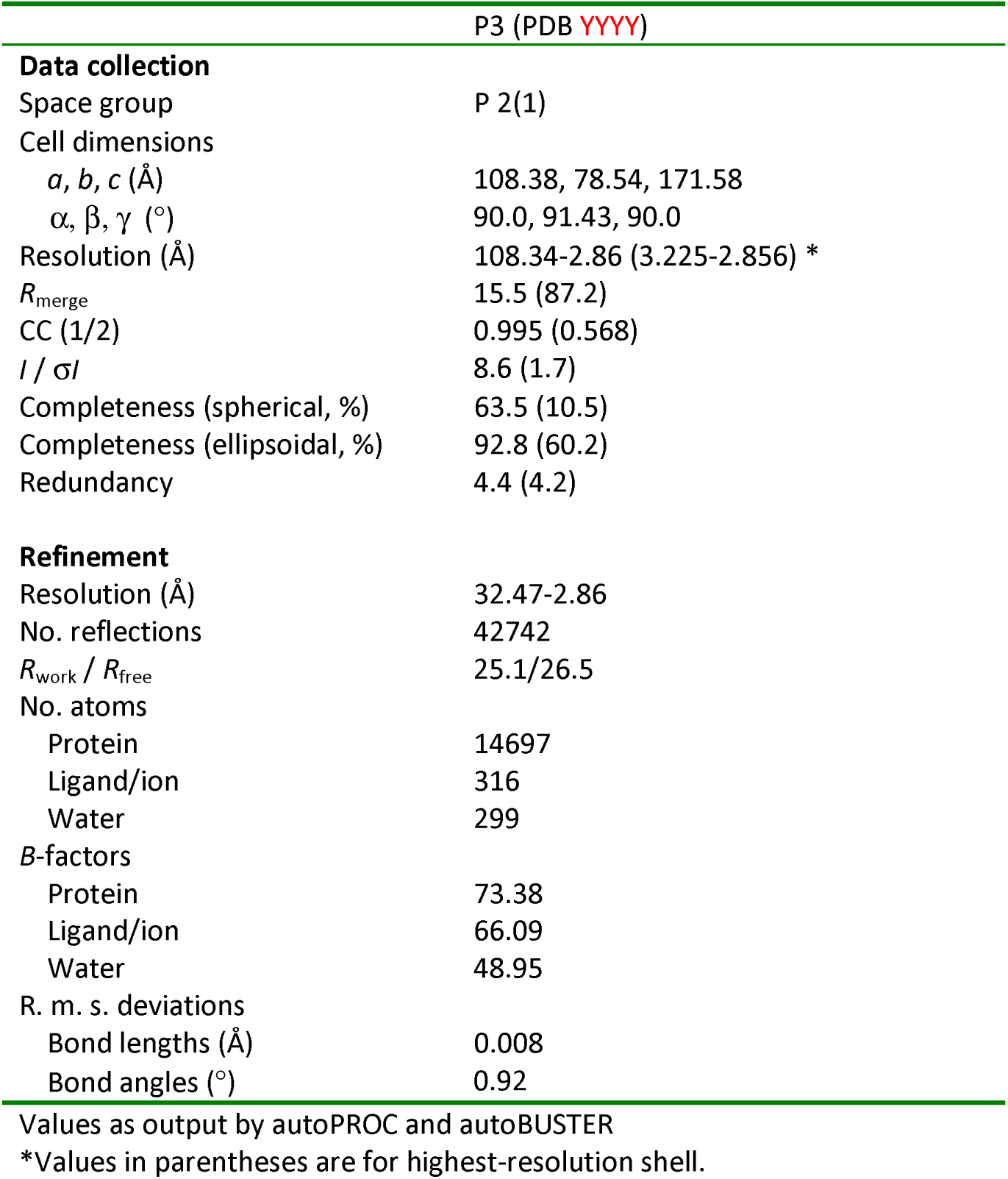
Data collection and refinement statistics for VCB : P3 : SMARCA2^BD^ ternary complex co-crystal structure

|  | P3 (PDB YYYY) |
| --- | --- |
| **Data collection** |  |
| Space group | P 2(1) |
| Cell dimensions |  |
| *a*, *b*, *c* (Å) | 108.38, 78.54, 171.58 |
| $\alpha$, $\beta$, $\gamma$ (°) | 90.0, 91.43, 90.0 |
| Resolution (Å) | 108.34-2.86 (3.225-2.856) * |
| *R*<sub>merge</sub> | 15.5 (87.2) |
| CC (1/2) | 0.995 (0.568) |
| *I* / σ*I* | 8.6 (1.7) |
| Completeness (spherical, %) | 63.5 (10.5) |
| Completeness (ellipsoidal, %) | 92.8 (60.2) |
| Redundancy | 4.4 (4.2) |
| **Refinement** |  |
| Resolution (Å) | 32.47-2.86 |
| No. reflections | 42742 |
| *R*<sub>work</sub> / *R*<sub>free</sub> | 25.1/26.5 |
| No. atoms |  |
| Protein | 14697 |
| Ligand/ion | 316 |
| Water | 299 |
| *B*-factors |  |
| Protein | 73.38 |
| Ligand/ion | 66.09 |
| Water | 48.95 |
| R. m. s. deviations |  |
| Bond lengths (Å) | 0.008 |
| Bond angles (°) | 0.92 |

Values as output by autoPROC and autoBUSTER
\*Values in parentheses are for highest-resolution shell.

**Table S5:**
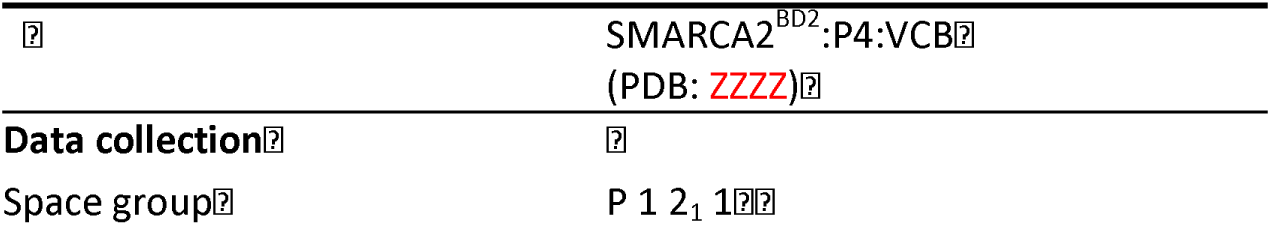
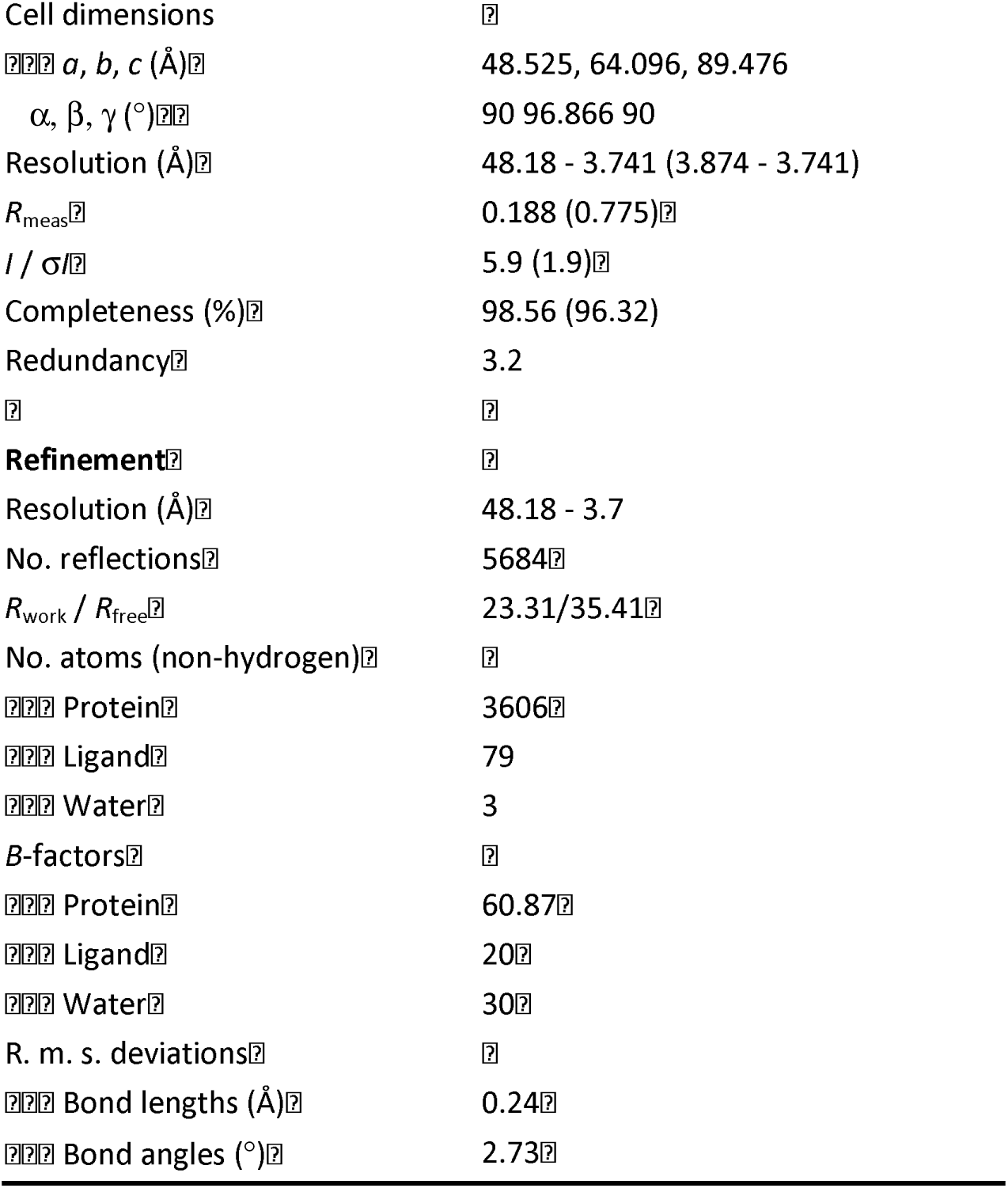
Data collection and refinement statistics for VCB : P4 : SMARCA2^BD^ ternary complex co-crystal structure

**Table S6:**
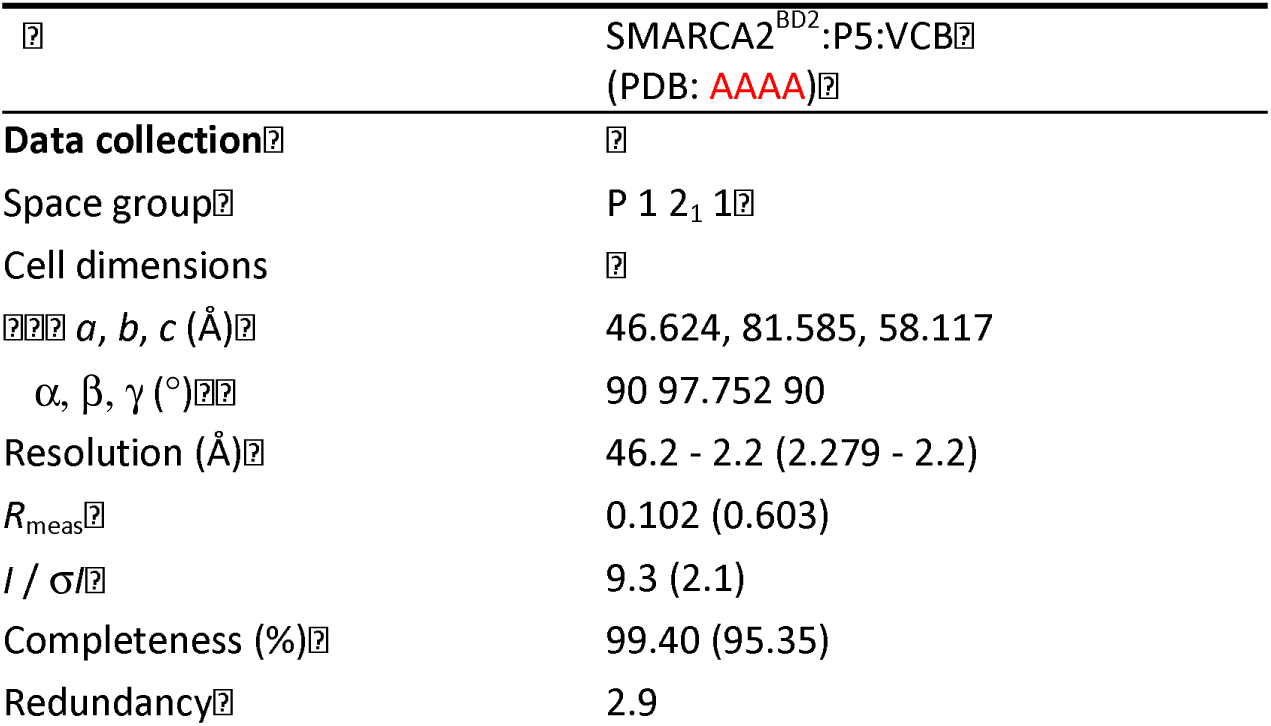
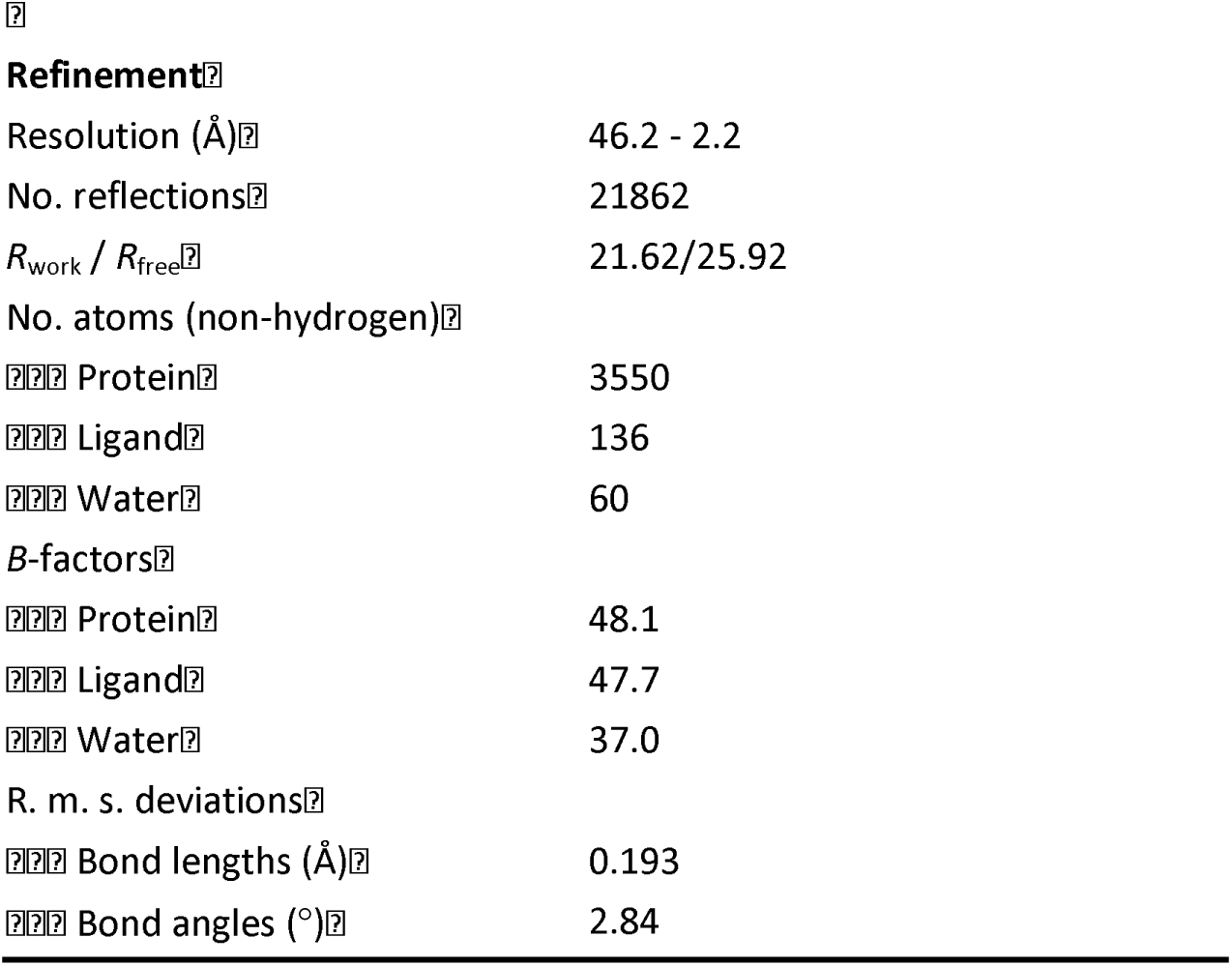
Data collection and refinement statistics for VCB : P5 : SMARCA2^BD^ ternary complex co-crystal structure

### Supplementary Methods

#### Analytical experiments

##### NMR

NMR experiments were recorded on Bruker Avance 500 MHz or 600 MHz spectrometer at 298 K. Samples were dissolved in 600 µL DMSO-d6 and TMS was added as an internal standard. 1D 1H spectra were acquired with 30° excitation pulses and an interpulse delay of 4.2 s with 64 k datapoints and 20 ppm sweep width. 1D 13C spectra were acquired in modulated mode with broadband composite pulse decoupling (WALTZ16) and an interpulse delay of 3.3 s with 64 k datapoints and a sweep width of 240 ppm. Processing and analysis of 1D spectra was performed with Bruker Topspin 3.6 software. No zero filling was performed, and spectra were manually integrated after automatic baseline correction. 13C DEPT modulated spectra were phased so that C, CH2 are positive, and CH, CH3 are negative. 2D HSQC spectra were recorded on all samples to aid the interpretation of the data and to identify signals hidden underneath solvent peaks. Spectra were acquired with sweep widths obtained by automatic sweep width detection from 1D reference spectra in the direct dimension with 1k datapoints and with 210 ppm and 256 datapoints in the indirect dimension. Spectra were processed with Topspin 3.6 software from Bruker and analyzed with ACDlabs NMR workbook 2023.

In certain cases, it was not possible to differentiate between peaks arising from rotamers and those arising from weak intensities due to either slow relaxing quaternary carbons and/or other causes, which is why the number of peaks in the carbon peak list is not always consistent with the number of carbons in the molecule. The same holds true for the proton intensities where due to overlapping rotamers the integrals are sometimes slightly smaller or larger than the expected number of protons.

##### HPLC-MS

All samples were analyzed on an Agilent 1200 series LC system coupled with an Agilent 6140 mass spectrometer. Purity was determined via UV detection with a bandwidth of 170 nm in the range from 230-400nm. LC parameters were as follows: Waters Xbridge C18 column, 2.5 µm particle size, 2.1 x 20 mm. Run time 2.1 minutes, flow 1 ml/min, column temperature 60°C and 5µl injections. Solvent A (20mM NH4HCO3/ NH3 pH 9), solvent B (MS grade acetonitrile). Start 10% B, gradient 10% - 95% B from 0.0 - 1.5 min, 95% B from 1.5 - 2.0 min, gradient 95% - 10% B from 2.0 – 2.1 min.

##### HRMS

The mass calibration was performed using the Pierce LTQ Velos ESI positive ion calibration solution from Thermo Scientific (Product Nr. 88323).

MS parameters: The scan window was set to 200-1200 amu with a maximum injection time of 500 ms and 1 microscan. The resolution of the Orbitrap was 120000 with a mass accuracy ≤ 5 ppm. The ion mode set to positive with a capillary temperature 275 °C and voltage of 60 eV. The tube lens potential was set to 110 eV. 12 NanoESI voltage was 1.45 kV and the N2-gas pressure set to 0.45 psi. Total sample volume was 1 µl and the acquisition time was 0.4 sec, with 10 scans of averaging per spectrum Sample dilution: 10 mM DMSO stock solution was diluted 1:1000 in 25 % acetonitrile +0.01 % formic acid.

#### Chemical synthesis

##### General information

All chemicals unless otherwise stated, were commercially available, at least 90% pure, and used without further purification. Commercially available dry solvents were used. All reactions were carried out under nitrogen atmosphere. Normal phase TLC was carried out on pre-coated silica plates (Kieselgel 60 F254, BDH) with visualization via UV light (UV 254 and/or 365 nm) and/or basic potassium permanganate solution. Isolute® phase separator columns from Biotage were used. Flash column chromatography (FCC) was performed using either a Teledyne Isco Combiflash Rf or Rf200i, or a Biotage Isolera One with prepacked Redisep RF Normal phase disposable Columns. Reverse phase chromatography was carried out on a Biotage Isolera One using SNAP-C18 Columns. NMR Spectra were recorded on a Bruker Ascend 400 MHz or 500 MHz as specified. Chemical shifts are quoted in ppm and referenced to the residual solvent signals: ^1^H NMR δ (ppm) = 7.26 (CDCl_3_), ^13^C NMR δ (ppm) = 77.16 (CDCl_3_); ^1^H NMR δ (ppm) = 5.32 (CD_2_Cl_2_), ^13^C NMR δ (ppm) = 53.84 (CD_2_Cl_2_), ^1^H NMR δ (ppm) = 2.50 (DMSO-d_6_). Signal splitting patterns are described as singlet (s), doublet (d), triplet (t), quartet (q), multiplet (m), broad (br) or a combination thereof. Coupling constants (*J*) are measured in Hertz (Hz). High Resolution Mass Spectra (HRMS) were recorded on a Bruker microTOF. Other resolution MS and analytical HPLC traces were recorded on an Agilent Technologies 1200 series HPLC connected to an Agilent Technologies 6130 quadrupole LC/MS with an Agilent diode array detector. The column used was a Waters XBridge column (50 mm × 2.1 mm, 3.5 μm particle size) and the compounds were eluted with a gradient 5−95% acetonitrile/water + 0.1 formic acid (“acidic method”). Preparative HPLC was performed on a Gilson Preparative HPLC System with a Waters XBridge C18 column (100 mm x 19 mm; 5 μm particle size) and a gradient of 5 % to 95 % acetonitrile in water over 10 minutes, flow 25 mL/min, with 0.1 % ammonia in the aqueous phase.

###### Abbreviations used

EtOAc for ethyl acetate, DMSO for dimethylsulfoxide, MeOH for methanol, EtOH for ethanol, DMF for *N*,*N*-dimethylformamide, THF for tetrahydrofuran, DCE for 1,2-dichloroethane, STAB for sodium triacetoxyborohydride, sat. for saturated, aq. for aqueous.

### P1

**Scheme S1:**
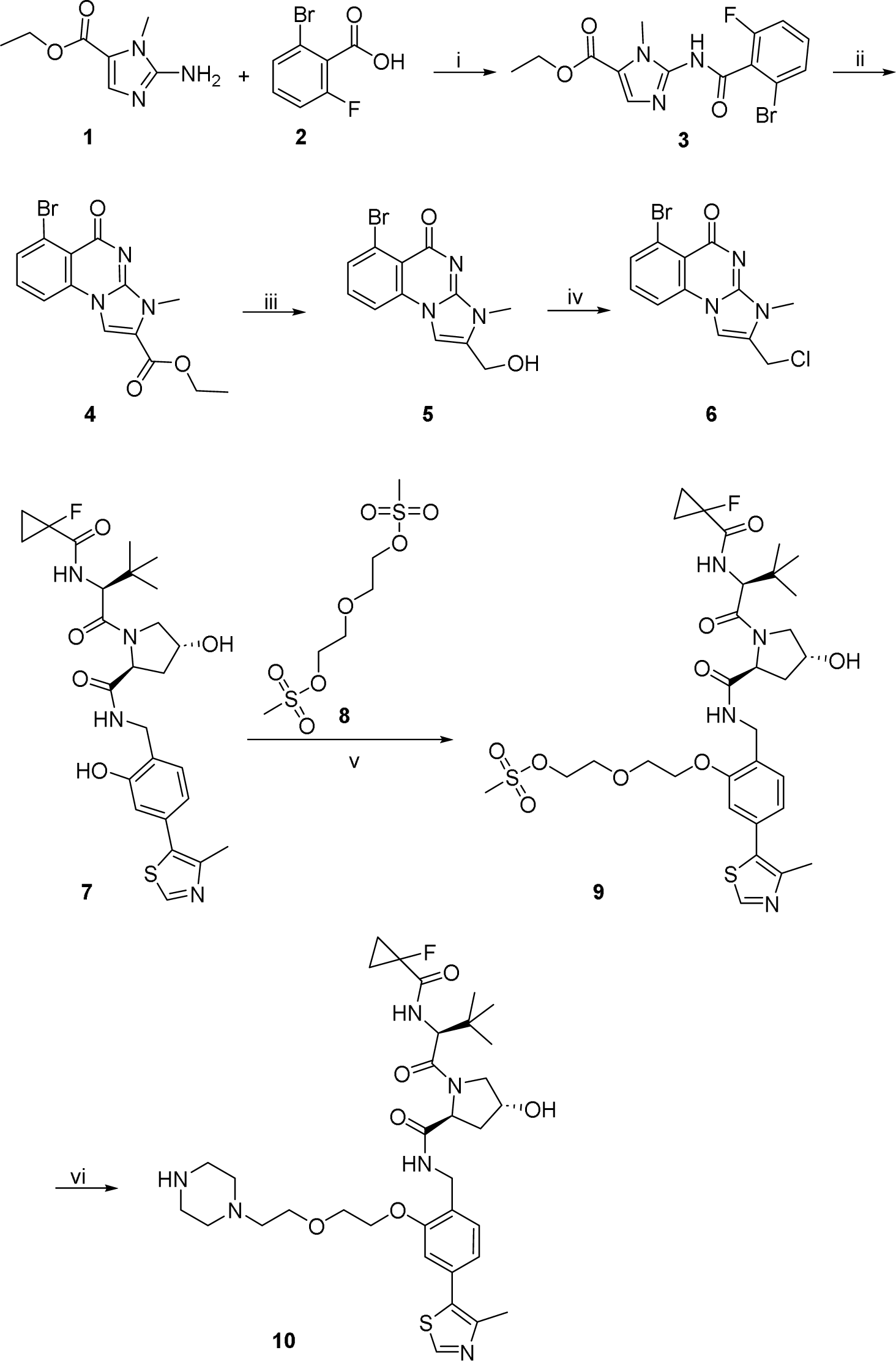
Synthesis of key intermediates for **P1**: i) a) **2**, CDI, dioxane, 95°C; b) **1**, HOBt, dioxane, RT; *then* add a) to b), 105°C; ii) K_2_CO_3_, DMF, 90°C, µw; iii) LiBH_4_, THF/MeOH, RT; iv) MsCl, TEA, DCE/DMSO, 50°C; v) **8**, K_2_CO_3_, DMF, 60°C; vi) piperazine, TEA, KI, MeCN, 80°C.

#### ethyl 2-(2-bromo-6-fluorobenzamido)-1-methyl-1H-imidazole-5-carboxylate 3

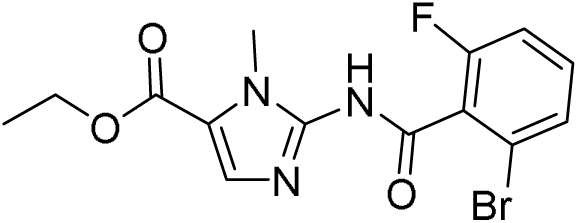

A solution of 2-Bromo-6-fluorobenzoic acid **2** (11.9 g, 54.4 mmol, 1.15 eq.) and CDI (11.5 g, 70.9 mmol, 1.50 eq.) in 1,4-dioxane (64 mL) was stirred at 95°C for 1 h.

In a second vessel, a solution of ethyl 2-amino-1-methyl-1H-imidazole-5-carboxylate **1** (8.0 g, 47.3 mmol) and HOBt (10.2 g, 75.7 mmol, 1.60 eq.) was stirred at RT for 1 h. The activated acid was added to this solution and the resulting mixture was stirred at 105°C for 12 h. The solvent was evaporated, and the crude residue was purified by silica gel column chromatography (0-100% EtOAc in petroleum ether) and again by acidic HPLC-chromatography to obtain **3** (9.0 g, 51% yield).

#### ethyl 6-bromo-3-methyl-5-oxo-3H,5H-imidazo[1,2-a]quinazoline-2-carboxylate 4

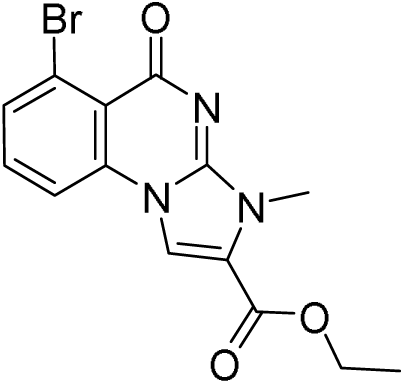

To a solution of **3** (8.5 g, 23.0 mmol) in DMF (85 mL), potassium carbonate (9.5 g, 68.9 mmol, 3.0 eq.) was added and the mixture was stirred at 90°C for 12 h. It was quenched with sat. NH_4_Cl-solution and the resulting precipitate was filtered off, rinsed with EtOAc and dried to give **4** (5.0 g, 58% yield), which was used in the next step without further purification.

#### 6-bromo-2-(hydroxymethyl)-3-methylimidazo[1,2-a]quinazolin-5(3H)-one 5

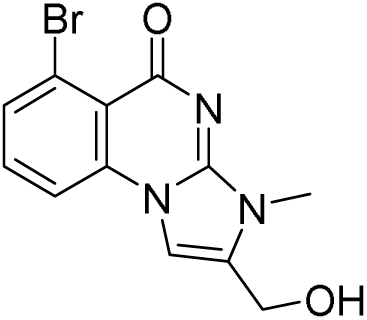

To a solution of **4** (5.0 g, 14.3 mmol) in THF (50 mL) and MeOH (5 mL), lithium borohydride (2M in THF) (14.3 mL, 28.6 mmol, 2.0 eq.) was added dropwise at 0°C over 30 min. After complete addition, the mixture was stirred at RT for 30 min. It was cooled to 0°C and quenched by dropwise addition of 30 mL 1M aq. HCl-solution. The mixture was concentrated under reduced pressure and the residue was purified by acidic prep. HPLC-chromatography to obtain **5** (2.4 g, 53% yield).

#### 6-bromo-2-(chloromethyl)-3-methylimidazo[1,2-a]quinazolin-5(3H)-one 6

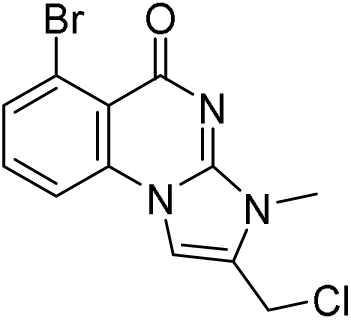

To a solution of **5** (2.0 g, 6.5 mmol) in chloroform (20 mL), thionyl chloride (2.4 mL, 33 mmol, 5.0 eq.) was added dropwise at RT over 15 min followed by dropwise addition of DMF (505 µL, 6.5 mmol, 1.0 eq.). The resulting mixture was stirred at RT for 2 h and concentrated under reduced pressure. The crude residue was purified by column chromatography to give **6** (0.8 g, 24% yield), which was immediately used in the following step.

#### 2-(2-(2-(((2*S*,4*R*)-1-((*S*)-2-(1-fluorocyclopropane-1-carboxamido)-3,3-dimethylbutanoyl)-4-hydroxypyrrolidine-2-carboxamido)methyl)-5-(4-methylthiazol-5-yl)phenoxy)ethoxy)ethyl methanesulfonate 9

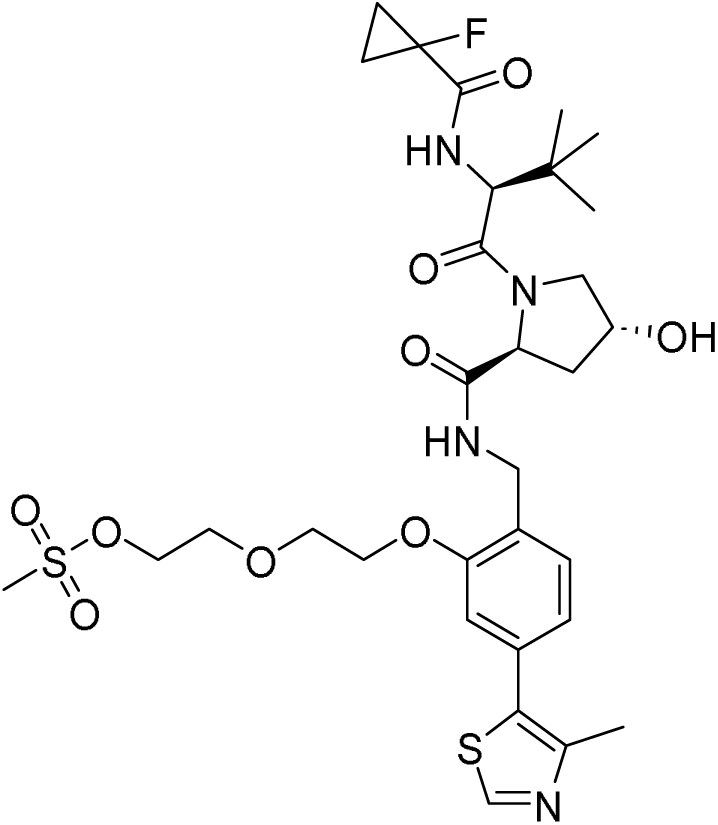

To a solution of **7** (500 mg, 939 µmol) and **8** (370 mg, 1.41 mmol, 1.5 eq.) in DMF (10 mL), potassium carbonate (390 mg, 2.82 mmol, 3.0 eq.) was added. The resulting mixture was stirred at 60°C for 12 h. It was poured into ice-water, stirred for 30 min and extracted with EtOAc. The combined organic layers were washed with brine, dried and concentrated to obtain crude **9** (600 mg, 60% yield), which was taken to the next step without further purification.

#### (2*S*,4*R*)-1-((*S*)-2-(1-fluorocyclopropane-1-carboxamido)-3,3-dimethylbutanoyl)-4-hydroxy-N-(4-(4-methylthiazol-5-yl)-2-(2-(2-(piperazin-1-yl)ethoxy)ethoxy)benzyl)pyrrolidine-2-carboxamide 10

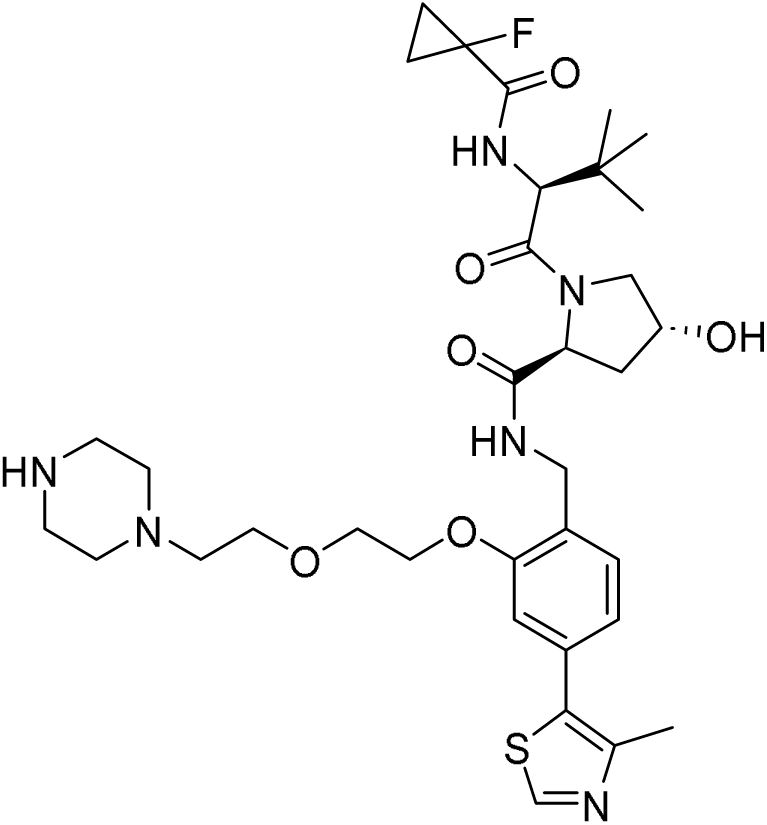

To a solution of **9** (600 mg, 562 µmol) and piperazine (145 mg, 1.68 mmol, 3.0 eq.) in MeCN (12 mL), TEA (235 µL, 1.69 mmol, 3.0 eq.) and potassium iodide (47 mg, 283 µmol, 0.5 eq.) were added. The resulting mixture was stirred at 80°C for 12 h. It was diluted with 1 mL water and purified by acidic HPLC-chromatography to give **10** (230 mg, 58% yield).

**Scheme S2:**
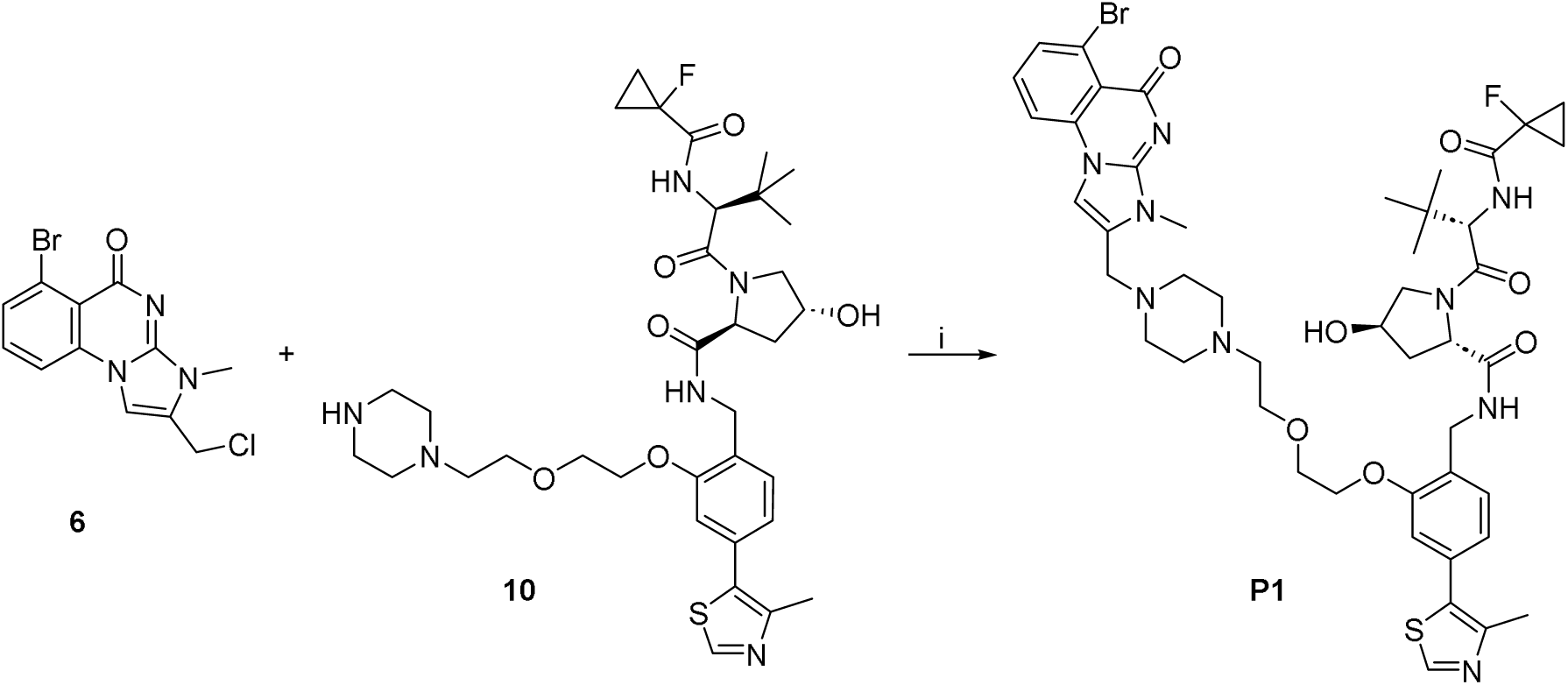
Synthesis of P1: i) K_2_CO_3_, KI, DMF, 100°C.

#### (2*S*,4*R*)-N-(2-(2-(2-(4-((6-bromo-3-methyl-5-oxo-3,5-dihydroimidazo[1,2-a]quinazolin-2-yl)methyl)piperazin-1-yl)ethoxy)ethoxy)-4-(4-methylthiazol-5-yl)benzyl)-1-((*S*)-2-(1-fluorocyclopropane-1-carboxamido)-3,3-dimethylbutanoyl)-4-hydroxypyrrolidine-2-carboxamide P1

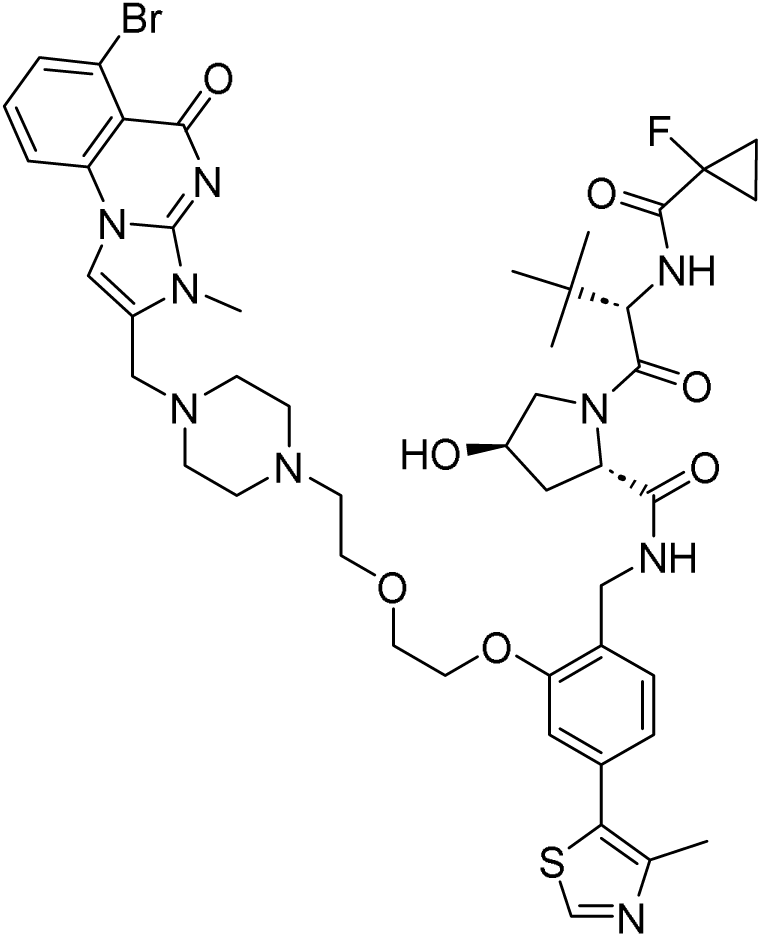

To a solution of **6** (346 mg, 668 µmol, 2.0 eq.) and **10** (230 mg, 334 µmol) in DMF (20 mL), potassium carbonate (230 mg, 1.66 mmol, 5.0 eq.) and potassium iodide (160 mg, 964 µmol, 2.89 eq.) were added. The resulting mixture was stirred at 100°C for 2 h. It was filtered and the filtrate was purified by acidic prep. HPLC to obtain **P1** (38 mg, 11% yield).

##### HRMS (ESI+) *m*/*z*

[M+H]^+^ calcd for C_46_H_57_BrFN_9_O_7_S 978.32815; found 978.33543

##### Analytical ^1^H NMR note

The analytical sample contains a protonated form of **P1** which affects the interchangeable peaks NH (broadened) and OH (not apparent).

##### ^1^H NMR (DMSO-d_6_) δ

10.71-10.96 (m, 1H), 9.02 (s, 2H), 8.71 (t, J=5.5 Hz, 0.1H rotamer), 8.58 (t, J=6.0 Hz, 1H), 8.40 (s, 2H), 8.14 (d, J=8.2 Hz, 1H), 7.90 (d, J=7.9 Hz, 1H), 7.79 (t, J=8.0 Hz, 1H), 7.41 (d, J=7.9 Hz, 1H), 7.28 (dd, J=9.1, 2.5 Hz, 1H), 7.20 (s, 0.1H rotamer), 7.03 (s, 1H), 6.98 (dd, J=7.9, 0.9 Hz, 1H), 4.71 (br t, J=7.4 Hz, 0.1H rotamer), 4.59 (d, J=9.1 Hz, 1H), 4.53 (t, J=8.2 Hz, 1H), 4.44 (br d, J=8.8 Hz, 0.1H rotamer), 4.36 (br s, 1H), 4.30 (dd, J=16.4, 6.0 Hz, 1H), 4.16-4.26 (range, 3H), 3.94 (br t, J=4.4 Hz, 2H), 3.81-3.88 (range, 4H), 3.69 (s, 3H), 3.32-3.39 (range, 3H), 3.04-3.26 (range, 5H), 2.69-2.84 (range, 2H), 2.46 (s, 3H), 2.11 (br dd, J=12.3, 7.9 Hz, 1H), 1.91 (ddd, J=12.9, 8.8, 4.4 Hz, 1H), 1.29-1.43 (m, 2H), 1.22 (dd, J=8.5, 3.2 Hz, 2H), 0.95 (s, 9H)

##### ^13^C NMR (DMSO-d_6_) δ

172.4, 169.4, 168.6 (d, CF=21 Hz), 161.3, 156.1, 152.1, 148.1, 137.0, 135.2, 134.2, 131.8, 131.3, 128.2, 127.6, 123.4, 121.5, 112.4, 78.6 (d, CF=233 Hz), 69.3, 69.4, 67.9, 65.2, 59.3, 57.2, 57.0, 51.4, 49.5, 48.7, 40.9, 40.8, 40.6, 40.6, 38.4, 37.7, 36.5, 30.4, 26.6, 16.4, 13.5 (d, CF=10 Hz), 13.2 (d, CF=11 Hz)

##### HPLC

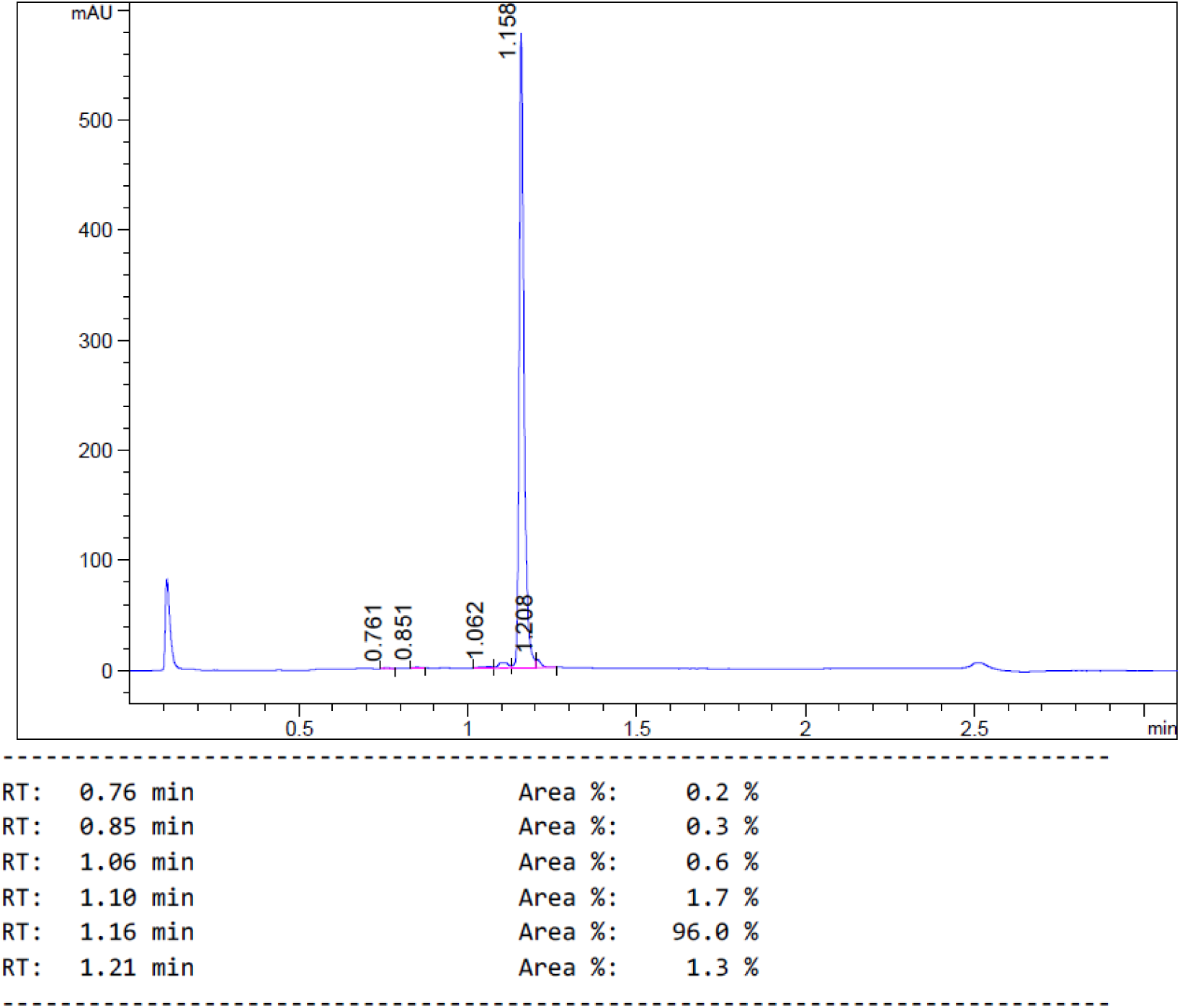

##### ^1^H and ^13^C NMR

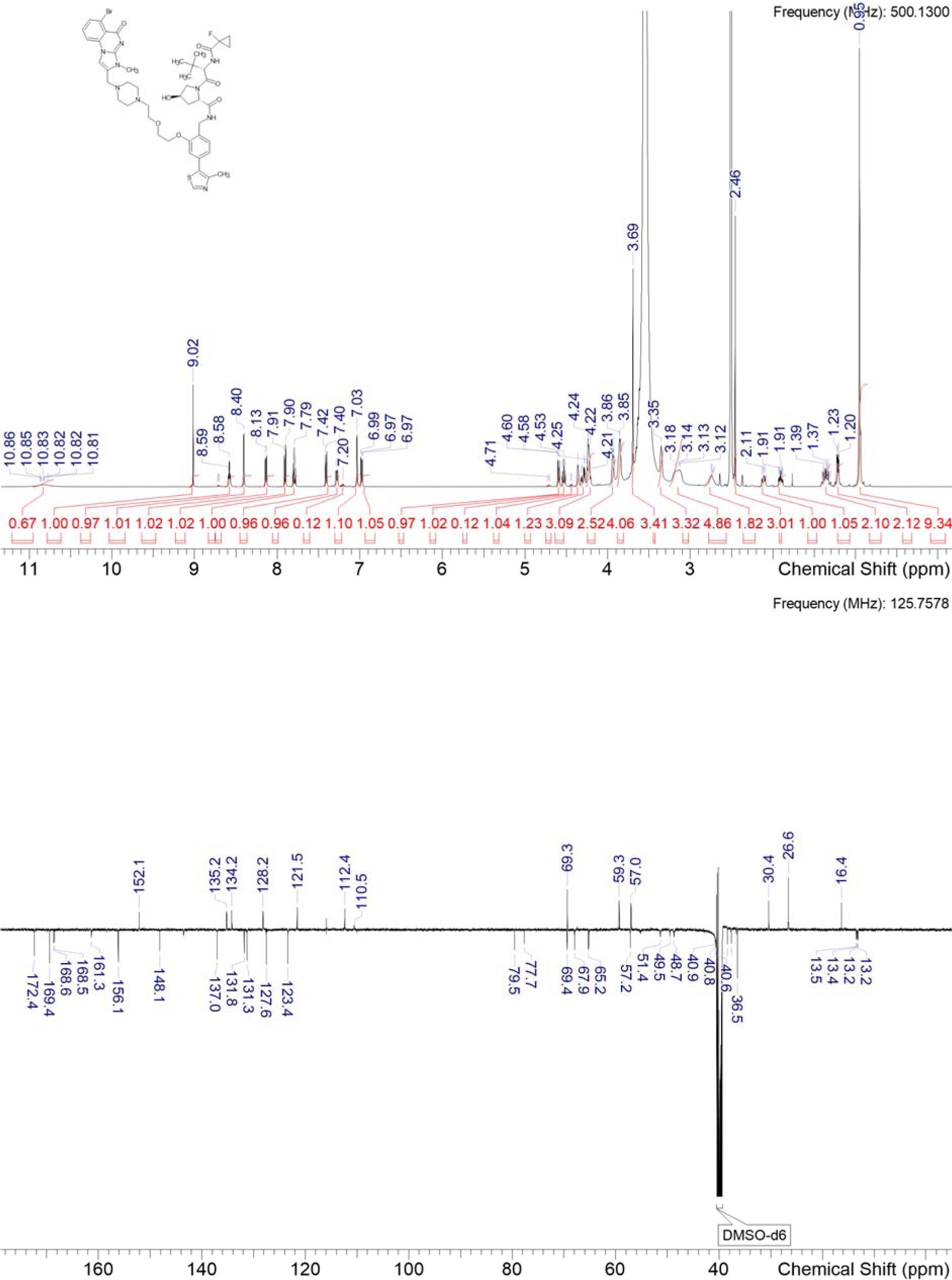

### P3

**Scheme S3:**
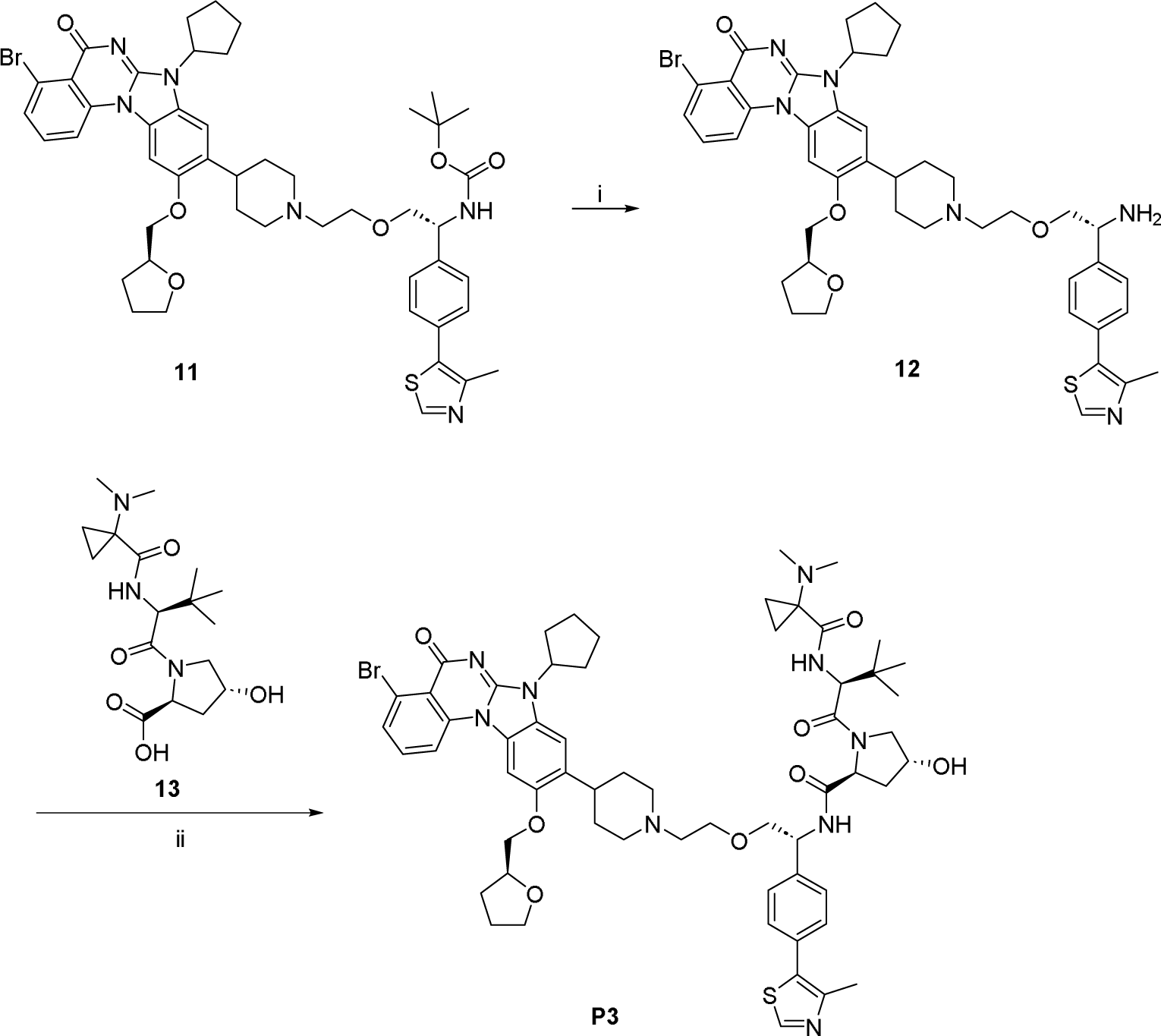
Synthesis of P3: i) conc. HCl, MeOH, RT; ii) intermediate **13**, HATU, TEA, DMF, RT.

#### (2*S*,4*R*)-N-((*R*)-2-(2-(4-(4-bromo-7-cyclopentyl-5-oxo-10-(((*S*)-tetrahydrofuran-2-yl)methoxy)-5,7-dihydrobenzo[4,5]imidazo[1,2-a]quinazolin-9-yl)piperidin-1-yl)ethoxy)-1-(4-(4-methylthiazol-5-yl)phenyl)ethyl)-1-((*S*)-2-(1-(dimethylamino)cyclopropane-1-carboxamido)-3,3-dimethylbutanoyl)-4-hydroxypyrrolidine-2-carboxamide P3

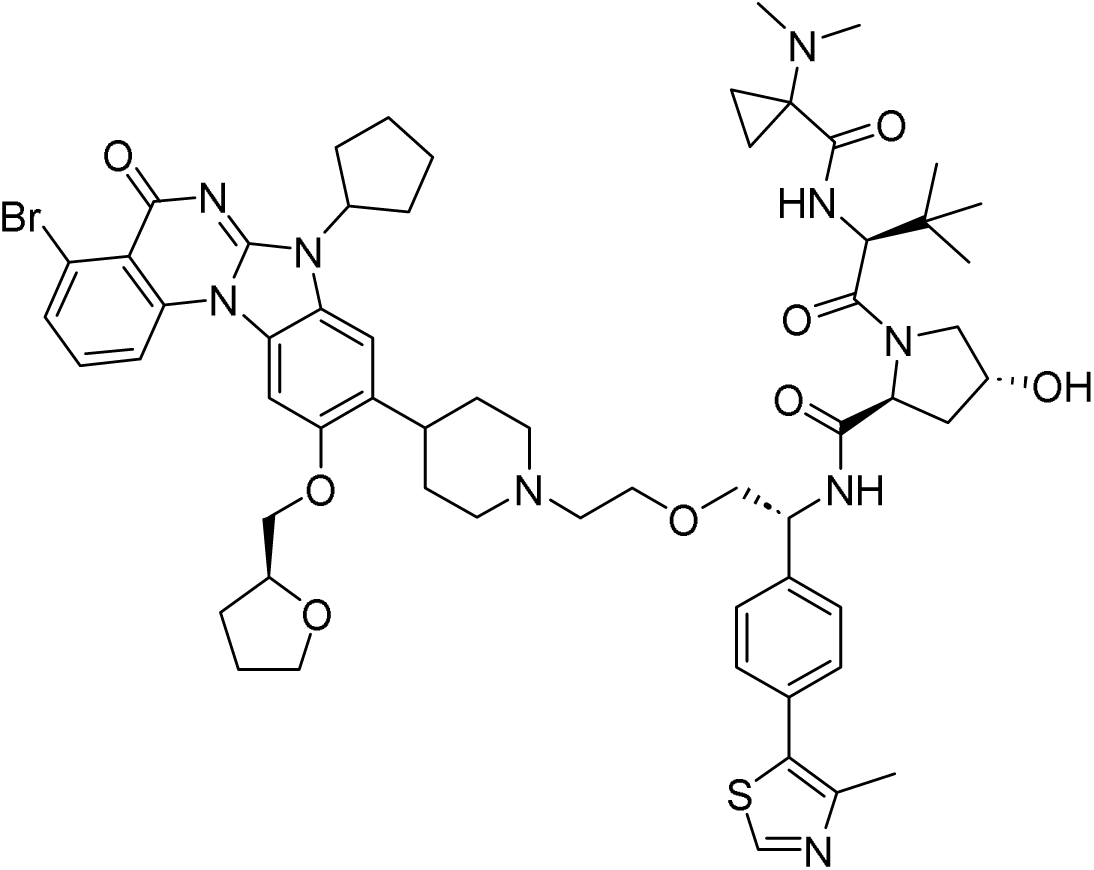

**11** (42 mg, 45 µmol) was taken up in methanol (1000 µL) and conc. aq. HCl (1000 µL) was added. The mixture was stirred at RT for 1 h. The solvents were removed under reduced pressure to obtain crude **12**.

**13** (24 mg, 68 µmol, 1.5 eq.) was dissolved in DMF (1000 µL) and TEA (32 µL, 0.23 mmol, 5.0 eq.) and HATU (52 mg, 0.14 mmol, 3.0 eq.) were added. The resulting mixture was stirred at RT for 30 minutes, then **12** (39 mg, 45 µmol) was added and stirring was continued at RT for 1 h. The reaction was diluted with water, filtered through a syringe filter and purified by basic HPLC-chromatography to obtain **P3** (32 mg, 61% yield).

##### HRMS (ESI+) *m*/*z*

[M+H]^+^ calcd for C_60_H_76_BrN_9_O_8_S 1162.47441; found 1162.48169

##### ^1^H NMR (DMSO-d_6_) δ

8.96 (s, 1H), 8.95 (br s, 0.1H rotamer), 8.69 (d, J=7.9 Hz, 0.1H rotamer), 8.51 (d, J=7.9 Hz, 1H), 8.48 (d, J=8.4 Hz, 1H), 8.12 (d, J=9.7 Hz, 1H), 7.95 (d, J=9.2 Hz, 0.1H rotamer), 7.83 (s, 1H), 7.80 (d, J=7.9 Hz, 1H), 7.69 (t, J=8.2 Hz, 1H), 7.44 (s, 4H), 7.41 (s, 1H), 7.38 (s, 0.1H rotamer), 5.25 (quin, J=8.7 Hz, 1H), 5.14 (d, J=3.5 Hz, 1H), 5.00-5.06 (m, 1H), 4.98 (d, J=2.8 Hz, 0.1H rotamer), 4.91-4.96 (m, 0.1H rotamer), 4.75 (t, J=7.4 Hz, 0.1H rotamer), 4.52 (t, J=8.3 Hz, 1H), 4.48 (d, J=9.7 Hz, 1H), 4.34 (d, J=9.4 Hz, 0.1H rotamer), 4.27-4.31 (m, 1H), 4.17-4.27 (range, 3H), 3.81-3.88 (m, 1H), 3.72-3.77 (m, 1H), 3.64-3.70 (m, 1H), 3.52-3.64 (range, 5H), 3.43-3.48 (m, 0.1H rotamer), 3.37-3.42 (m, 0.1H rotamer), 2.94-3.05 (range, 3H), 2.44 (s, 3H), 2.19-2.26 (m, 2H), 2.16 (s, 6H), 1.93-2.12 (range, 10H), 1.76-1.92 (range, 3H), 1.68-1.76 (range, 6H), 1.00-1.04 (m, 1H), 0.96 (s, 9H), 0.93-0.95 (m, 3H)

##### ^13^C NMR (DMSO-d_6_) δ

172.6, 171.5, 169.7, 164.4, 152.8, 152.0, 149.0, 148.2, 141.2, 139.1, 133.8, 132.7, 131.5, 130.5, 129.1, 127.9, 125.3, 124.6, 123.2, 117.1, 116.0, 109.0, 99.9, 77.2, 73.4, 72.5, 69.3, 69.0, 68.2, 59.0, 57.8, 57.0, 56.8, 54.9, 54.3, 52.5, 47.9, 42.1, 40.5, 38.2, 36.6, 35.7, 32.2, 28.6, 28.1, 26.8, 26.7, 26.1, 25.2, 16.4, 12.8, 10.8

##### HPLC

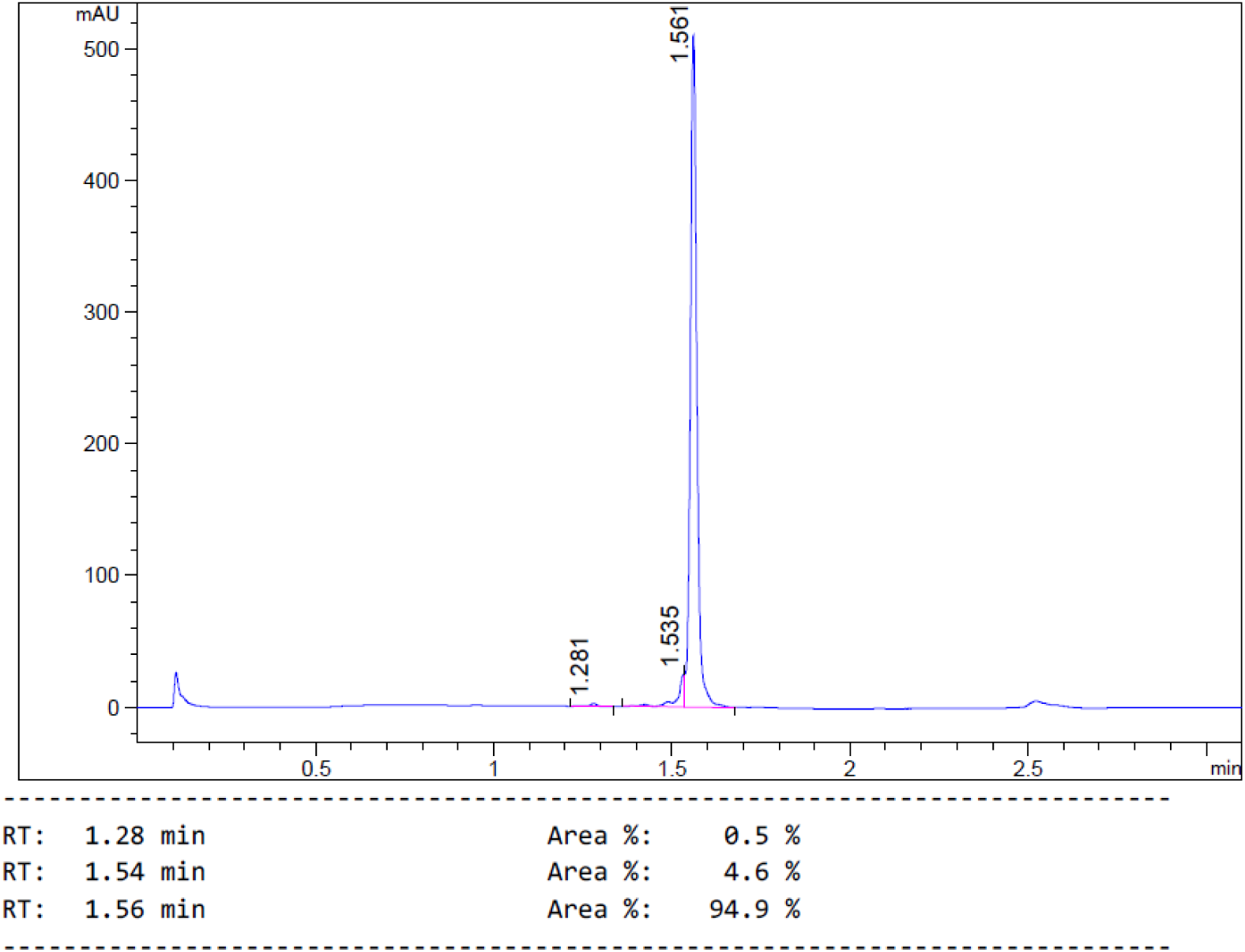

##### ^1^H and ^13^C NMR

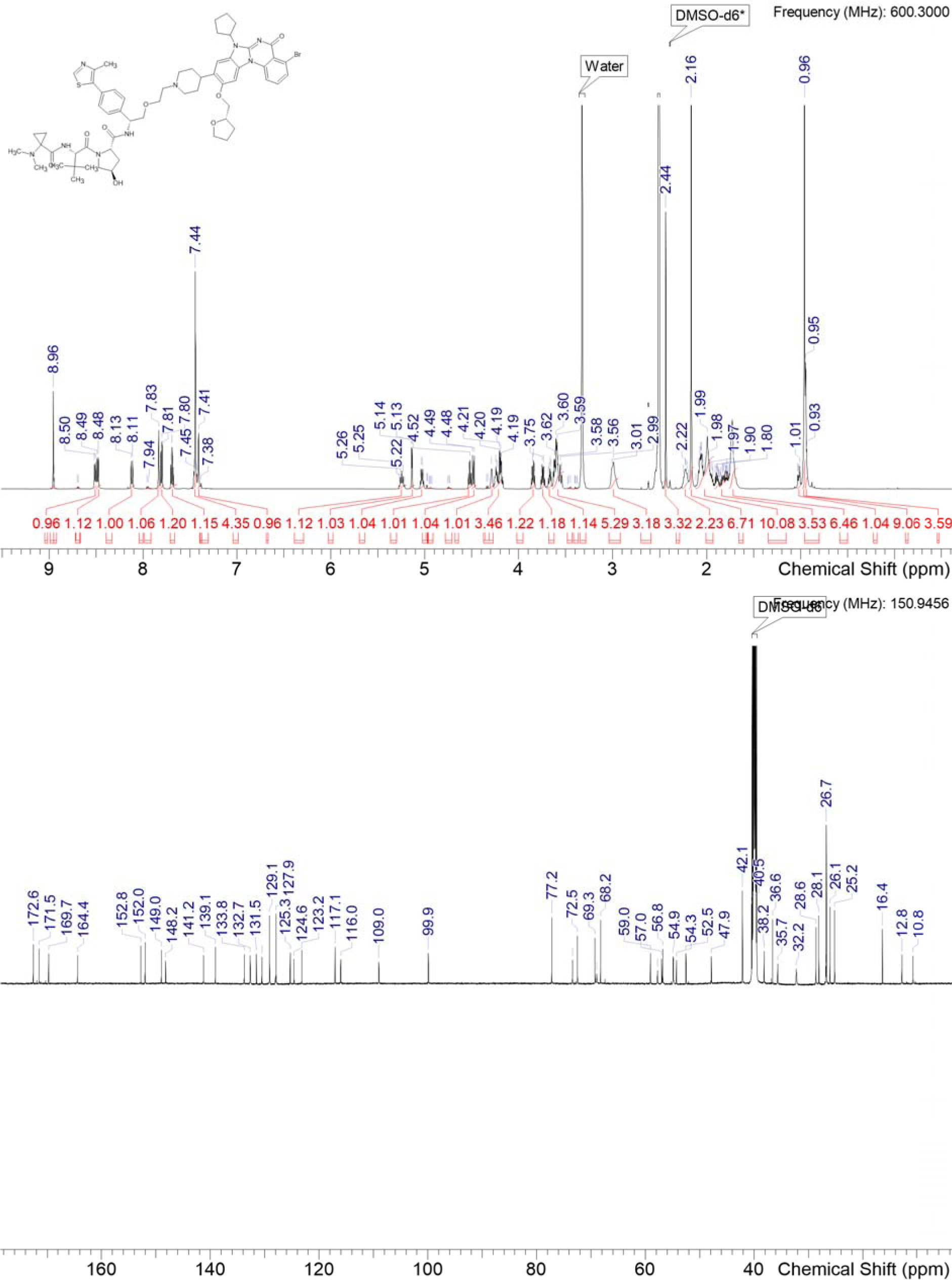

### P4

**Scheme S4:**
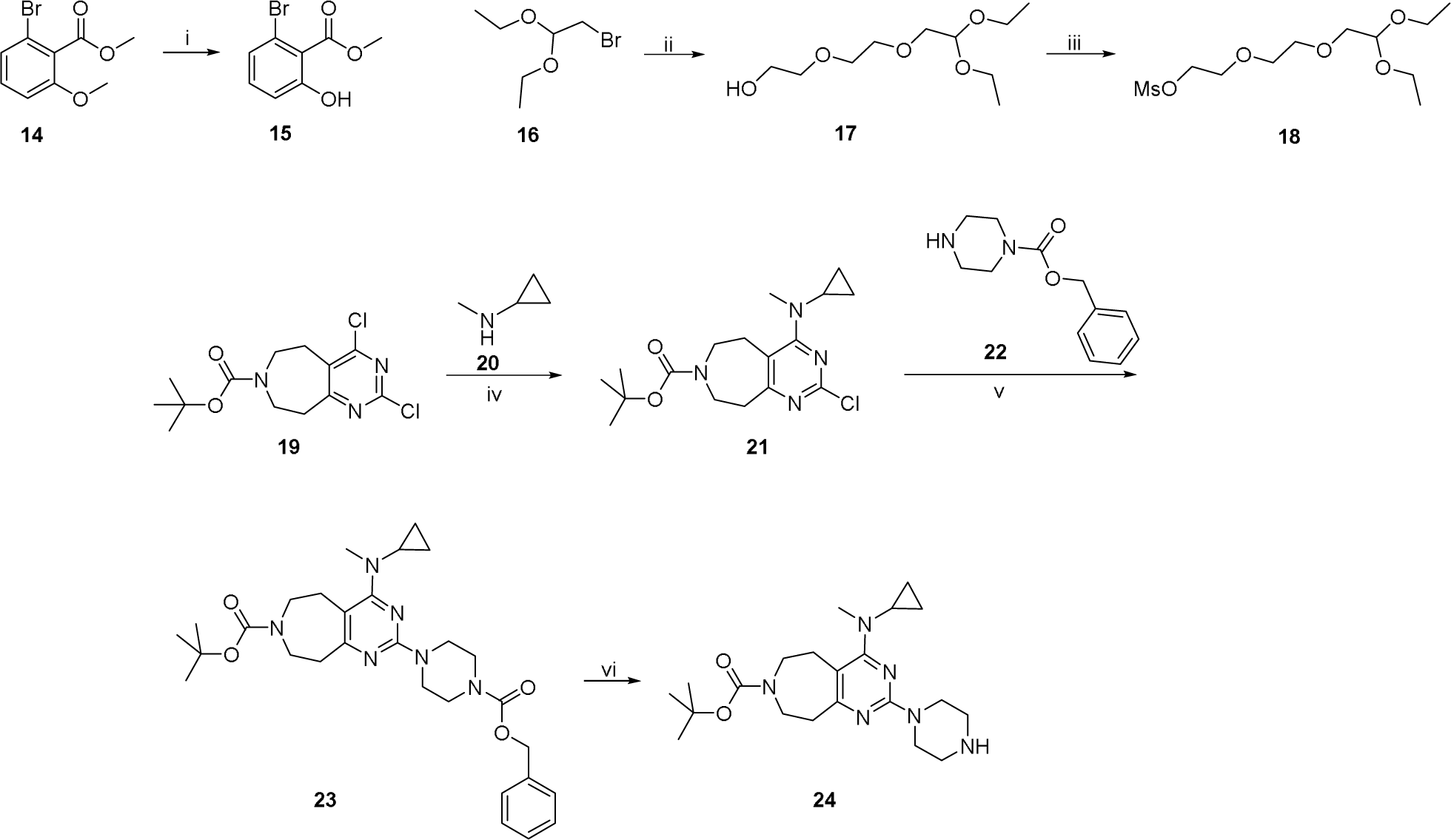
Synthesis of key intermediates for **P4**: i) BBr_3_, DCM, RT; ii) diethylene glycol, KOH, 115 °C; iii) methanesulfonic anhydride, TEA, DCM, 0 °C-RT; iv) **20**, TEA, DMSO, RT; v) **22**, DIPEA, NMP, 150 °C, µw; vi) Pd(OH)_2_/C, H_2_, MeOH, RT.

#### methyl 2-bromo-6-hydroxybenzoate 15

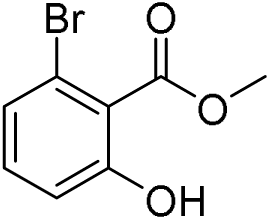

A solution of boron tribromide (1M in DCM, 31.6 mL, 31.6 mmol, 2.5 eq.) in DCM (15 mL) was cooled to 0°C and a solution of methyl 2-bromo-6-methoxybenzoate **14** (3.1 g, 12.6 mmol) in DCM (60 mL) was added dropwise. The mixture was stirred at 0°C for 1 h and quenched with water. It was diluted with DCM and the layers were separated. The aqueous layer was extracted with EtOAc. The combined organics were dried and concentrated. The crude residue was purified by silica gel column chromatography (0-15% EtOAc in cyclohexane) to obtain **15** (2.1 g, 72% yield).

#### 2-[2-(2,2-Diethoxy-ethoxy)-ethoxy]-ethanol 17

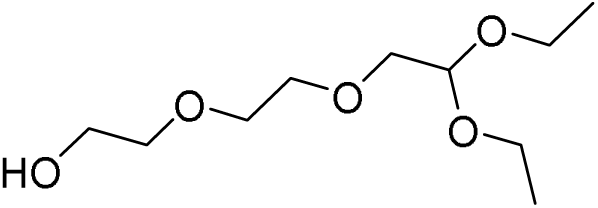

Potassium hydroxide (1.0 g, 18 mmol, 2.7 eq) was added to diethylene glycol (4.7 g, 44 mmol, 6.6 eq.) at 80°C and the resulting mixture was stirred until all solids were dissolved. Then 2-Bromo-1,1-diethoxy-ethane **16** (1.3 g, 6.7 mmol) was added, and the mixture was stirred at 115°C for 72 h. It was cooled to RT, quenched with water, extracted with DCM, dried over MgSO_4_ and concentrated. The crude residue was purified by silica gel column chromatography (0-10% MeOH in DCM) to obtain **17** (0.8 g, 54% yield).

#### 2-[2-(2,2-diethoxyethoxy)ethoxy]ethyl methanesulfonate 18

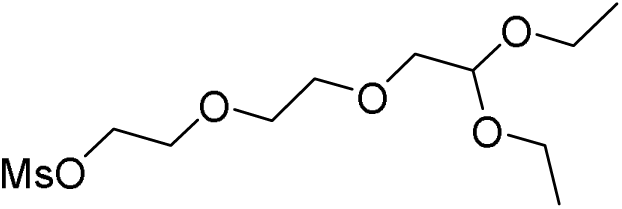

**17** (917 mg, 4.1 mmol) was dissolved in DCM (9 mL) and TEA (1.2 mL, 8.3 mmol, 2.0 eq.) was added. It was cooled to 0°C and methanesulfonic anhydride (740 mg, 4.1 mmol, 1.0 eq.) was added. The resulting mixture was stirred at RT for 2 h and quenched with water. It was extracted with DCM, washed with water and brine, dried and concentrated to give crude **18** (513 mg, 41% yield), which was used in the next step without further purification.

#### 2-Chloro-4-(cyclopropyl-methyl-amino)-5,6,8,9-tetrahydro-pyrimido[4,5-d]azepine-7-carboxylic acid tert-butyl ester 21

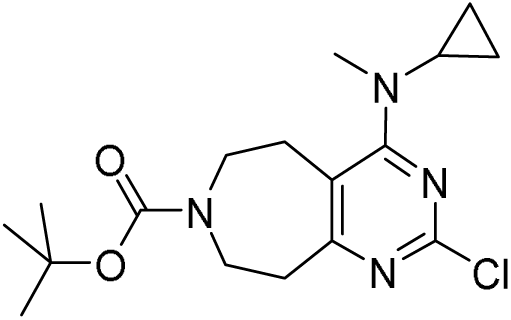

To a solution of 2,4-Dichloro-5,6,8,9-tetrahydro-pyrimido[4,5-d]azepine-7-carboxylic acid tert-butyl ester **19** (100 mg, 314 µmol) in DMSO (1000 µL), TEA (131 µL, 943 µmol, 3.0 eq.) and Cyclopropyl-methyl-amine **20** (45 mg, 629 µmol, 2.0 eq.) were added. The resulting mixture was stirred at RT for 16 h. The reaction was diluted with MeCN and water, filtered through a syringe filter and purified by acidic HPLC-chromatography to obtain **21** (82 mg, 74% yield).

#### benzyl 4-{7-[(tert-butoxy)carbonyl]-4-[cyclopropyl(methyl)amino]-5H,6H,7H,8H,9H-pyrimido[4,5-d]azepin-2-yl}piperazine-1-carboxylate 23

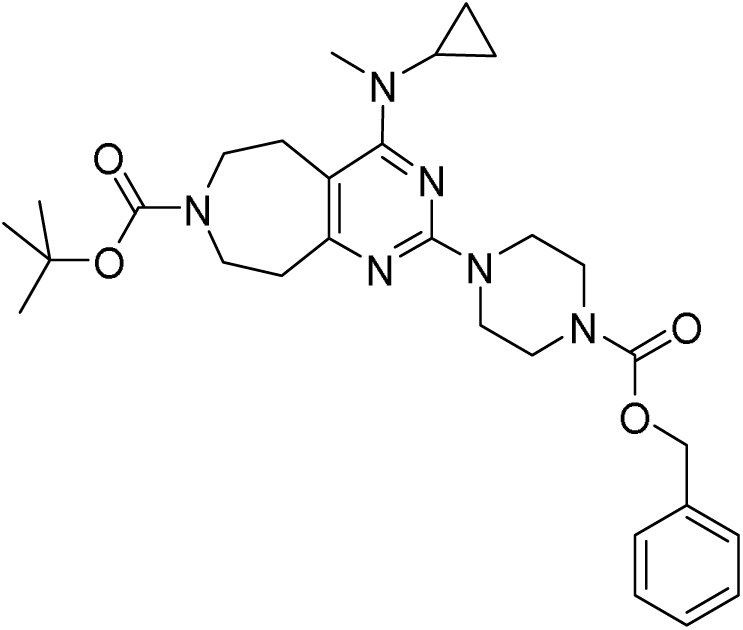

To a solution of **21** (233 mg, 584 µmol) in NMP (1000 µL), DIPEA (124 µL, 730 µmol, 1.3 eq.) and 1-(Benzyloxycarbonyl)piperazine **22** (287 µL, 1.46 mmol, 2.5 eq.) were added. The resulting mixture was irradiated in the microwave at 150°C for 1.5 h. The mixture was diluted with sat. aq. potassium carbonate solution and extracted with DCM. The combined organic layers were dried and concentrated. The crude was dissolved in MeCN and water, filtered through a syringe filter and purified by acidic HPLC-chromatography to give **23** (164 mg, 52% yield).

#### tert-butyl 4-[cyclopropyl(methyl)amino]-2-(piperazin-1-yl)-5H,6H,7H,8H,9H-pyrimido[4,5-d]azepine-7-carboxylate 24

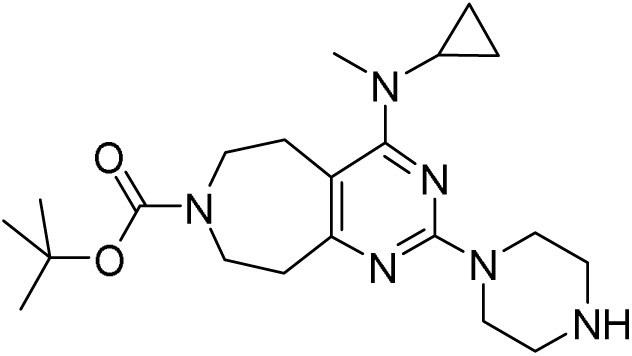

To a solution of **23** (164 mg, 306 µL) in MeOH (10 mL) in a steel reactor, palladium hydroxide (20% on charcoal) (69 mg, 98 µmol, 0.32 eq.) was added. The reactor was flushed with nitrogen and filled with hydrogen. The reaction mixture was stirred under 6 bar of hydrogen atmosphere for 2.5 h. The catalyst was filtered off over a Celite pad and rinsed with MeOH. The filtrate was concentrated under reduced pressure and the resulting crude **24** was taken to the next step without further purification.

**Scheme S5:**
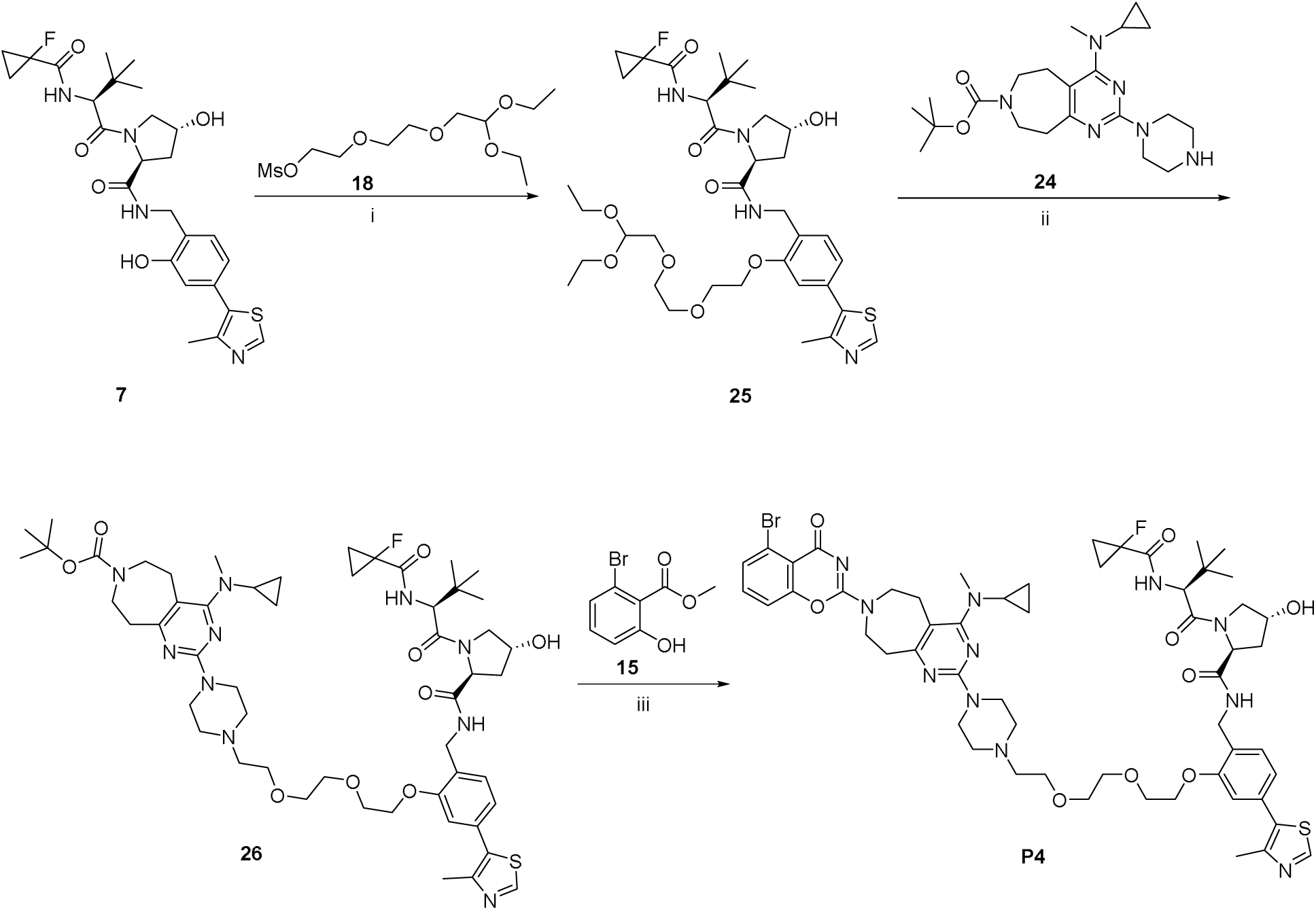
Synthesis of P4: i) **18**, K_2_CO_3_, DMF, 75°C; ii) (a) 1M HCl, dioxane, 50°C; (b) TEA, **24**, DMF *then* STAB, RT; iii) (a) 4M HCl, dioxane, RT; (b) **15**, TEA, CNBr, acetone, 0°C-RT, *then* (a).

#### (2S,4R)-N-[(2-{2-[2-(2,2-diethoxyethoxy)ethoxy]ethoxy}-4-(4-methyl-1,3-thiazol-5-yl)phenyl)methyl]-1-[(2S)-2-[(1-fluorocyclopropyl)formamido]-3,3-dimethylbutanoyl]-4-hydroxypyrrolidine-2-carboxamide 25

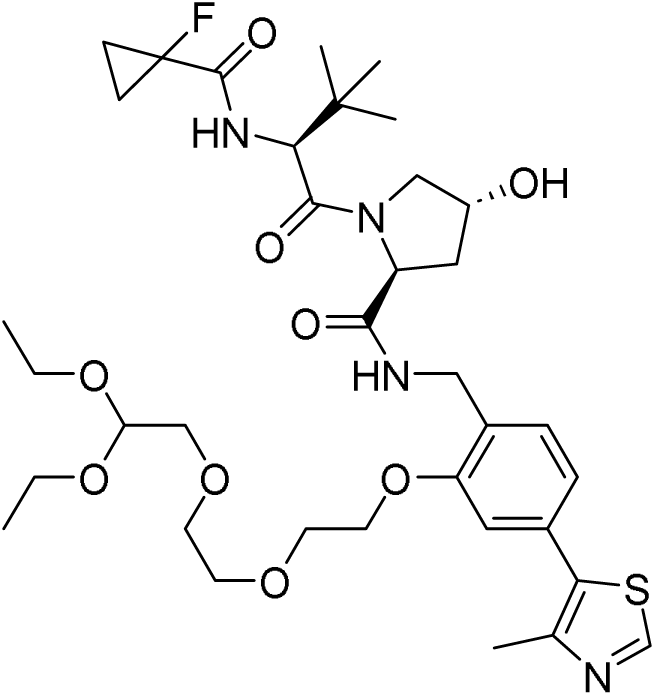

To a solution of **7** (100 mg, 188 µmol) in DMF (1000 µL), potassium carbonate (65 mg, 469 µmol, 2.5 eq.) and **18** (85 mg, 282 µmol, 1.5 eq.) were added and the resulting mixture was stirred at 75°C for 11 h. It was diluted with MeCN and water, filtered through a syringe filter and purified by basic HPLC-chromatography to give **25** (68 mg, 49% yield).

#### tert-butyl 4-[cyclopropyl(methyl)amino]-2-{4-[2-(2-{2-[2-({[(2S,4R)-1-[(2S)-2-[(1-fluorocyclopropyl)formamido]-3,3-dimethylbutanoyl]-4-hydroxypyrrolidin-2-yl]formamido}methyl)-5-(4-methyl-1,3-thiazol-5-yl)phenoxy]ethoxy}ethoxy)ethyl]piperazin-1-yl}-5H,6H,7H,8H,9H-pyrimido[4,5-d]azepine-7-carboxylate 26

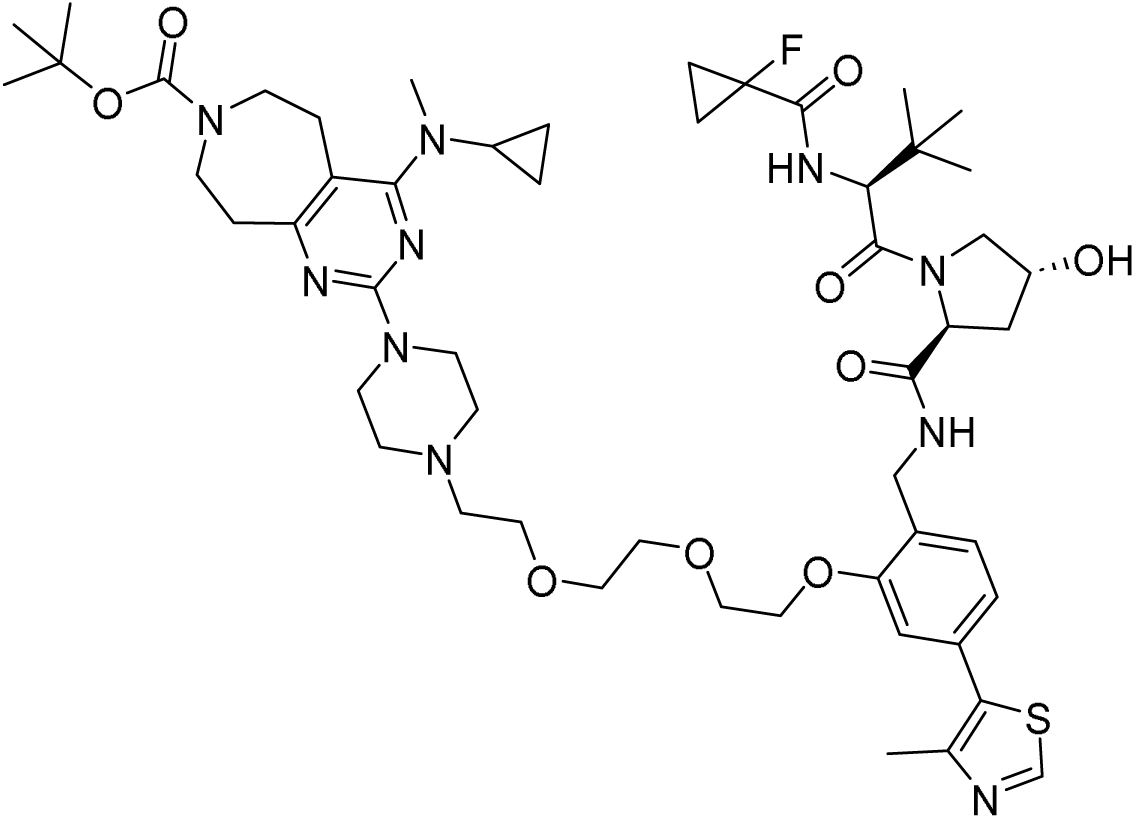

To a solution of **25** (100 mg, 122 µmol) in 1,4-dioxane (2 mL), 1M aq. HCl (122 µL, 122 µmol, 1.0 eq.) was added and the mixture was stirred at 50°C for 16 h. Then TEA (188 µL, 1.4 mmol, 11 eq.) was added, followed by a solution of **24** (55 mg, 137 µmol, 1.1 eq.) in DMF (550 µL) and a a tip of spatula of Mg_2_SO_4_. The resulting mixture was stirred at RT for 5 min before STAB (153 mg, 700 µmol, 5.7 eq.) was added and stirring was continued at RT for 1 h. The reaction was quenched with water and the solvents were removed under reduced pressure. The residue was dissolved in MeCN and water, filtered through a syringe filter and purified by basic HPLC-chromatography to give **26** (43 mg, 34% yield).

#### (2S,4R)-N-[(2-{2-[2-(2-{4-[7-(5-bromo-4-oxo-4H-1,3-benzoxazin-2-yl)-4-[cyclopropyl(methyl)amino]-5H,6H,7H,8H,9H-pyrimido[4,5-d]azepin-2-yl]piperazin-1-yl}ethoxy)ethoxy]ethoxy}-4-(4-methyl-1,3-thiazol-5-yl)phenyl)methyl]-1-[(2S)-2-[(1-fluorocyclopropyl)formamido]-3,3-dimethylbutanoyl]-4-hydroxypyrrolidine-2-carboxamide P4

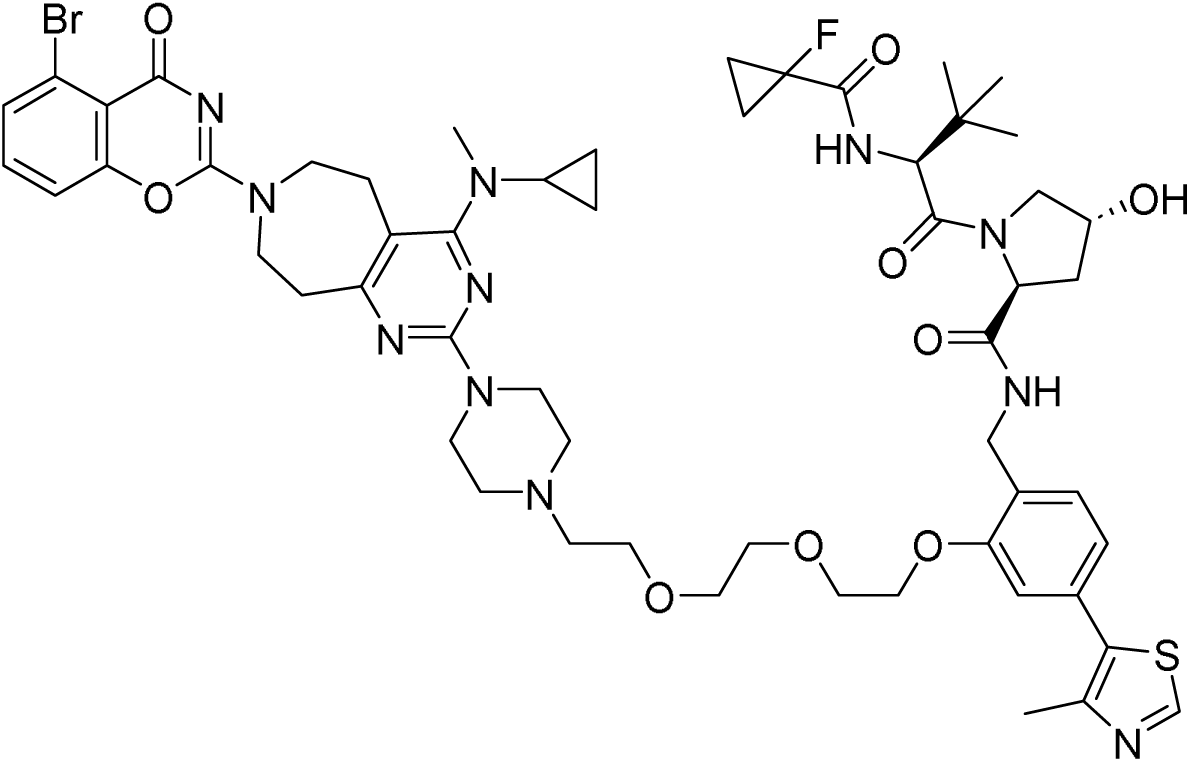

A solution of **15** (14 mg, 61 µmol, 1.5 eq.) and TEA (29 µL, 205 µmol, 5.0 eq.) in acetone (0.5 mL) was cooled to 0°C and cyanogen bromide (5M in MeCN) (12 µL, 61 µmol, 1.5 eq.) was added slowly. The mixture was stirred at RT for 1 h.

In a second vessel, **26** (43 mg, 41 µmol) was taken up in 4M HCl in dioxane (2 mL, 8 mmol, 195 eq.) and stirred at RT for 10 min. It was concentrated under reduced pressure, the residue was dissolved in acetone (1 mL) and added dropwise to the above reaction mixture.

The resulting reaction mixture was stirred at RT for 1 h. It was concentrated to dryness, the residue was dissolved in MeCN and water, filtered through a syringe filter and purified by basic HPLC-chromatography to give **P4** (43 mg, 34% yield).

##### HRMS (ESI+) *m*/*z*

[M+H]^+^ calcd for C_56_H_71_BrFN_11_O_9_S 1172.43571; found 1172.44299

##### ^1^H NMR (DMSO-d_6_) δ

8.98 (s, 1H), 8.55 (t, J=5.5 Hz, 0.1H rotamer), 8.49 (t, J=5.8 Hz, 1H), 8.11 (d, J=7.9 Hz, 0.1H rotamer), 7.64 (d, J=7.6 Hz, 1H), 7.56 (t, J=8.0 Hz, 1H), 7.49 (t, J=8.0 Hz, 1H), 7.42 (d, J=7.6 Hz, 1H), 7.35-7.40 (m, 0.1H rotamer), 7.29 (dd, J=9.1, 2.2 Hz, 1H), 7.20 (d, J=7.9 Hz, 0.1H rotamer), 7.16 (t, J=7.4 Hz, 0.1H rotamer), 7.03-7.07 (m, 1H), 6.97 (d, J=7.9 Hz, 1H), 5.17 (d, J=3.8 Hz, 1H), 5.04 (d, J=2.8 Hz, 0.1H rotamer), 4.69 (t, J=7.6 Hz, 0.1H rotamer), 4.60 (d, J=9.1 Hz, 1H), 4.52 (t, J=8.2 Hz, 1H), 4.45 (d, J=9.1 Hz, 0.1H rotamer), 4.32-4.38 (m, 1H), 4.31 (br d, J=6.3 Hz, 1H), 4.17-4.25 (m, 3H), 4.14 (br d, J=5.0 Hz, 0.1H rotamer), 4.11 (br d, J=5.0 Hz, 0.1H rotamer), 3.88 (br d, J=4.4 Hz, 2H), 3.76-3.85 (m, 4H), 3.66 (br d, J=3.8 Hz, 0.3H rotamer), 3.62-3.66 (range, 2H), 3.59-3.62 (range, 4H), 3.52-3.57 (range, 4H), 3.47-3.49 (m, 0.19H rotamer), 2.89-2.98 (range, 4H), 2.86 (br d, J=8.2 Hz, 4H), 2.46 (s, 5H), 2.38-2.43 (m, 4H), 2.21-2.29 (m, 0.1H rotamer), 2.09 (br dd, J=12.5, 7.7 Hz, 1H), 1.99-2.05 (m, 0.1H rotamer), 1.92 (ddd, J=12.9, 8.8, 4.4 Hz, 1H), 1.31-1.46 (m, 2H), 1.16-1.27 (m, 2H), 0.96 (s, 9H), 0.68 (br d, J=6.0 Hz, 2H), 0.41 (br s, 2H)

##### ^13^C NMR (DMSO-d_6_) δ

172.3, 169.4, 168.5 (d, CF=20 Hz), 168.1, 166.5, 166.2, 163.7, 158.9, 158.9, 156.3, 155.7, 155.5, 148.3, 134.6, 132.1, 131.7, 131.3, 128.2, 127.6, 125.9, 121.5, 121.0, 120.6, 118.9, 116.8, 115.9, 112.6, 111.4, 108.7, 108.7, 78.6 (d, CF=232 Hz), 70.5, 70.2, 69.4, 69.4, 68.8, 68.4, 59.3, 57.8, 57.2, 57.0, 53.5, 47.7, 46.9, 45.1, 44.4, 43.7, 39.4, 38.8, 38.4, 37.7, 36.5, 35.1, 34.9, 28.0, 27.5, 26.6, 16.5, 13.5(d, CF=10 Hz), 13.2(d, CF=11 Hz), 8.8, 8.7

##### HPLC

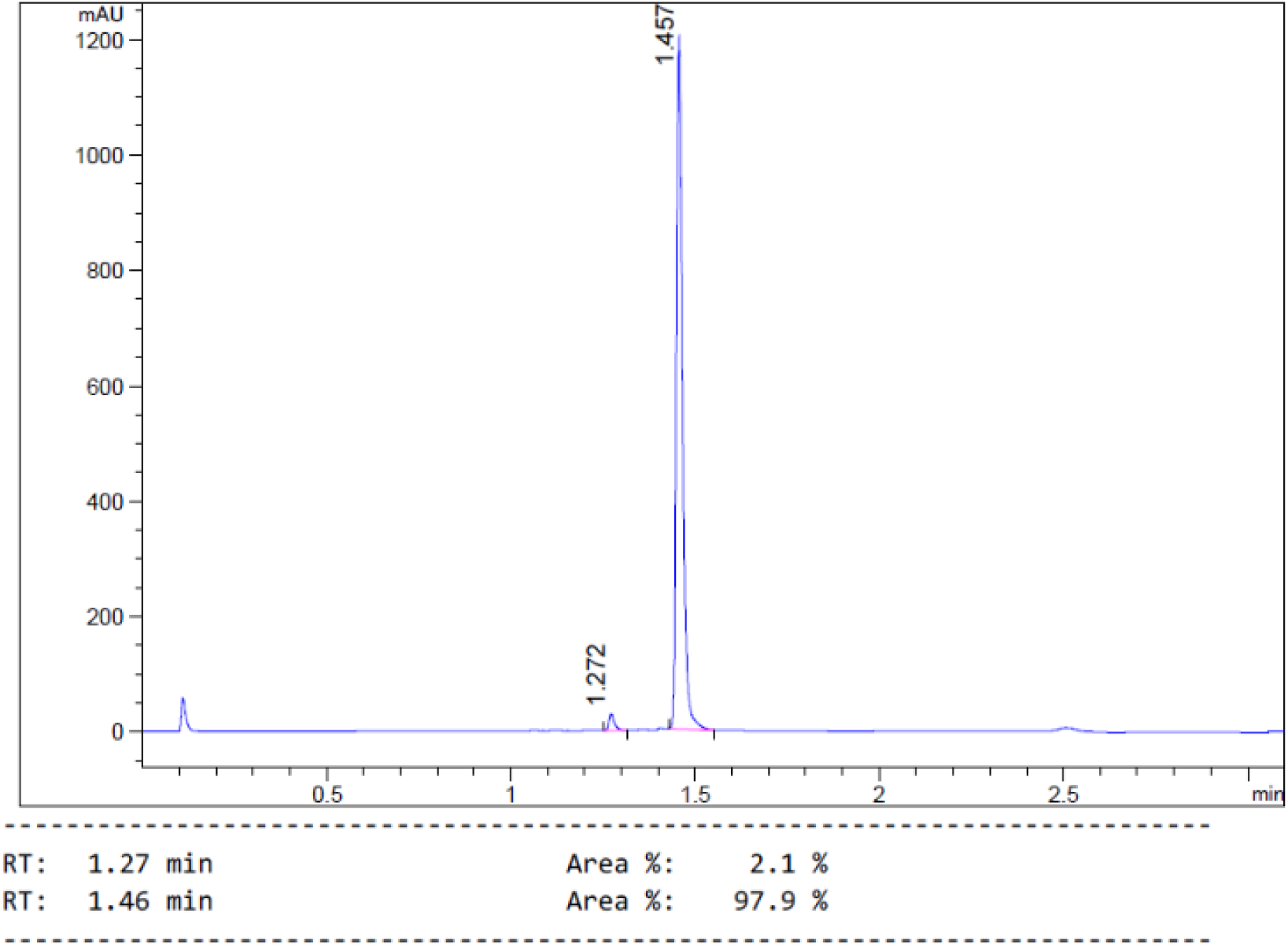

##### ^1^H and ^13^C NMR

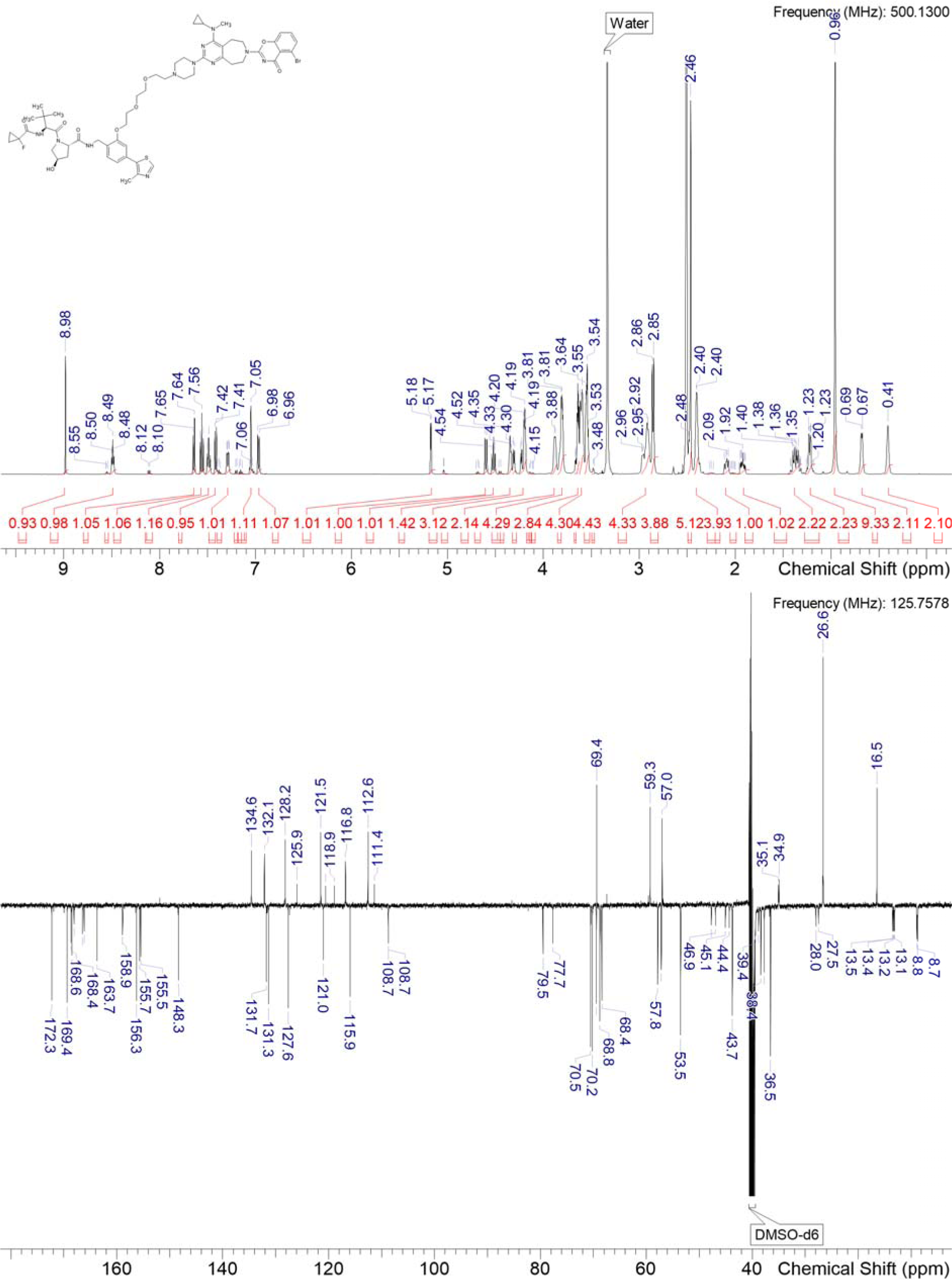

### P6

#### (2*S*,4*R*)-N-(2-(4-((4-(3-amino-6-(2-hydroxyphenyl)pyridazin-4-yl)piperazin-1-yl)methyl)-2-fluorophenethoxy)-4-(4-methylthiazol-5-yl)benzyl)-1-((*S*)-2-(1-fluorocyclopropane-1-carboxamido)-3,3-dimethylbutanoyl)-4-hydroxypyrrolidine-2-carboxamide P6

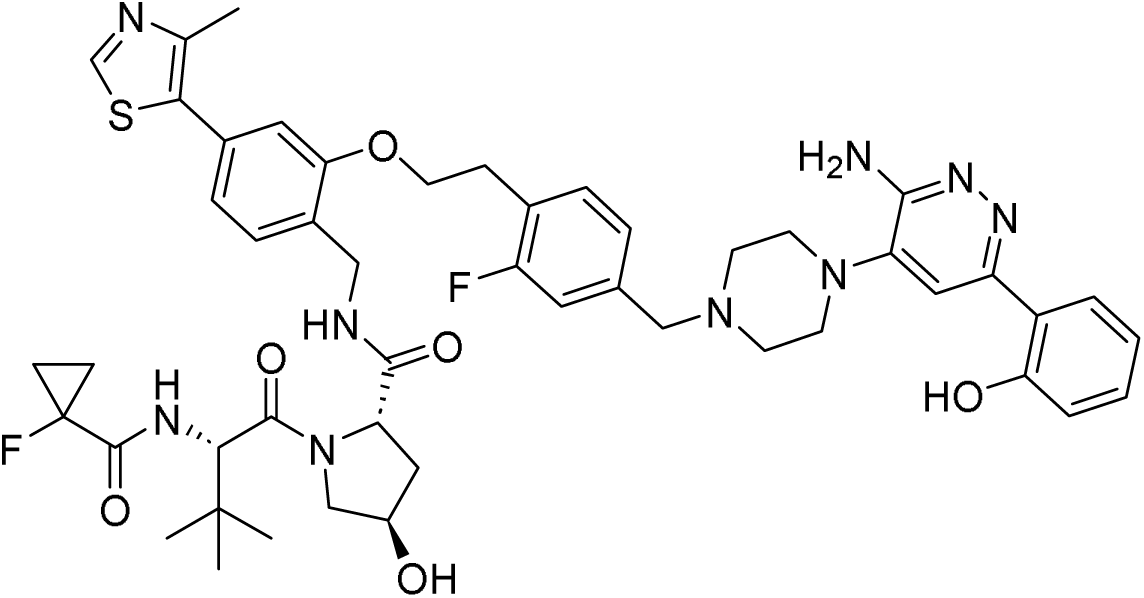

Synthesis described in [2].

##### HRMS (ESI+) *m*/*z*

[M+H]^+^ calcd for C_49_H_57_F_2_N_9_O_6_S 938.41392; found 938.4212

##### ^1^H NMR (DMSO-d_6_) δ

14.23 (br s, 1H), 8.98 (s, 1H), 8.55 (br t, J=5.5 Hz, 0.1H rotamer), 8.47 (br t, J=5.9 Hz, 1H), 7.91 (br d, J=7.9 Hz, 1H), 7.51 (s, 1H), 7.43 (br t, J=7.9 Hz, 1H), 7.40 (br d, J=7.7 Hz, 1H), 7.28 (br d, J=9.2 Hz, 1H), 7.24 (br t, J=7.7 Hz, 1H), 7.13-7.18 (range, 2H), 7.03 (s, 1H), 6.95 (br d, J=7.7 Hz, 1H), 6.85-6.92 (range, 2H), 6.24 (s, 2H), 5.17 (br d, J=2.8 Hz, 1H), 5.04 (br s, 0.1H rotamer), 4.64-4.71 (m, 0.1H rotamer), 4.59 (br d, J=9.0 Hz, 1H), 4.51 (br t, J=8.3 Hz, 1H), 4.46-4.48 (m, 0.1H rotamer), 4.32-4.37 (m, 1H), 4.25-4.32 (m, 2H), 4.22 (br dd, J=16.7, 6.4 Hz, 1H), 4.11 (br dd, J=16.6, 5.2 Hz, 1H), 4.05-4.07 (m, 0.1H rotamer), 3.58-3.67 (m, 2H), 3.57 (s, 2H), 3.05-3.21 (range, 6H), 2.74 (t, J=6.9 Hz, 0.1H rotamer), 2.57-2.67 (range, 4H), 2.45 (s, 3H), 2.06-2.14 (m, 1H), 1.87-1.96 (m, 1H), 1.30-1.43 (m, 2H), 1.16-1.28 (m, 2H), 0.94 (s, 9H), 0.82-0.88 (m, 0.1H rotamer)

##### ^13^C NMR (DMSO-d_6_) δ

172.3, 169.4, 168.5 (d, CF=20 Hz), 161.2 (d, CF=243 Hz), 159.0, 156.0, 155.1, 153.6, 151.9, 148.3, 140.9, 139.7 (d, CF=8 Hz), 131.9 (d, CF=5 Hz), 131.7, 131.3, 130.6, 128.1, 127.4, 126.6, 125.1, 124.2 (d, CF=17 Hz), 121.4, 118.9, 118.3, 117.8, 115.6 (d, CF=23 Hz), 112.1, 110.9, 78.6 (d, CF=233 Hz), 69.4, 67.8, 61.5, 59.3, 57.2, 57.0, 52.4, 49.0, 38.4, 37.6, 36.5, 28.6, 26.6, 16.4, 13.4 (d, CF=10 Hz), 13.2 (d, CF=10 Hz)

##### HPLC

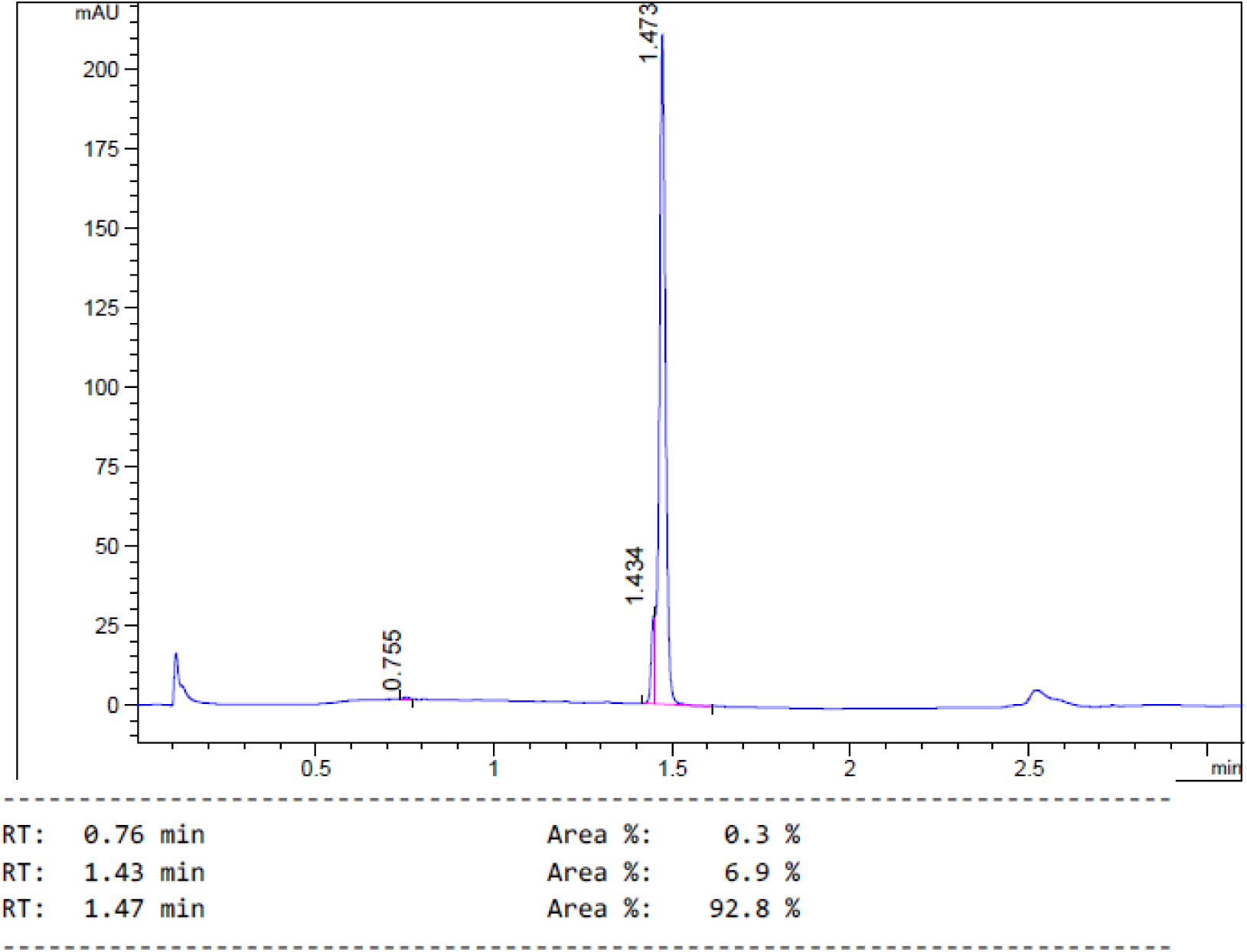

##### 1H and ^13^C NMR

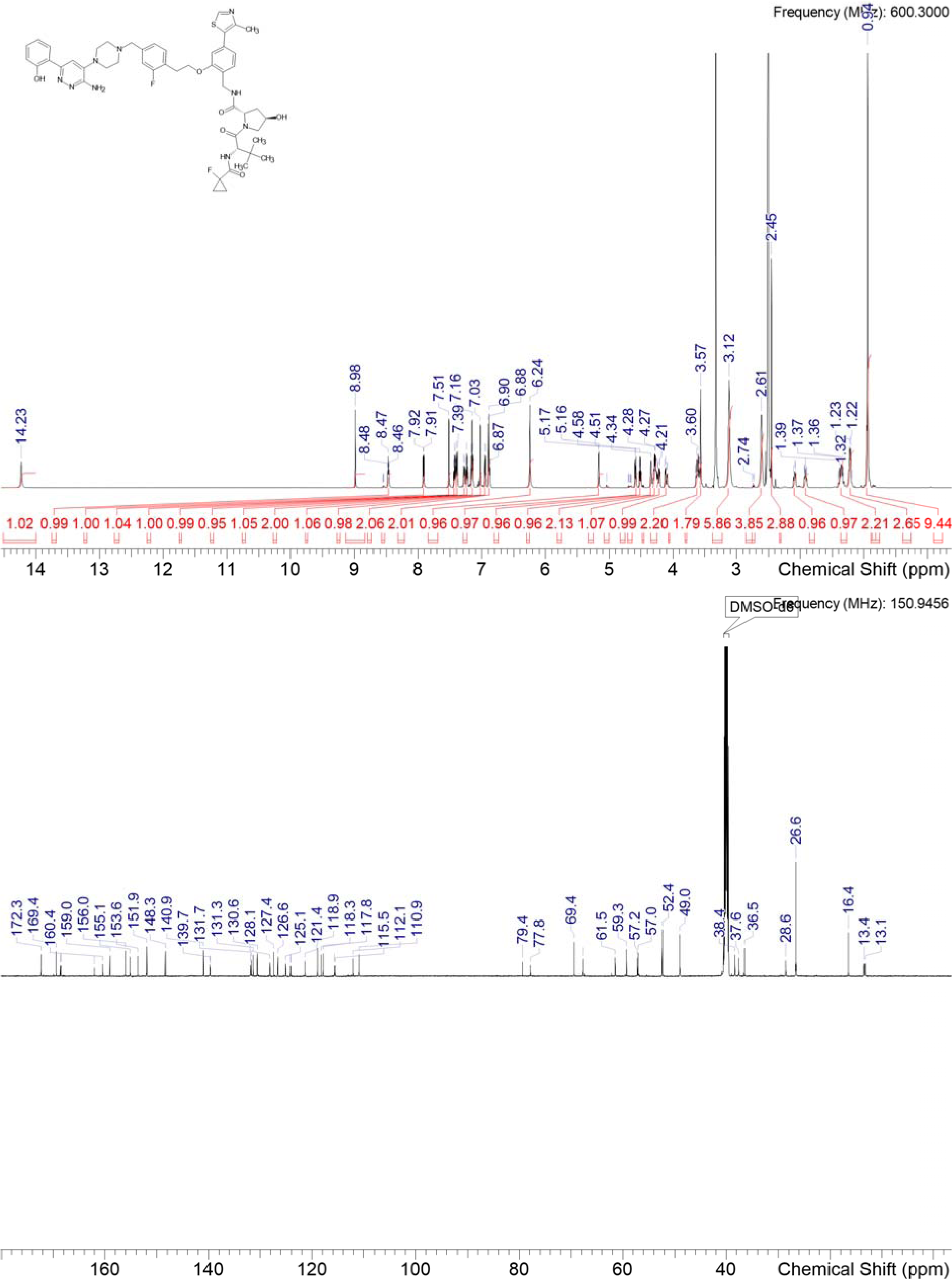

### P7

#### (2S,4R)-N-[(2-{4-[4-({4-[3-amino-6-(2-hydroxyphenyl)pyridazin-4-yl]piperazin-1-yl}methyl)phenoxy]butoxy}-4-(4-methyl-1,3-thiazol-5-yl)phenyl)methyl]-1-[(2*S*)-2-[(1-fluorocyclopropyl)formamido]-3,3-dimethylbutanoyl]-4-hydroxypyrrolidine-2-carboxamide P7

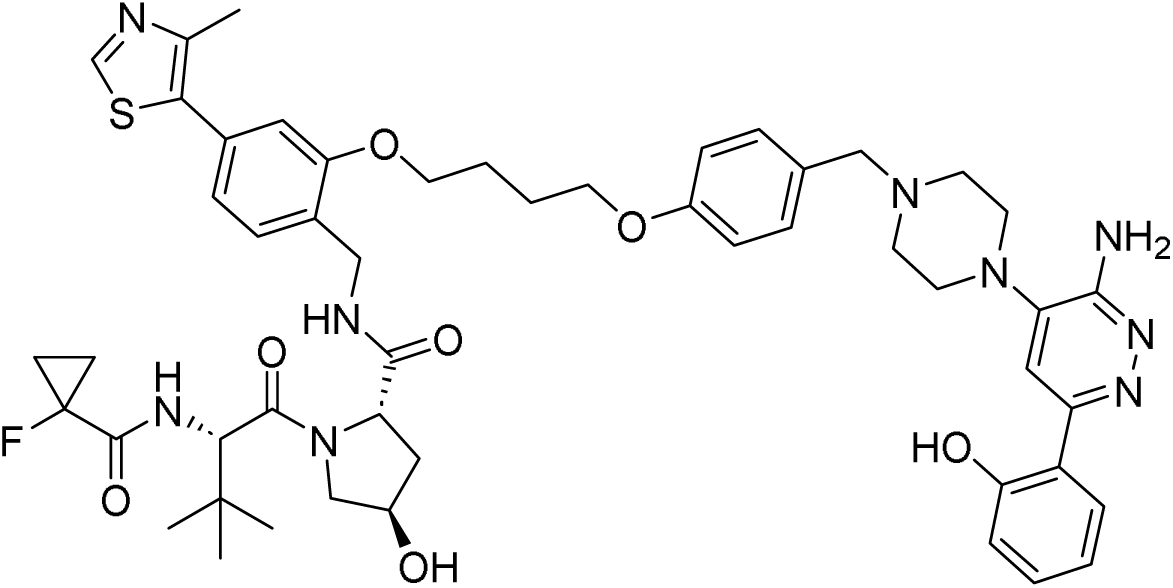

Synthesis described in [2].

##### HRMS (ESI+) *m*/*z*

[M+H] ^+^ calcd for C_51_H_62_FN_9_O_7_S 964.44951; found 964.45679

##### ^1^H NMR (DMSO-d_6_) δ

14.22 (br s, 1H), 8.98 (s, 1H), 8.57 (br t, J=5.6 Hz, 0.1H rotamer), 8.50 (t, J=5.9 Hz, 1H), 7.91 (dd, J=8.3, 1.3 Hz, 1H), 7.86 (d, J=8.8 Hz, 0.05H rotamer), 7.50 (s, 1H), 7.42 (d, J=7.7 Hz, 1H), 7.30 (dd, J=9.2, 2.2 Hz, 1H), 7.25 (d, J=1.5 Hz, 0.33H rotamer), 7.23 (d, J=8.4 Hz, 3H), 7.20 (d, J=7.7 Hz, 0.1H rotamer), 7.13 (d, J=8.6 Hz, 0.1H rotamer), 7.03 (br s, 0.1H rotamer), 7.02 (d, J=1.1 Hz, 1H), 6.96 (dd, J=7.8, 1.2 Hz, 1H), 6.83-6.93 (range, 4H), 6.22 (s, 2H), 5.19 (d, J=3.5 Hz, 1H), 5.06 (d, J=2.8 Hz, 0.1H rotamer), 4.68 (t, J=7.4 Hz, 0.1H rotamer), 4.60 (d, J=9.4 Hz, 1H), 4.52 (t, J=8.3 Hz, 1H), 4.46 (br t, J=9.4 Hz, 0.1H rotamer), 4.36 (br s, 1H), 4.31 (dd, J=16.3, 6.1 Hz, 1H), 4.22 (dd, J=16.3, 5.5 Hz, 1H), 4.13 (br t, J=4.1 Hz, 2H), 4.05 (br t, J=4.1 Hz, 2H), 3.65 (dd, J=10.8, 3.8 Hz, 1H), 3.61 (br d, J=3.6 Hz, 1H), 3.49 (s, 2H), 3.04-3.16 (m, 4H), 2.55-2.61 (m, 4H), 2.46 (s, 3H), 2.25 (br t, J=9.9 Hz, 0.1H rotamer), 2.10 (br dd, J=12.5, 7.9 Hz, 1H), 2.02 (ddd, J=12.4, 7.0, 5.0 Hz, 0.1H rotamer), 1.87-1.97 (range, 4H), 1.28-1.45 (m, 2H), 1.14-1.27 (m, 2H), 0.96 (s, 9H)

##### ^13^C NMR (DMSO-d_6_) δ

172.3, 169.4, 168.6 (d, CF=20 Hz), 159.0, 158.2, 156.3, 155.1, 153.6, 151.9, 148.3, 140.9, 131.8, 131.4, 130.6, 130.2, 128.2, 127.4, 126.6, 121.2, 119.0, 118.3, 117.8, 114.6, 112.1, 110.8, 78.6 (d, CF=233 Hz), 69.4, 67.6, 61.9, 59.3, 57.1, 57.0, 52.3, 49.0, 38.3, 37.7, 36.5, 26.6, 26.0, 16.4, 13.4 (d, CF=10 Hz), 13.2 (d, CF=10 Hz)

##### HPLC

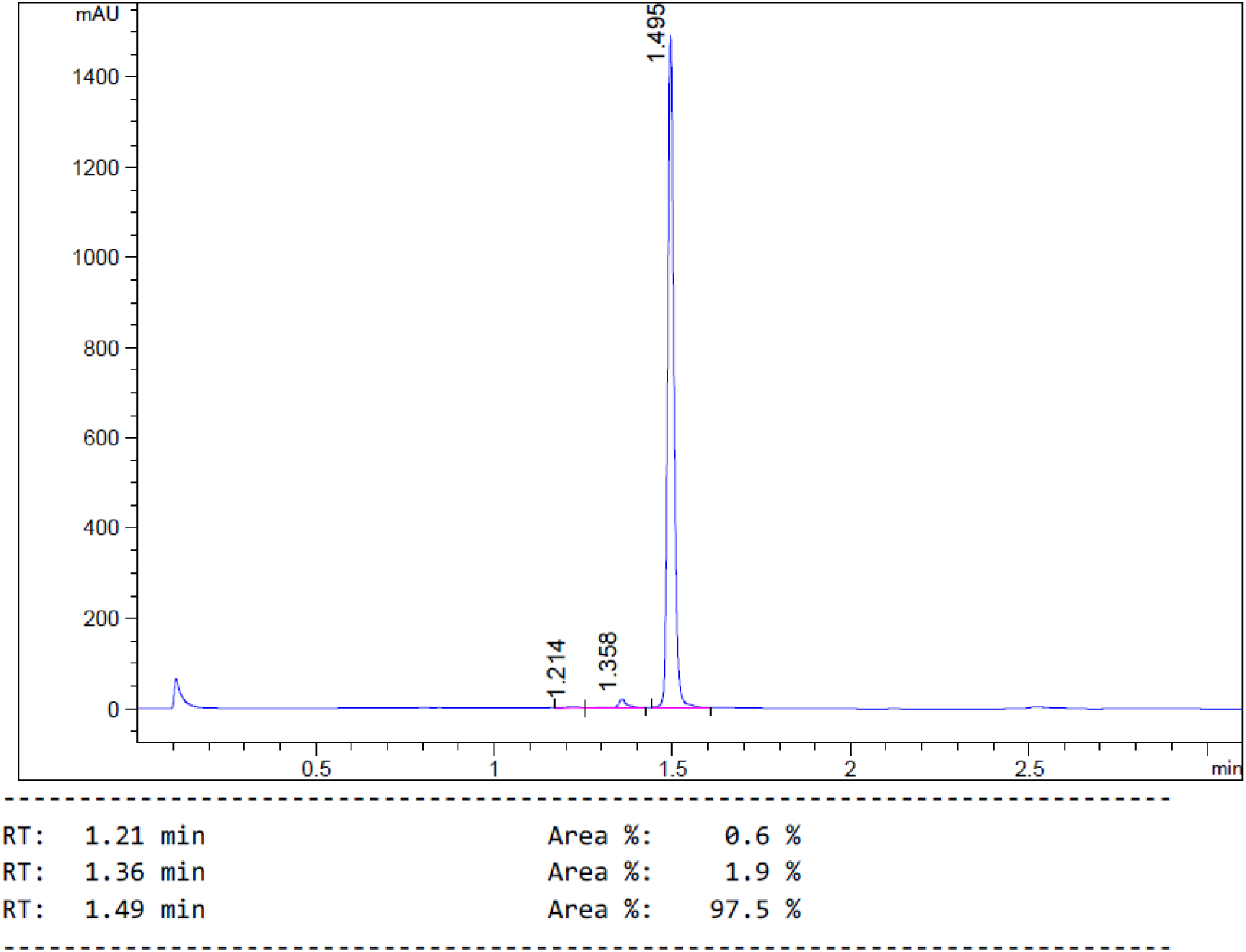

##### ^1^H and ^13^C NMR

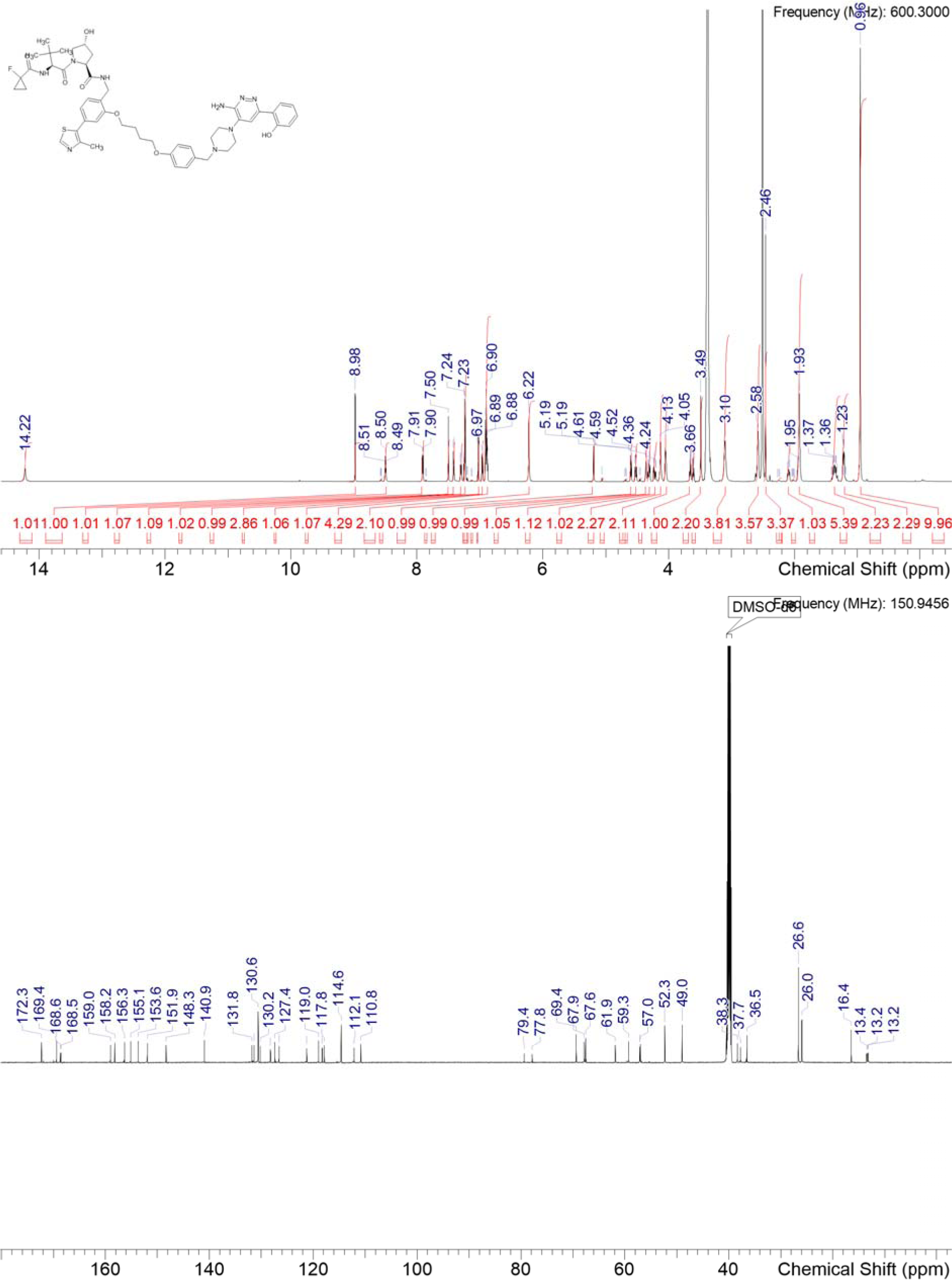

### P9

**Scheme S6:**
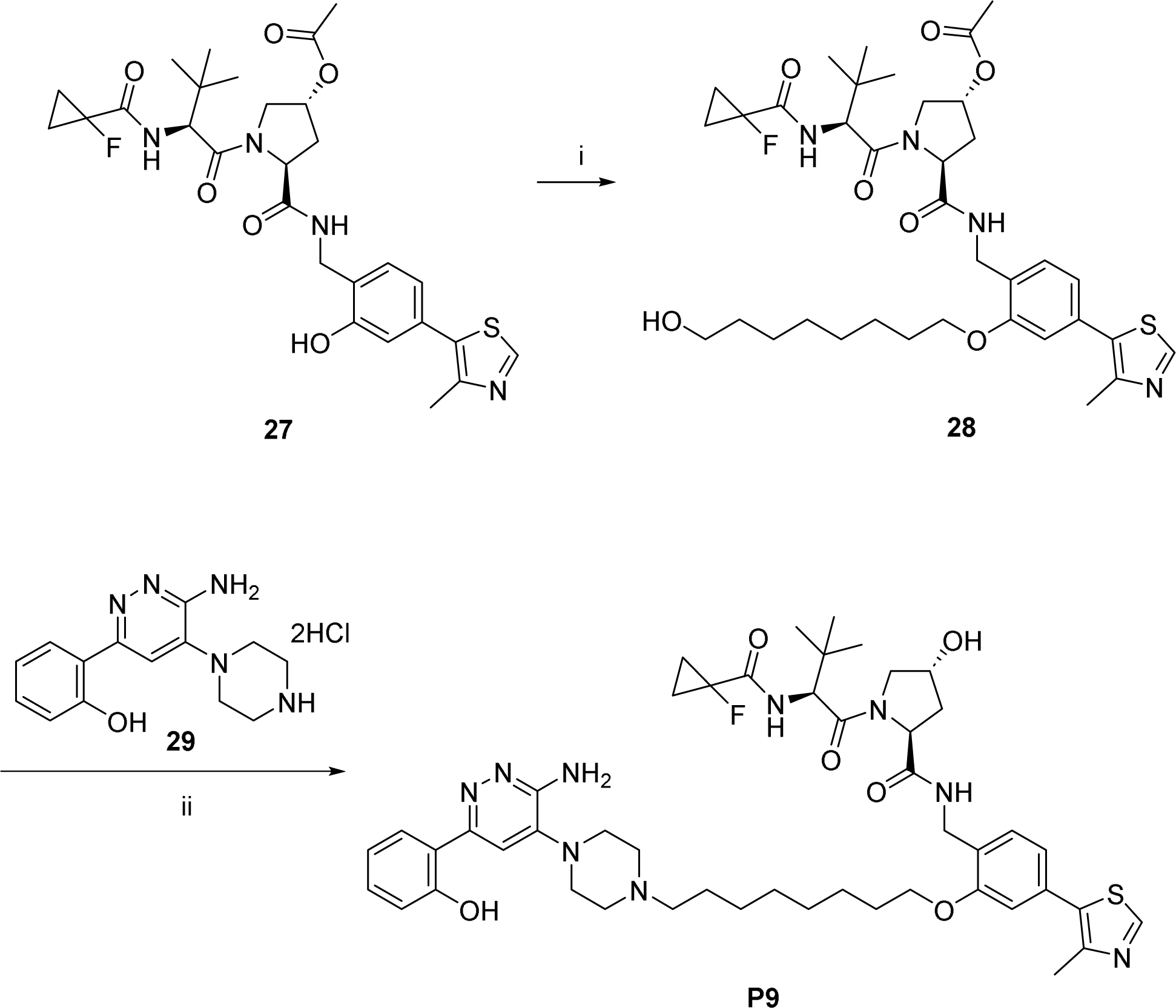
Synthesis of P9: i) 8-bromo octanol-1-ol, K_2_CO_3_, DMF, 90°C; ii) COCl_2_, DMSO, TEA, DCM, -78°C; *then* **29**, TEA, DCE, STAB, RT; *then* NaOH, EtOH, 50°C.

Starting material syntheses are reported in the literature:

(2*S*,4*R*)-1-((*S*)-2-(1-fluorocyclopropane-1-carboxamido)-3,3-dimethylbutanoyl)-4-hydroxy-*N*-(2-hydroxy-4-(4-methylthiazol-5-yl)benzyl)pyrrolidine-2-carboxamide was synthesized as discussed in [3],

(3*R*,5*S*)-1-((*S*)-2-(1-fluorocyclopropane-1-carboxamido)-3,3-dimethylbutanoyl)-5-((2-hydroxy-4-(4-methylthiazol-5-yl)benzyl)carbamoyl)pyrrolidin-3-yl acetate was synthesised from (2*S*,4*R*)-1-((*S*)-2-(1-fluorocyclopropane-1-carboxamido)-3,3-dimethylbutanoyl)-4-hydroxy-*N*-(2-hydroxy-4-(4-methylthiazol-5-yl)benzyl)pyrrolidine-2-carboxamide as in [2].

#### (3*R*,5*S*)-1-((S)-2-(1-fluorocyclopropane-1-carboxamido)-3,3-dimethylbutanoyl)-5-((2-((8-hydroxyoctyl)oxy)-4-(4-methylthiazol-5-yl)benzyl)carbamoyl)pyrrolidin-3-yl acetate 28

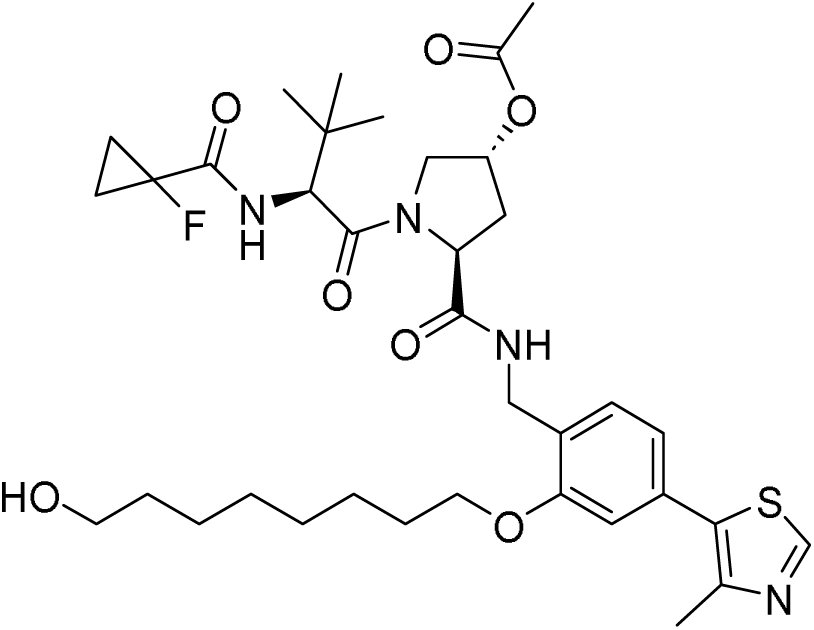

A mixture of (3*R*,5*S*)-1-((*S*)-2-(1-fluorocyclopropane-1-carboxamido)-3,3-dimethylbutanoyl)-5-((2-hydroxy-4-(4-methylthiazol-5-yl)benzyl)carbamoyl)pyrrolidin-3-yl acetate **27** (80 mg, 0.139 mmol) and K_2_CO_3_ (22 mg, 0.159 mmol) in DMF (2 mL) was pre-stirred at 90°C for 45 minutes before addition of 8-bromo-octan-1-ol (2 µL, 0.146 mmol). The resulting reaction mixture was stirred at 90°C for 5 hours. The reaction mixture was cooled to room temperature, diluted with H_2_O and DCM, and passed through a phase separator. The filtrate was concentrated under reduced pressure. The remaining residue was passed through a silica column (0-10% MeOH in CH_2_Cl_2_) and the desired fractions concentrated under reduced pressure to afford **28** (79.7 mg) containing octan-1-ol related impurities which was taken directly into the next reaction without further purification.

#### (2*S*,4*R*)-*N*-(2-((8-(4-(3-amino-6-(2-hydroxyphenyl)pyridazin-4-yl)piperazin-1-yl)octyl)oxy)-4-(4-methylthiazol-5-yl)benzyl)-1-((*S*)-2-(1-fluorocyclopropane-1-carboxamido)-3,3-dimethylbutanoyl)-4-hydroxypyrrolidine-2-carboxamide P9

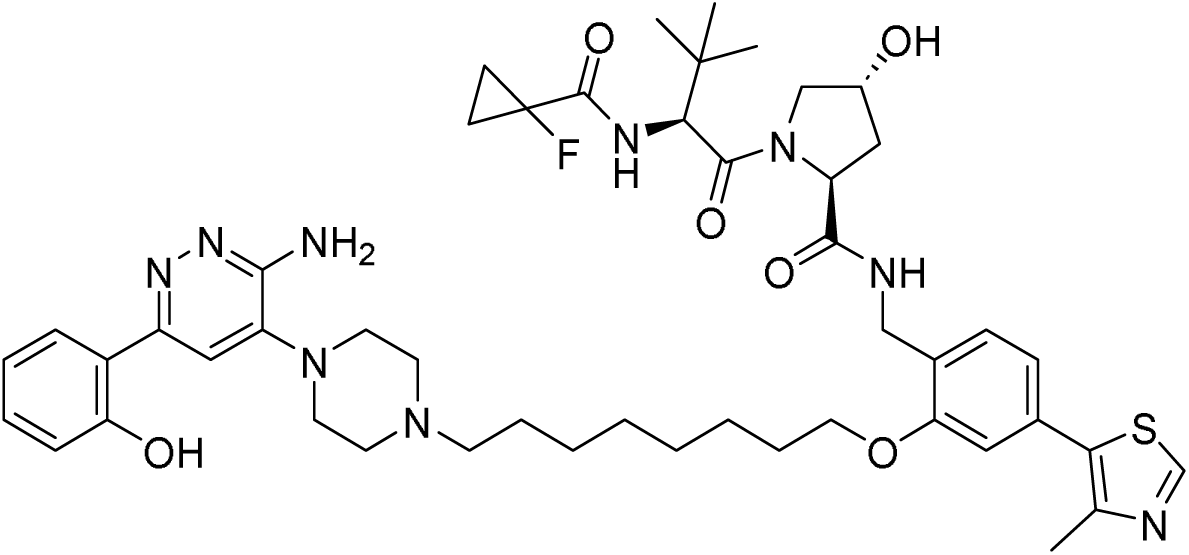

DMSO (15 µL, 0.211 mmol) was added to a solution of oxalyl chloride (15 µL, 0.175 mmol) in CH_2_Cl_2_ (1 mL) at −78 °C. The resulting reaction mixture was stirred at −78 °C for 15 min, after which (3*R*,5*S*)-1-((*S*)-2-(1-fluorocyclopropane-1-carboxamido)-3,3-dimethylbutanoyl)-5-((2-((8-hydroxyoctyl)oxy)-4-(4-methylthiazol-5-yl)benzyl)carbamoyl)pyrrolidin-3-yl acetate **28** (79.7 mg) in CH_2_Cl_2_ (1 mL) was added dropwise. The reaction mixture was stirred at −78 °C for 1 hour, after which triethylamine (76 µL, 0.525 mmol) was added. The reaction mixture was allowed to warm up to room temperature and stirred at ambient temperature for 1 hour. The reaction mixture was then concentrated under reduced pressure. The crude residue was used directly without purification.

The crude residue was added to a solution of 2-(6-amino-5-(piperazin-1-yl)pyridazin-3-yl)phenol dihydrochloride **29** (37 mg, 0.107 mmol) and triethylamine (500 µL, 0.526 mmol) in DCE (3 mL) and DMSO (0.6 mL). STAB (116 mg, 0.547 mmol) was added shortly after, followed by magnesium sulfate. The reaction mixture was stirred at room temperature overnight. The reaction mixture was filtered and concentrated under reduced pressure. The residue was purified by reverse phase chromatography under acidic conditions (0-60% CH_3_CN in 0.1% aq. HCO_2_H). Desired fractions were concentrated under reduced pressure. The resulting residue was dissolved in EtOH (3.000 mL) and aq. 2M NaOH (3 mL, 6 mmol) and stirred at 50 °C for 1 hour. The reaction mixture was neutralized to pH 6 with aq. 1M HCl and concentrated under reduced pressure. The residue was purified by reverse phase chromatography on acidic conditions (0-60% CH_3_CN in 0.1% aq. HCO_2_H) and the desired fractions concentrated under reduced pressure to afford **P9** (46.5 mg, 0.051 mmol, 37 % over 3 steps from (3*R*,5*S*)-1-((*S*)-2-(1-fluorocyclopropane-1-carboxamido)-3,3-dimethylbutanoyl)-5-((2-hydroxy-4-(4-methylthiazol-5-yl)benzyl)carbamoyl)pyrrolidin-3-yl acetate).

^1^H NMR (400 MHz, CD_2_Cl_2_) δ 8.69 (s, 1H), 7.65 (d, *J* = 7.9 Hz, 1H), 7.39 (s, 1H), 7.33 (d, *J* = 7.7 Hz, 1H), 7.28 (t, *J* = 7.7 Hz, 1H), 7.20 (t, *J* = 5.7 Hz, 1H), 7.09 – 7.01 (m, 1H), 6.97 (t, *J* = 7.9 Hz, 2H), 6.94 – 6.87 (m, 2H), 5.30 (s, 2H), 4.67 (t, *J* = 7.7 Hz, 1H), 4.59 (d, *J* = 8.9 Hz, 1H), 4.55 – 4.35 (m, 3H), 4.04 (t, *J* = 6.2 Hz, 2H), 3.89 (d, *J* = 11.0 Hz, 1H), 3.64 (dd, *J* = 11.1, 3.9 Hz, 1H), 3.36 – 3.26 (m, 4H), 3.02 (s, 3H), 2.73 (dd, *J* = 9.4, 5.2 Hz, 2H), 2.51 (s, 3H), 2.48 – 2.39 (m, 1H), 2.13 – 2.04 (m, 1H), 1.91 – 1.79 (m, 2H), 1.66 (s, 2H), 1.53 (s, 2H), 1.46 – 1.21 (m, 11H), 0.94 (s, 9H); LC-MS: t_Ret._ = 1.33 min; MS (M+H)^+^ = 914

### P10

**Scheme S7:**
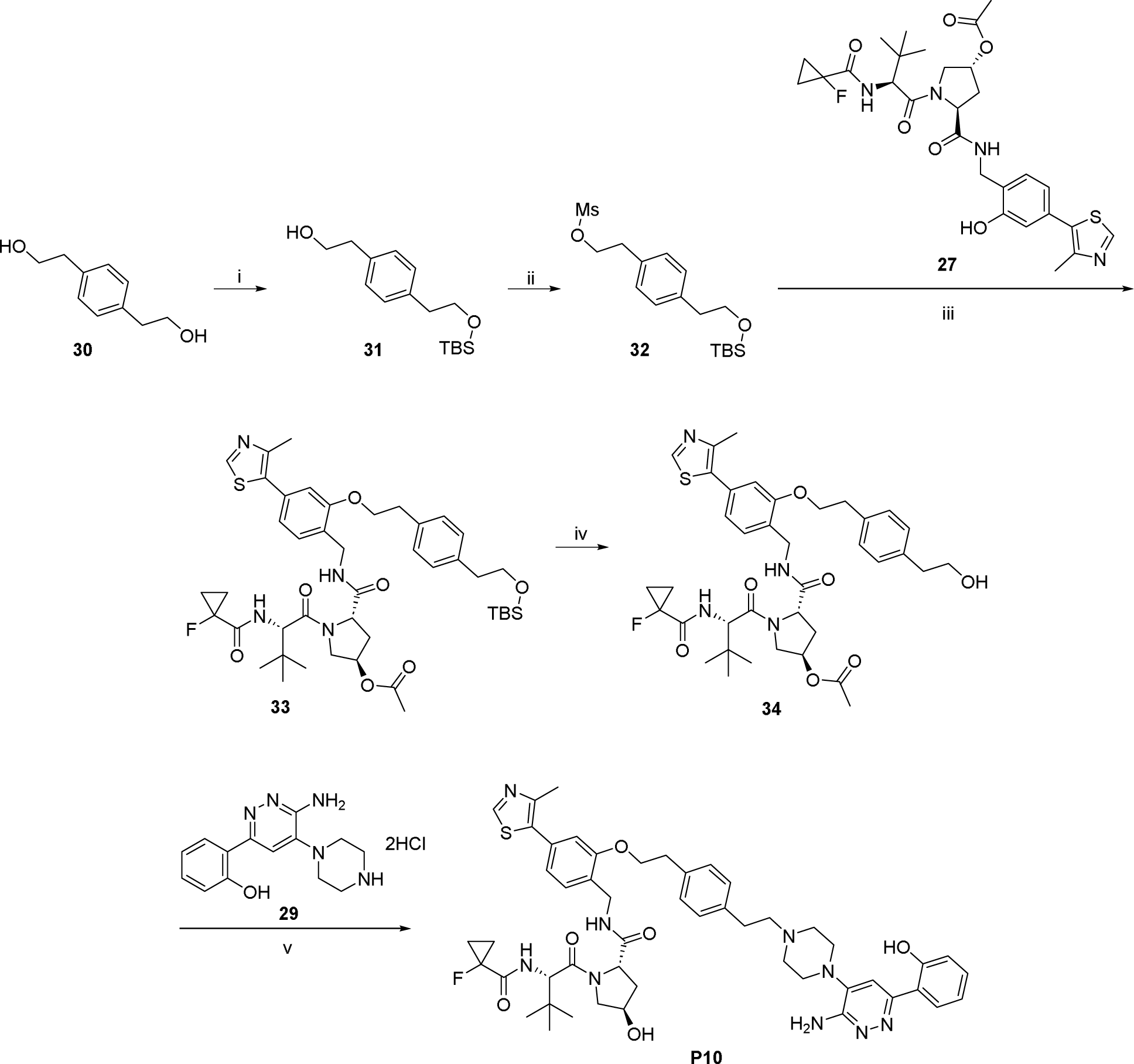
Synthesis of P10: i) TBSCl, imidazole, DMF, 50°C; ii) MsCl, TEA, DCM, 0°C; iii) **27**, K_2_CO_3_, DMF, 70°C; iv) TBAF, THF, RT; v) COCl_2_, DMSO, TEA, DCM, - 78°C; *then* **29**, TEA, DCE, STAB, RT; *then* NaOH, EtOH, 50°C.

#### 2-(4-(2-((*tert*-butyldimethylsilyl)oxy)ethyl)phenyl)ethan-1-ol 31

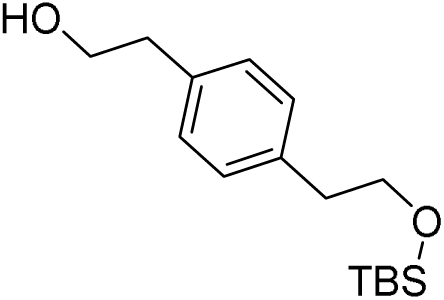

*tert*-Butyl-chloro-dimethyl-silane (1.50 g, 0.010 mol) was added to a mixture of 2,2’-(1,4-phenylene)bis(ethan-1-ol) **30** (1.60 g, 0.010 mol) and imidazole (1.30 g, 0.019 mol) in DMF (20 mL). The resulting reaction mixture was stirred at 50°C overnight. The reaction mixture was cooled to room temperature, diluted with H_2_O, and extracted with EtOAc. The combined organics were washed with brine, dried over MgSO_4_, filtered, and concentrated under reduced pressure. The residue was purified by column chromatography on silica gel (0-50% EtOAc in heptane) and the desired fractions concentrated under reduced pressure to afford **31** (1.00 g, 3.56 mmol, 37%).

#### 4-(2-((*tert*-butyldimethylsilyl)oxy)ethyl)phenethyl methanesulfonate 32

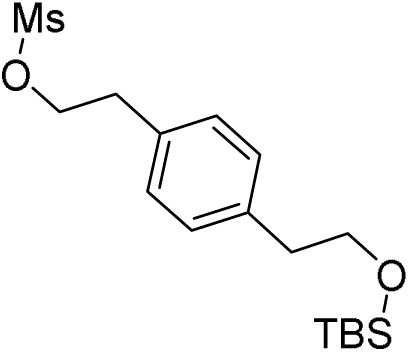

Methanesulfonyl chloride (400 µL, 5.24 mmol) was added to a solution of **31** (775 mg, 2.76 mmol) and triethylamine (2.0 mL, 13.8 mmol) in CH_2_Cl_2_ (8 mL) and THF (8 mL) at 0°C. The reaction mixture was stirred at 0°C for 120 minutes. Sat. aq. NaHCO_3_ was added, and the reaction mixture filtered through a phase separator. The filtrate was concentrated under reduced pressure and the residue purified by column chromatography on silica gel (0-50% EtOAc in Heptane). Desired fractions were concentrated under reduced pressure to afford **32** (161 mg, 0.449 mmol, 16%).

#### (3*R*,5*S*)-5-((2-(4-(2-((*tert*-butyldimethylsilyl)oxy)ethyl)phenethoxy)-4-(4-methylthiazol-5-yl)benzyl)carbamoyl)-1-((*S*)-2-(1-fluorocyclopropane-1-carboxamido)-3,3-dimethylbutanoyl)pyrrolidin-3-yl acetate 33

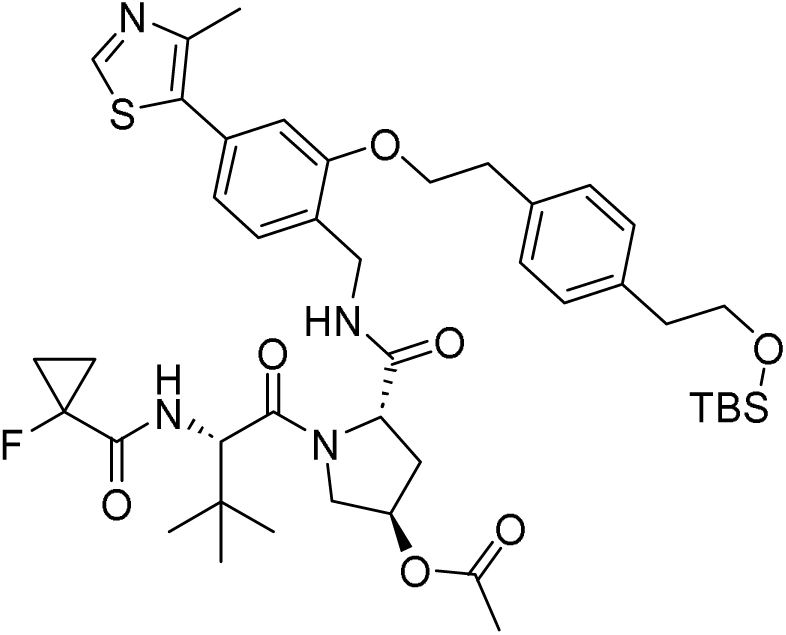

**32** (161 mg, 0.449 mmol) was added to a solution of **27** (258 mg, 0.449 mmol) and K_2_CO_3_ (150 mg, 1.08 mmol) in DMF (5 mL). The resulting reaction mixture was stirred at 70°C overnight. The reaction mixture was diluted with CH_2_Cl_2_, quenched with water and passed through a phase separator. The organic phase was concentrated under reduced pressure and the residue purified by column chromatography on silica gel (0-10% MeOH in CH_2_Cl_2_) to afford **33** (237 mg, 0.282 mmol, 63%).

#### (3*R*,5*S*)-1-((*S*)-2-(1-fluorocyclopropane-1-carboxamido)-3,3-dimethylbutanoyl)-5-((2-(4-(2-hydroxyethyl)phenethoxy)-4-(4-methylthiazol-5-yl)benzyl)carbamoyl)pyrrolidin-3-yl acetate 34

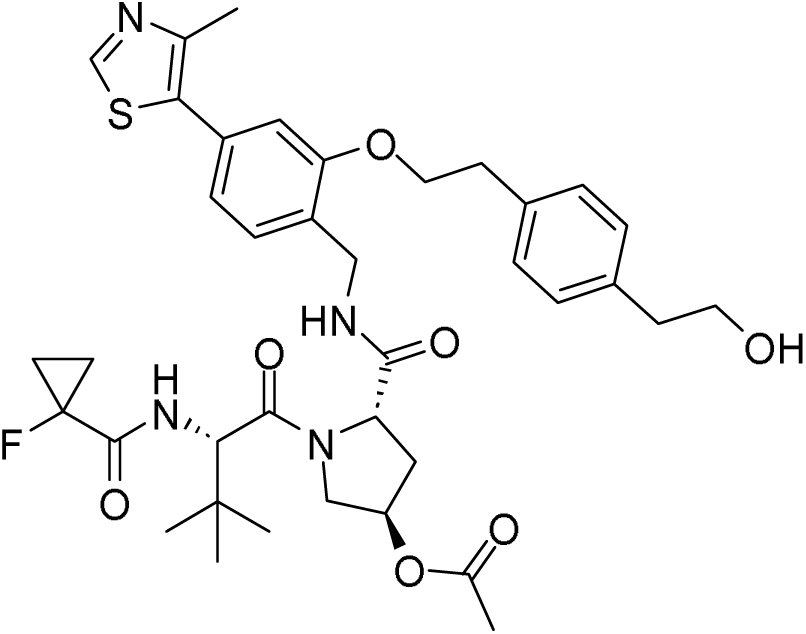

**33** (236 mg, 0.283 mmol) was dissolved in THF (5 mL) and TBAF (300 µL, 1M in THF, 0.3 mmol) was added. The reaction mixture was stirred at room temperature for 2 hours before being diluted with CH_2_Cl_2_, dried over MgSO_4_, filtered, and concentrated under reduced pressure. The residue was purified by column chromatography on silica gel (0-5% MeOH in CH_2_Cl_2_) and the desired fractions concentrated under reduced pressure to afford **34** (172 mg, 0.237 mmol, 84%).

#### (2*S*,4*R*)-*N*-(2-(4-(2-(4-(3-amino-6-(2-hydroxyphenyl)pyridazin-4-yl)piperazin-1-yl)ethyl)phenethoxy)-4-(4-methylthiazol-5-yl)benzyl)-1-((*S*)-2-(1-fluorocyclopropane-1-carboxamido)-3,3-dimethylbutanoyl)-4-hydroxypyrrolidine-2-carboxamide P10

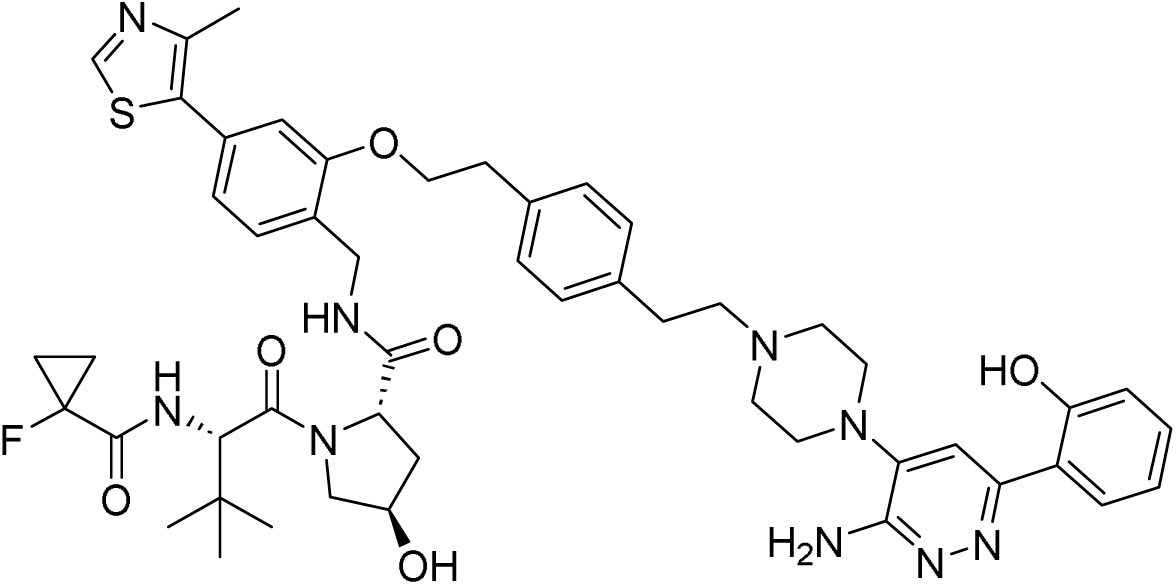

DMSO (34 µL, 0.478 mmol) was added to a solution of oxalyl chloride (30 µL, 0.350 mmol) in CH_2_Cl_2_ (2 mL) at −78°C. The resulting reaction mixture was stirred at −78°C for 15 min before dropwise addition of a solution of **34** (171 mg, 0.237 mmol) in CH_2_Cl_2_ (2 mL). The reaction mixture is stirred at −78°C for 1 hour, after which triethylamine (137 µL, 0.946 mmol) was added. The reaction mixture is allowed to warm up to room temperature and is stirred at the same temperature for 1 hour. The reaction mixture was warmed to room temperature and concentrated under reduced pressure. The crude residue was used directly without purification.

The crude residue was added to a solution of **29** (81 mg, 0.237 mmol) and triethylamine (1.03 mL, 7.11 mmol) in DCE (4 mL) and DMSO (1 mL). STAB (251 mg, 1,186 mmol) was added, followed by magnesium sulfate. The reaction mixture was stirred at room temperature overnight before being filtered and concentrated under reduced pressure. The residue was purified by reverse phase chromatography under acidic conditions (0-60% CH_3_CN in 0.1% aq. HCO_2_H). Desired fractions were concentrated under reduced pressure and the residue dissolved in ethanol (3 mL) and 2M aq. NaOH (3 mL, 6 mmol). The resulting reaction mixture was stirred at 50°C for 1 hour. The reaction mixture was neutralized to pH 6 with 1M aq. HCl and concentrated under reduced pressure. The residue was purified by preparative HPLC under basic conditions (5-95% CH_3_CN in 0.1% aq. NH_3_) to afford **P10** (41.6 mg, 0.044 mmol, 19%).

##### HRMS (ESI+) *m*/*z*

[M+H] ^+^ calcd for C_50_H_60_FN_9_O_6_S 934.43925; found 934.44653

##### ^1^H NMR (DMSO-d_6_) δ

14.24 (br s, 1H), 8.98 (s, 1H), 8.57 (br t, J=5.6 Hz, 0.1H rotamer), 8.53 (s, 0.1H rotamer), 8.47 (t, J=5.9 Hz, 1H), 7.92 (dd, J=8.3, 1.3 Hz, 1H), 7.51 (s, 1H), 7.39 (d, J=7.9 Hz, 1H), 7.23-7.32 (range, 4H), 7.21 (d, J=7.9 Hz, 2H), 7.15 (d, J=8.1 Hz, 0.3H rotamer), 7.02 (br s, 0.1H rotamer), 7.01 (d, J=1.1 Hz, 1H), 6.95 (dd, J=7.8, 1.2 Hz, 1H), 6.85-6.92 (range, 2H), 6.23 (s, 2H), 5.19 (br s, 1H), 5.06 (br s, 0.1H rotamer), 4.70 (t, J=7.4 Hz, 0.1H rotamer), 4.60 (d, J=9.2 Hz, 1H), 4.52 (t, J=8.3 Hz, 1H), 4.48 (br d, J=8.8 Hz, 0.1H rotamer), 4.36 (br s, 1H), 4.30-4.34 (m, 0.1H rotamer), 4.20-4.29 (range, 3H), 4.15 (br dt, J=16.5, 5.5 Hz, 1H), 4.09 (br dd, J=16.2, 4.9 Hz, 0.1H rotamer), 3.65 (dd, J=10.8, 3.8 Hz, 1H), 3.60 (d, J=6.6 Hz, 1H), 3.46-3.52 (m, 0.1H rotamer), 3.08-3.19 (range, 4H), 3.05 (br t, J=6.2 Hz, 2H), 2.76 (br t, J=7.6 Hz, 2H), 2.64-2.72 (range, 4H), 2.56-2.64 (m, 2H), 2.45 (s, 3H), 2.20-2.29 (m, 0.1H rotamer), 2.10 (br dd, J=12.6, 7.8 Hz, 1H), 2.00-2.06 (m, 0.1H rotamer), 1.93 (ddd, J=12.9, 8.8, 4.4 Hz, 1H), 1.31-1.45 (m, 2H), 1.16-1.28 (m, 2H), 0.96 (s, 9H)

##### ^13^C NMR (DMSO-d_6_) δ

172.3, 169.4, 168.6, 159.0, 156.2, 155.1, 153.7, 151.9, 148.3, 140.9, 138.8, 136.5, 131.7, 131.3, 130.6, 129.5, 129.1, 128.1, 127.3, 126.6, 121.3, 118.9, 118.3, 117.8, 112.1, 110.8, 78.6, 69.4, 69.1, 60.1, 59.3, 57.2, 57.0, 52.6, 49.0, 40.5, 38.4, 37.7, 36.5, 35.2, 32.8, 26.7, 26.6, 16.4, 13.5, 13.2

##### HPLC

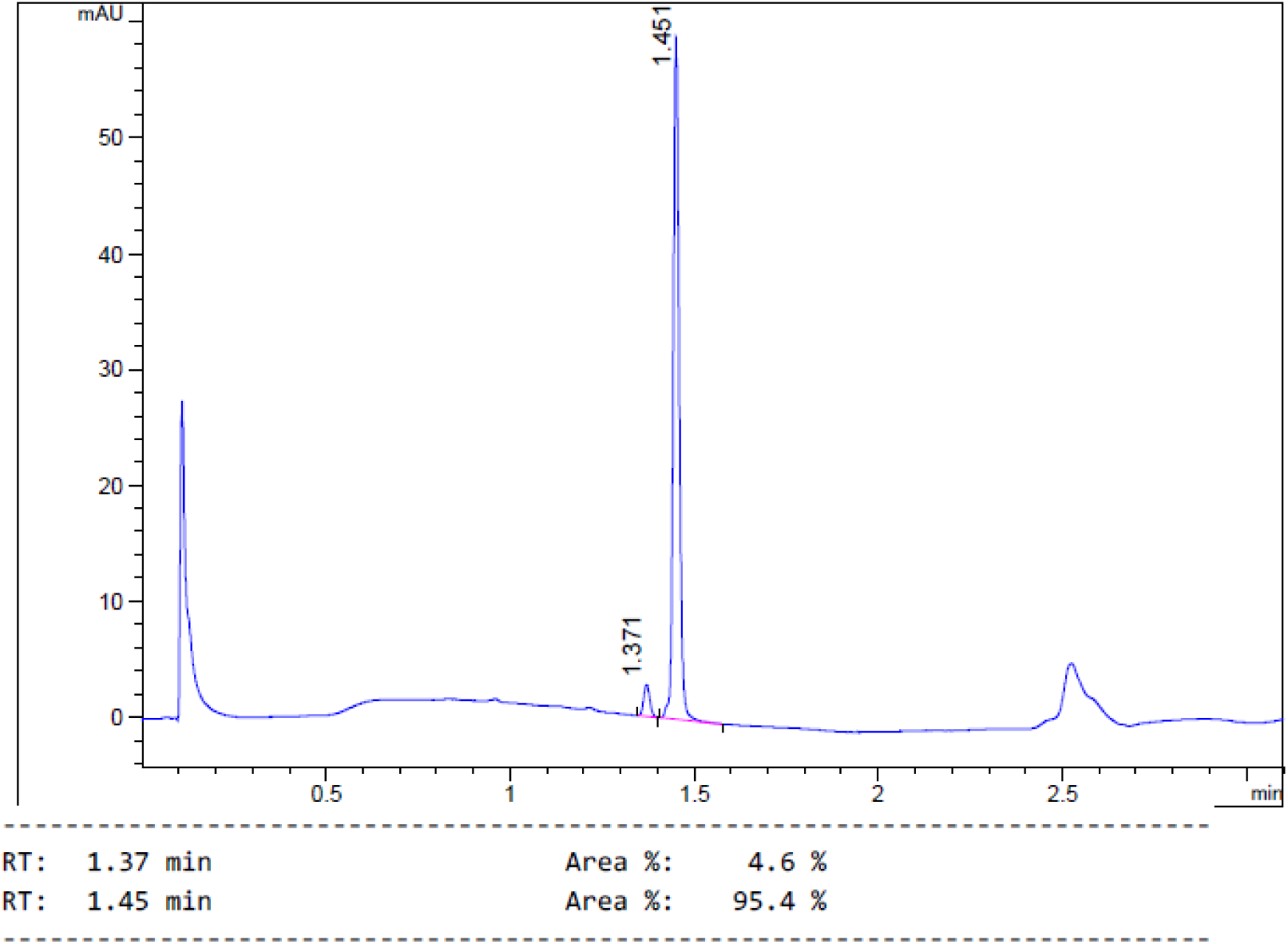

##### 1H and ^13^C NMR

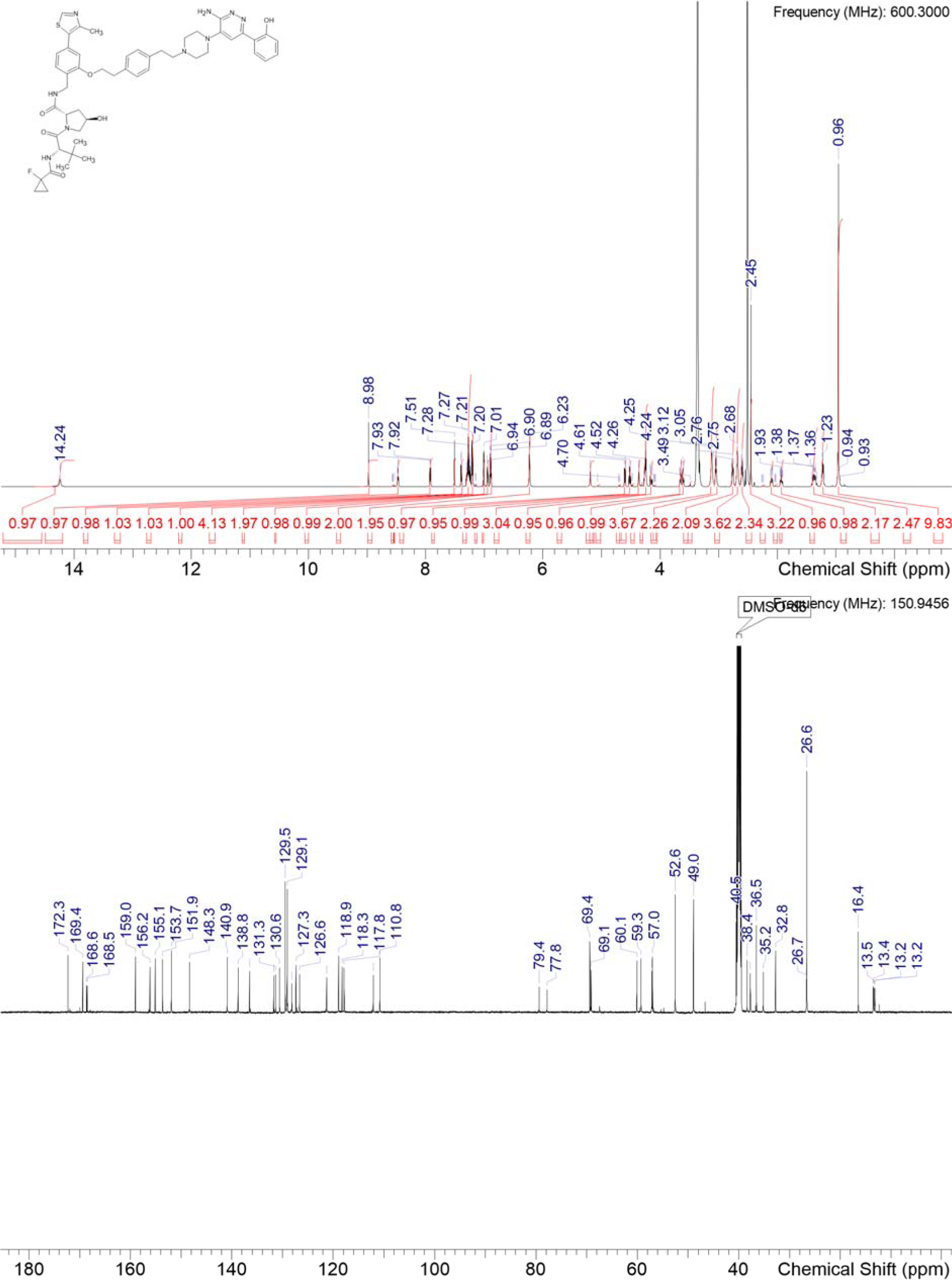

### Synthesis of P11

**Scheme S8:**
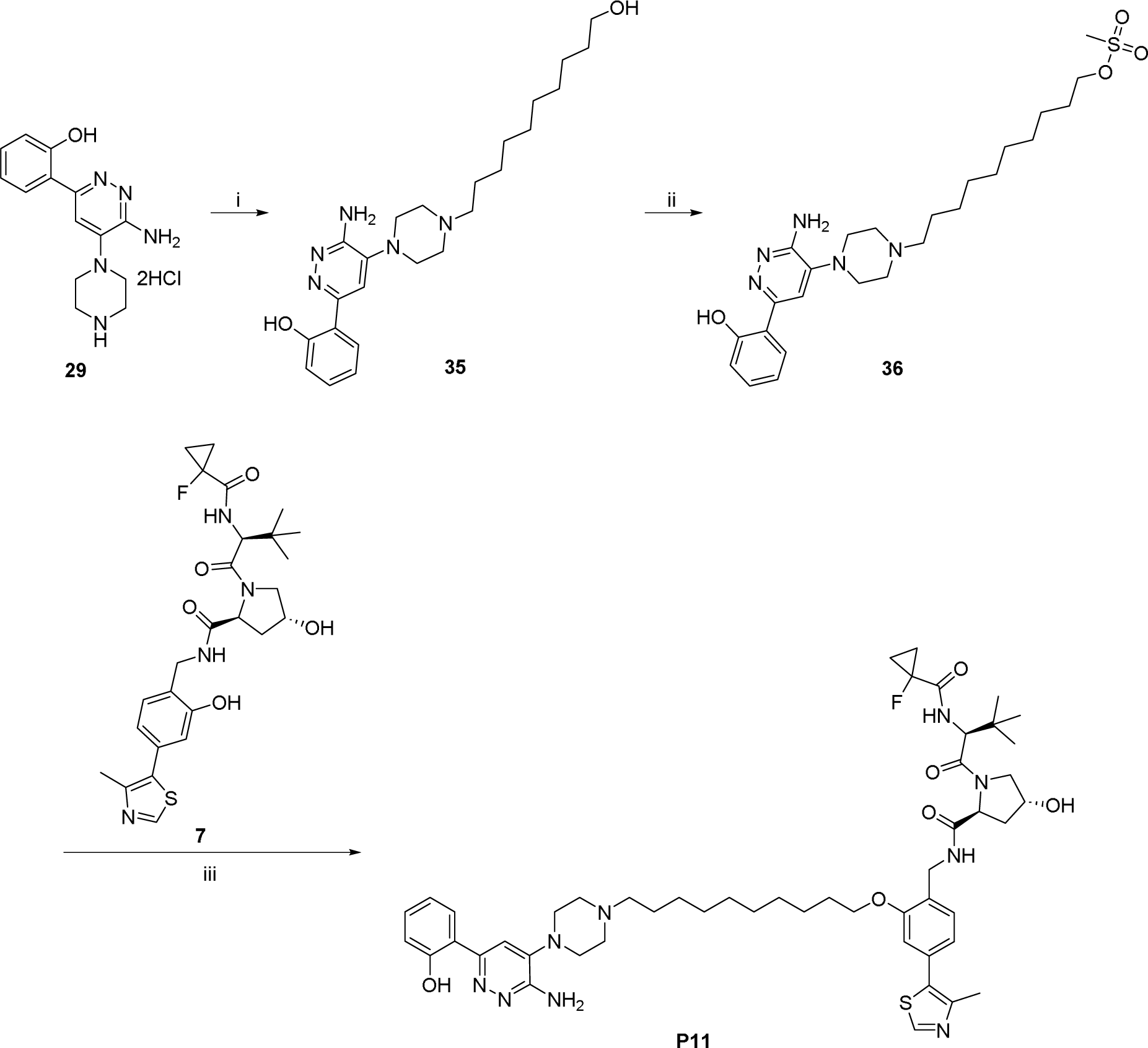
Synthesis of P11: i) 10-chlorodecan-1-ol, K_2_CO_3_, DMF, 70°C; ii) MsCl, TEA, THF, RT; iii) **7**, K_2_CO_3_, DMF, 75°C.

#### 2-{6-amino-5-[4-(10-hydroxydecyl)piperazin-1-yl]pyridazin-3-yl}phenol 35

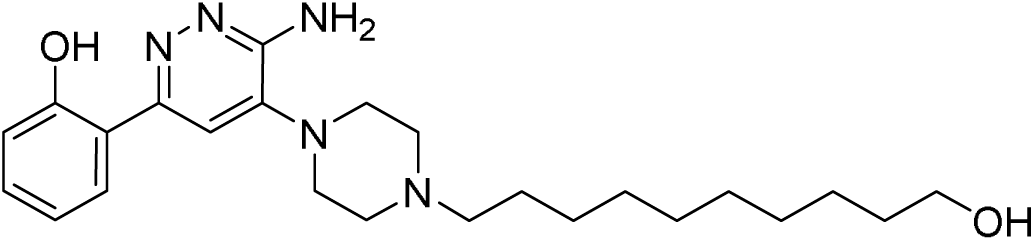

**29** (100 mg, 0.37 mmol) and 10-chlorodecan-1-ol (75 µL, 0.37 mmol, 1.0 eq.) were dissolved in DMF (500 µL) and potassium carbonate (61 mg, 0.44 mmol, 1.2 eq.) was added. The resulting mixture was stirred at 70°C for 36 h. It was diluted with MeCN and water, filtered through a syringe filter and purified by basic HPLC-chromatography to obtain **35** (32 mg, 20% yield).

#### 10-{4-[3-amino-6-(2-hydroxyphenyl)pyridazin-4-yl]piperazin-1-yl}decyl methanesulfonate 36

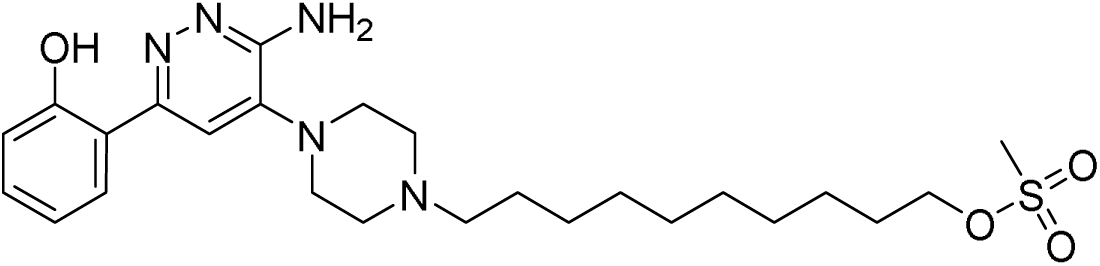

**35** (32 mg, 75 µmol) was dissolved in THF (500 µL) and TEA (13 µL, 90 µmol, 1.2 eq.) was added, followed by addition of methanesulfonyl chloride (12 µL, 0.16 mmol, 2.1 eq.). The resulting mixture was stirred at RT for 1 h. It quenched with sat. sodium bicarbonate solution, diluted with DCM and stirred for 1 h. The reaction mixture was extracted with DCM, dried over MgSO_4_ and concentrated to obtain crude **36**.

#### (2*S*,4*R*)-N-(2-((10-(4-(3-amino-6-(2-hydroxyphenyl)pyridazin-4-yl)piperazin-1-yl)decyl)oxy)-4-(4-methylthiazol-5-yl)benzyl)-1-((*S*)-2-(1-fluorocyclopropane-1-carboxamido)-3,3-dimethylbutanoyl)-4-hydroxypyrrolidine-2-carboxamide P11

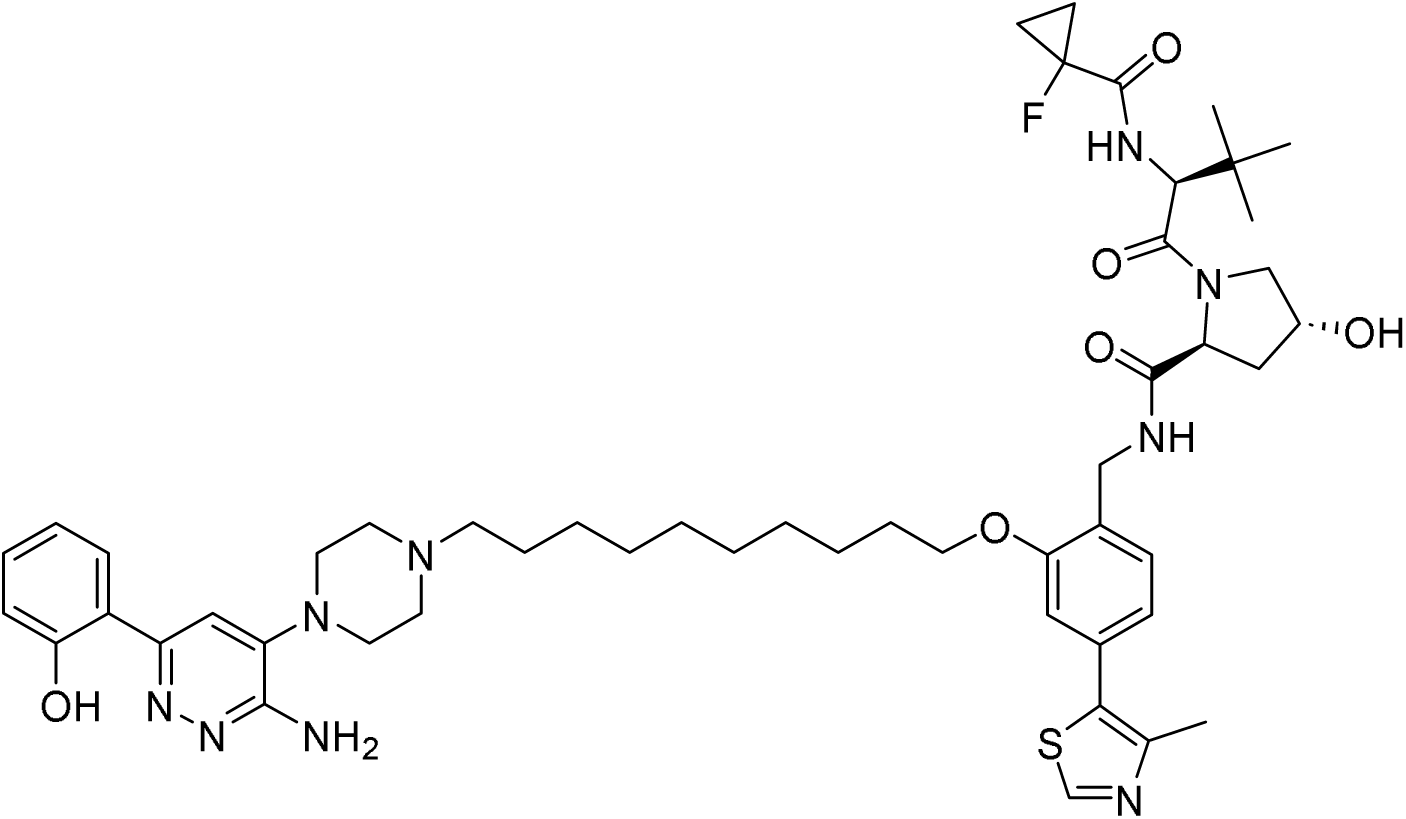

To a mixture of **7** (20 mg, 38 µmol) and potassium carbonate (16 mg, 0.11 mmol, 3.0 eq.) in DMF (200 µL), **36** (19 mg, 38 µmol, 1.0 eq), dissolved in DMF (200 µL), was added. The resulting mixture was stirred at 75°C for 1 h. It was diluted with MeCN and water, filtered through a syringe filter and purified by basic HPLC-chromatography to obtain **P11** (19 mg, 54% yield).

##### HRMS (ESI+) *m*/*z*

[M+H] ^+^ calcd for C_50_H_68_FN_9_O_6_S 942.50181; found 942.50909

##### ^1^H NMR (DMSO-d_6_) δ

14.23 (br s, 1H), 14.16 (br s, 0.1H rotamer), 8.98 (s, 1H), 8.56 (br t, J=5.5 Hz, 0.1H rotamer), 8.49 (br t, J=6.0 Hz, 1H), 7.92 (br d, J=7.5 Hz, 1H), 7.55 (s, 0.1H rotamer), 7.51 (br s, 1H), 7.40 (d, J=7.7 Hz, 1H), 7.29 (dd, J=9.2, 2.4 Hz, 1H), 7.24 (t, J=8.3 Hz, 1H), 7.18 (d, J=8.1 Hz, 0.1H rotamer), 7.00 (d, J=1.3 Hz, 1H), 6.95 (dd, J=7.7, 1.3 Hz, 1H), 6.87-6.92 (range, 2H), 6.39 (s, 0.2H rotamer), 6.23 (br s, 2H), 5.17 (d, J=3.7 Hz, 1H), 5.04 (d, J=2.8 Hz, 0.1H rotamer), 4.68 (t, J=7.4 Hz, 0.1H rotamer), 4.60 (d, J=9.2 Hz, 1H), 4.52 (t, J=8.3 Hz, 1H), 4.45 (d, J=9.4 Hz, 0.1H rotamer), 4.36 (br s, 1H), 4.29 (dd, J=16.5, 6.2 Hz, 1H), 4.20 (dd, J=16.5, 5.7 Hz, 1H), 4.11 (br d, J=5.0 Hz, 0.1H rotamer), 4.02-4.07 (range, 2H), 3.65 (dd, J=6.2, 3.7 Hz, 2H), 3.58-3.63 (m, 2H), 3.01-3.21 (range, 4H), 2.54-2.61 (m, 2H), 2.45-2.47 (m, 3H), 2.35 (br d, J=5.9 Hz, 1H), 2.09 (br dd, J=12.6, 7.8 Hz, 1H), 1.93 (ddd, J=12.8, 8.7, 4.5 Hz, 1H), 1.70-1.81 (m, 2H), 1.41-1.52 (range, 4H), 1.33-1.40 (range, 4H), 1.28-1.33 (range, 8H), 1.22 (dd, J=8.3, 2.8 Hz, 2H), 0.91-1.01 (m, 9H)

##### ^13^C NMR (DMSO-d_6_) δ

172.3, 169.4, 168.6 (d, CF=20 Hz), 159.0, 158.9, 156.3, 155.1, 153.7, 151.9, 148.3, 140.8, 131.8, 131.3, 130.6, 128.1, 127.4, 126.6, 121.1, 119.0, 118.3, 117.8, 112.1, 110.8, 78.6 (d, CF=233 Hz), 69.4, 68.2, 67.5, 59.3, 58.2, 57.1, 57.0, 52.6, 48.9, 48.9, 40.8, 40.6, 40.5, 38.3, 37.7, 36.7, 36.5, 29.5, 29.3, 29.2, 29.1, 27.4, 26.7, 26.6, 26.0, 25.9, 16.5, 16.4, 13.4 (d, CF=10 Hz), 13.2 (d, CF=10 Hz)

##### HPLC

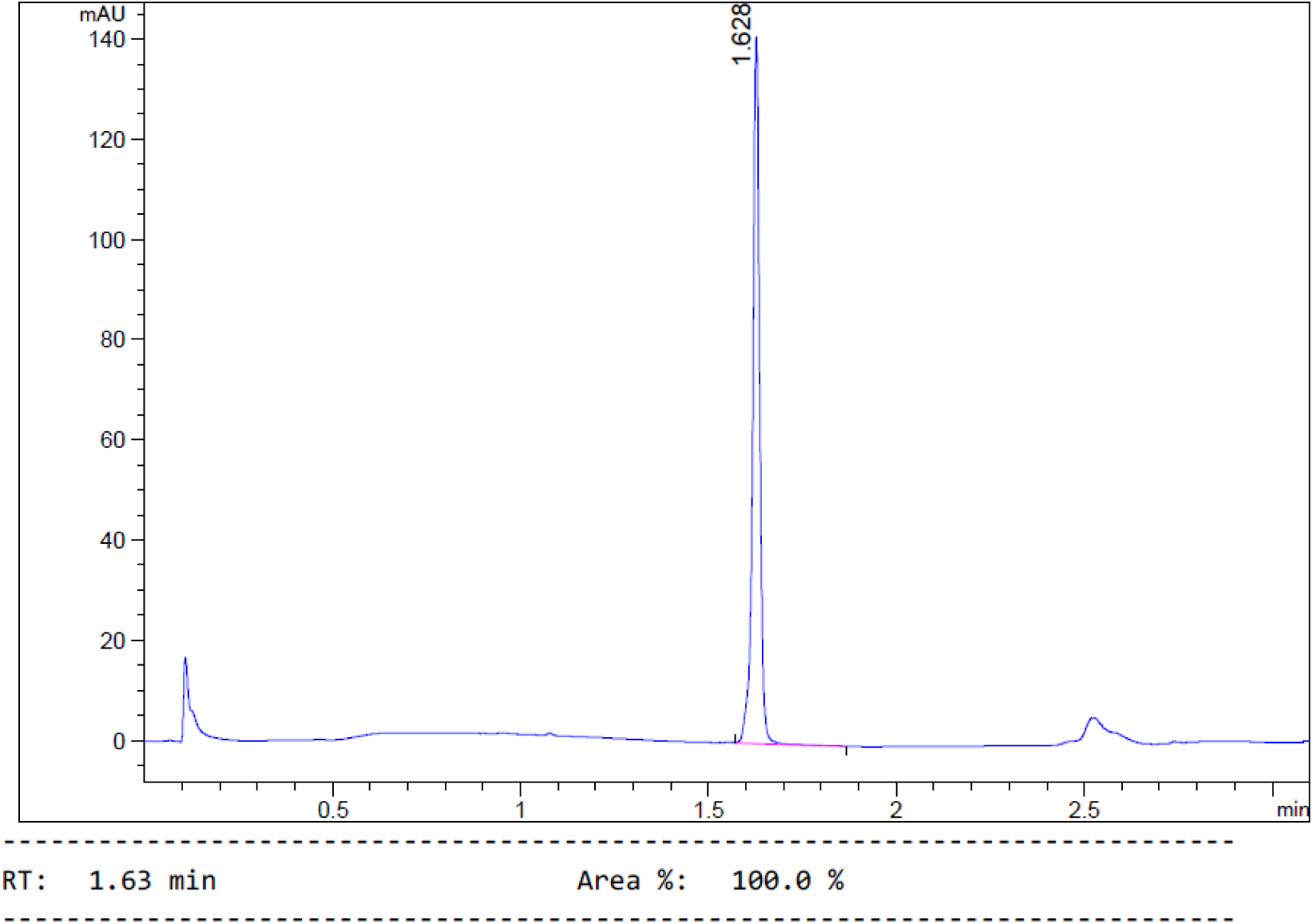

##### ^1^H and ^13^C NMR

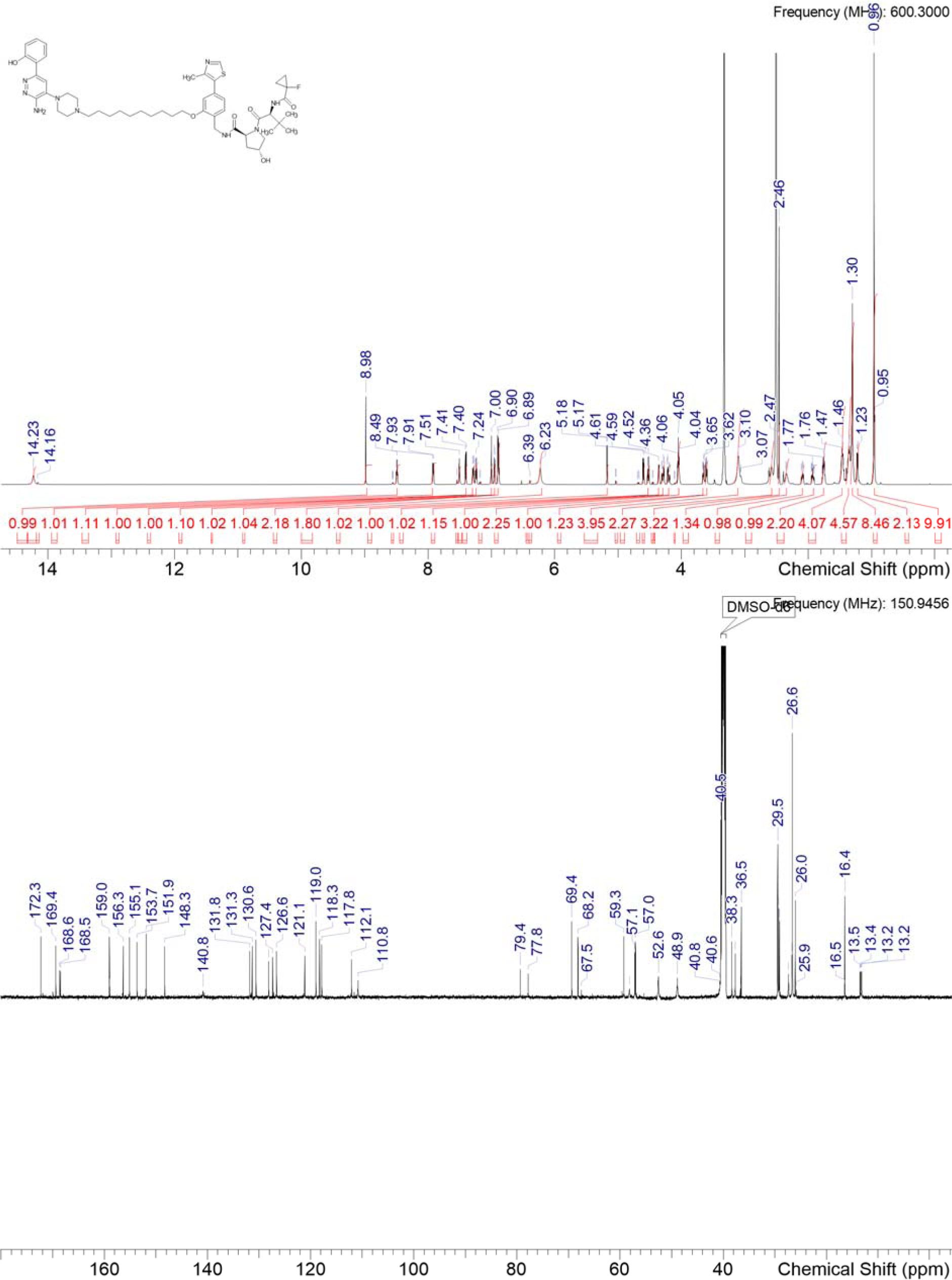

### P16

#### (2*S*,4*R*)-*N*-(2-(4-(4-(3-amino-6-(2-hydroxyphenyl)pyridazin-4-yl)piperazine-1-carbonyl)phenethoxy)-4-(4-methylthiazol-5-yl)benzyl)-1-((*S*)-2-(1-fluorocyclopropane-1-carboxamido)-3,3-dimethylbutanoyl)-4-hydroxypyrrolidine-2-carboxamide

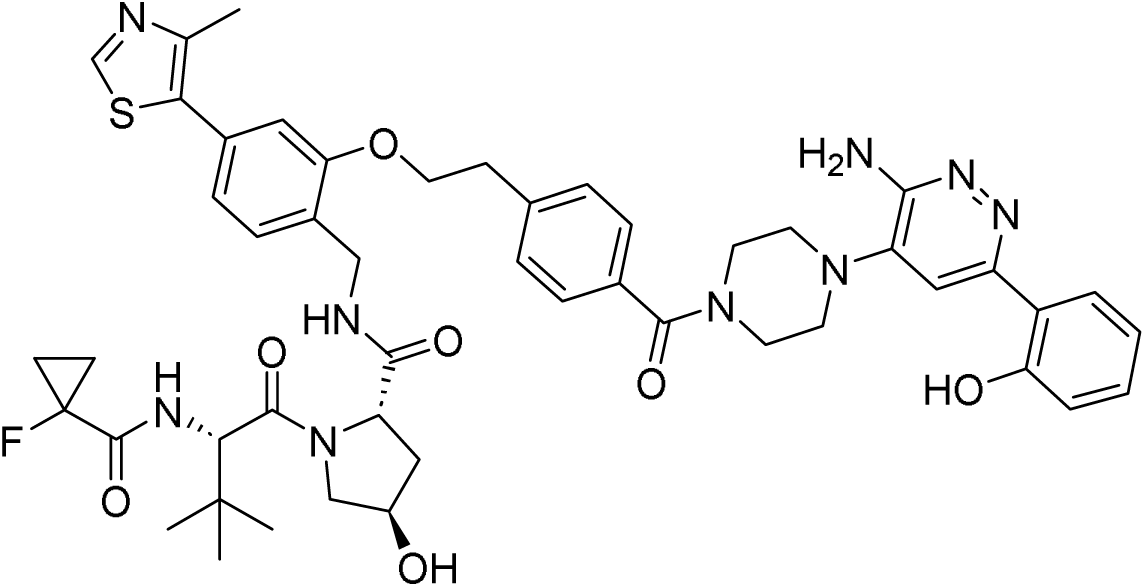

Synthesis in [2].

##### HRMS (ESI+) *m*/*z*

[M+H] ^+^ calcd for C_49_H_56_FN_9_O_7_S 934.40330; found 934.41058

##### ^1^H NMR (DMSO-d_6_) δ

14.17 (br s, 1H), 8.98 (s, 1H), 8.57 (q, J=5.4 Hz, 0.1H rotamer), 8.49 (t, J=6.0 Hz, 1H), 7.92 (dd, J=8.2, 1.2 Hz, 1H), 7.60-7.63 (m, 0.1H rotamer), 7.57 (s, 1H), 7.56 (br s, 0.1H rotamer), 7.51 (d, J=7.9 Hz, 0.1H rotamer), 7.44-7.49 (range, 2H), 7.37-7.44 (range, 3H), 7.28 (br dd, J=9.3, 2.3 Hz, 1H), 7.24 (dt, J=8.4, 1.6 Hz, 1H), 7.19 (d, J=7.7 Hz, 0.1H rotamer), 7.11 (s, 0.1H rotamer), 7.04 (s, 1H), 6.95 (dd, J=7.8, 1.0 Hz, 1H), 6.85-6.92 (range, 2H), 6.44 (s, 2H), 5.31 (s, 0.1H rotamer), 5.16 (d, J=3.5 Hz, 1H), 5.04 (d, J=2.8 Hz, 0.1H rotamer), 4.69 (t, J=7.3 Hz, 0.1H rotamer), 4.58 (d, J=9.4 Hz, 1H), 4.52 (t, J=8.3 Hz, 1H), 4.47 (br d, J=9.2 Hz, 0.1H rotamer), 4.28-4.38 (range, 3H), 4.24 (dd, J=16.6, 6.3 Hz, 1H), 4.12 (dd, J=16.6, 5.4 Hz, 1H), 4.08 (br d, J=4.8 Hz, 0.1H rotamer), 3.78-3.96 (range, 2H), 3.54-3.68 (range, 4H), 3.51 (s, 0.2H rotamer), 3.46-3.50 (m, 0.2H rotamer), 3.05-3.20 (range, 6H), 2.46 (s, 3H), 2.38 (s, 0.1H rotamer), 2.09 (br dd, J=12.6, 7.8 Hz, 1H), 1.85-1.97 (m, 1H), 1.29-1.44 (m, 2H), 1.18-1.28 (m, 2H), 0.97 (s, 0.55H rotamer), 0.92-0.95 (m, 9H)

##### ^13^C NMR (DMSO-d_6_) δ

172.3, 169.6, 169.4, 168.5 (d, CF=20 Hz), 159.0, 156.1, 155.3, 153.6, 151.9, 148.3, 141.0, 140.5, 134.1, 131.7, 131.4, 130.6, 129.6, 128.1, 127.6, 127.4, 126.6, 121.3, 118.9, 118.2, 117.9, 112.1, 111.7, 78.6 (d, CF=233 Hz), 69.4, 68.7, 59.3, 57.2, 57.0, 49.2, 47.2, 41.7, 38.4, 37.7, 36.5, 35.3, 26.6, 16.5, 13.4 (d, CF=10 Hz), 13.2 (d, CF=10 Hz)

##### HPLC

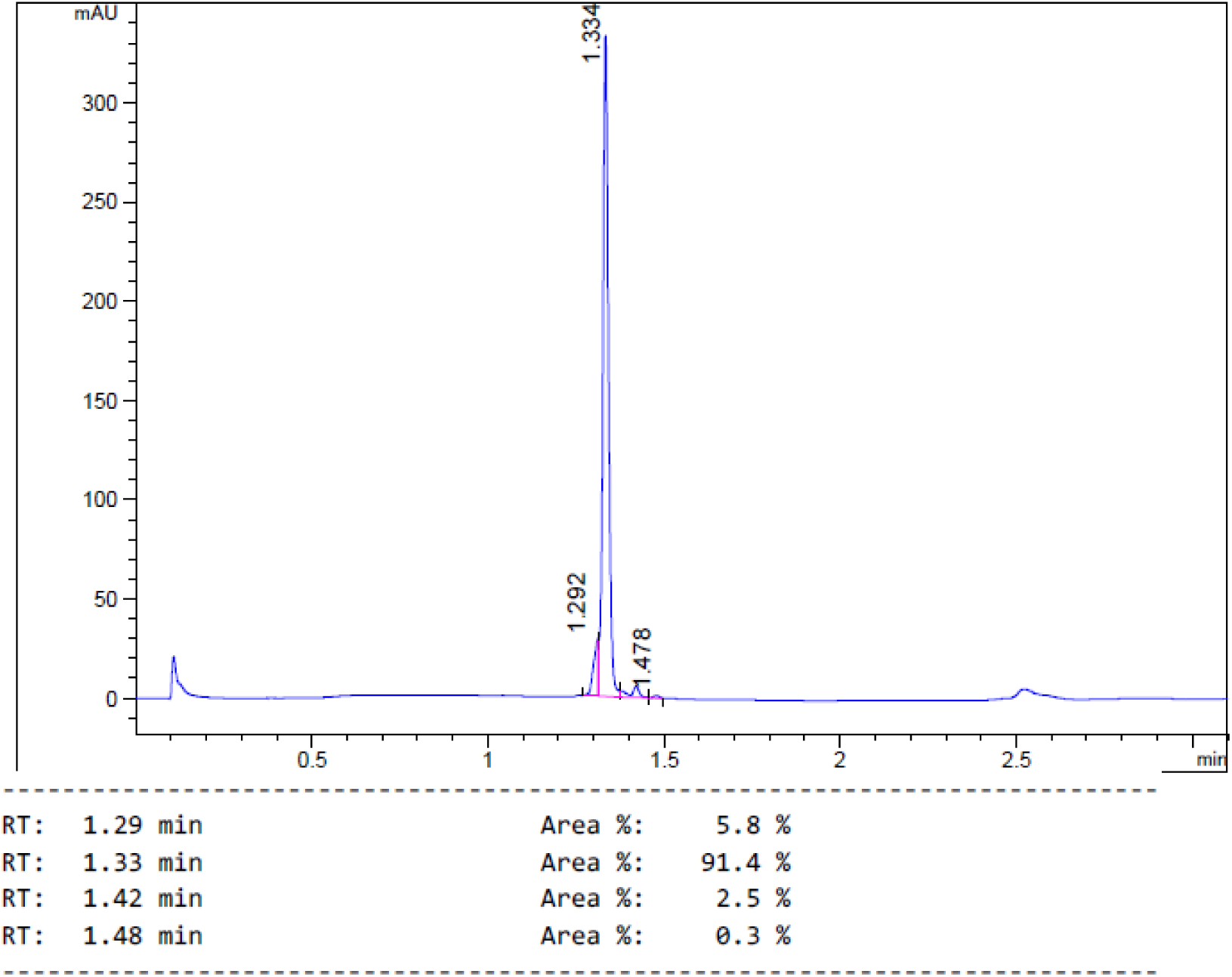

##### 1H and ^13^C NMR

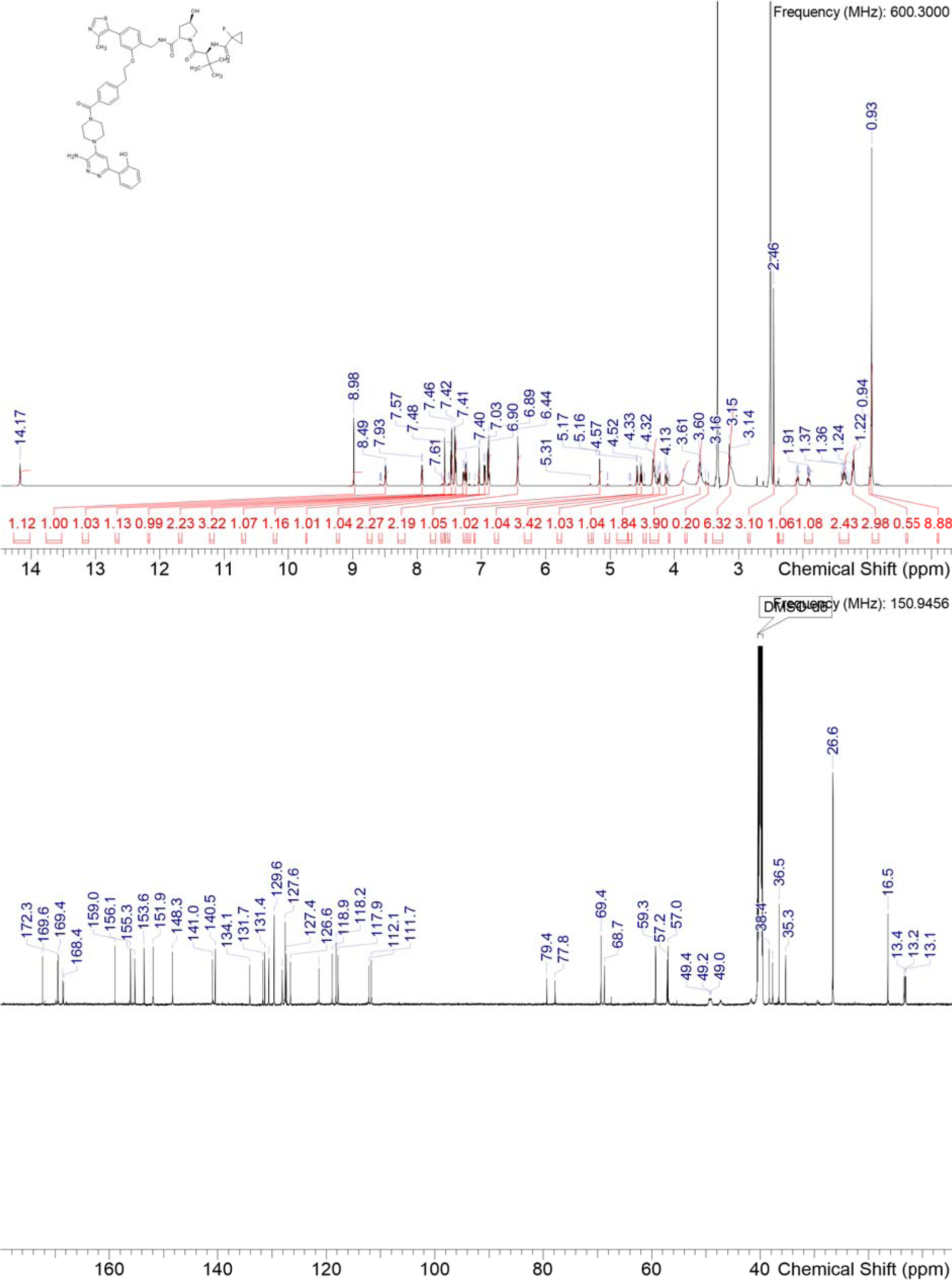

### Synthesis of P18

**Scheme S9:**
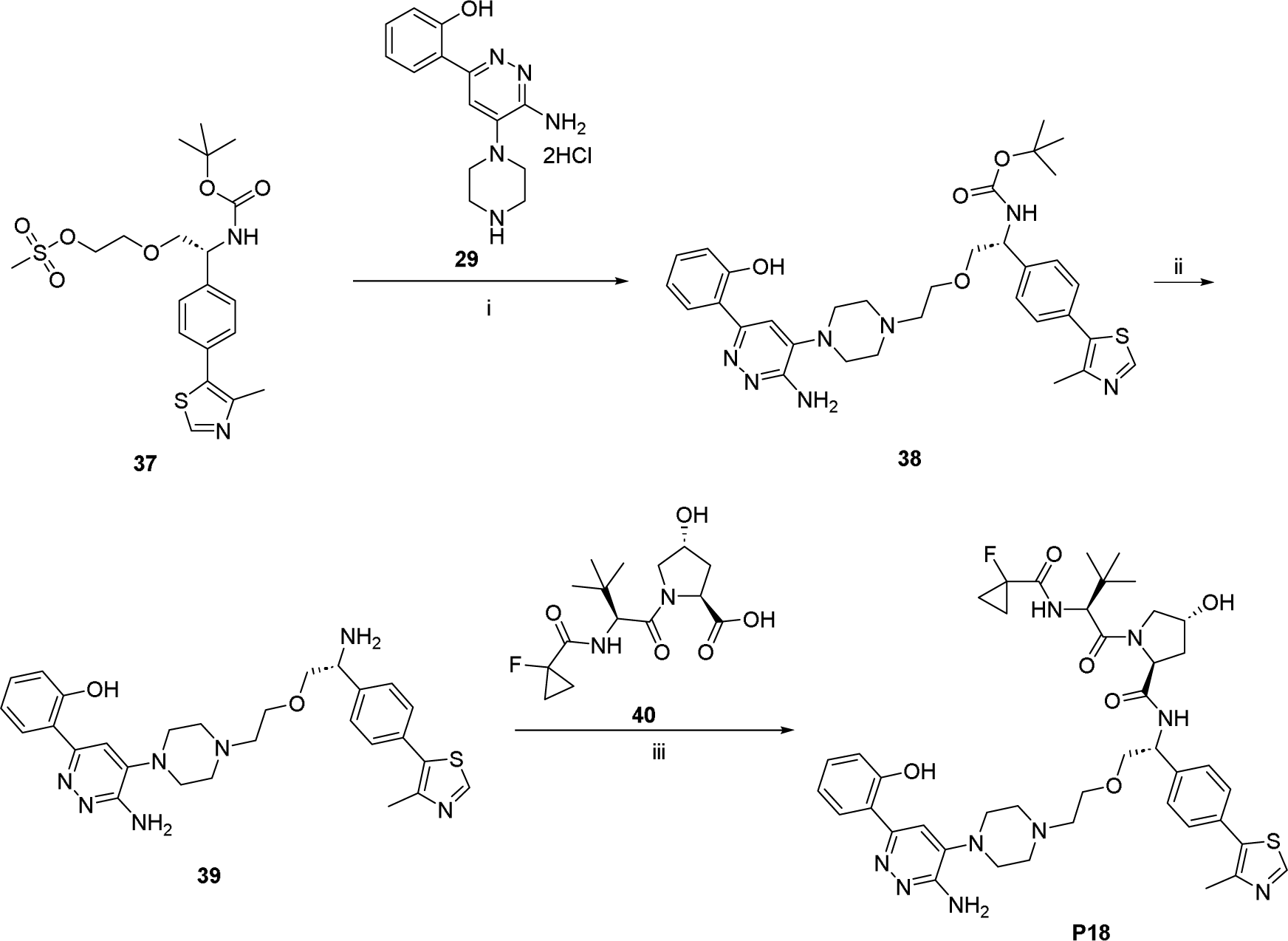
Synthesis of P18: i) **29**, DIPEA, DMF, 80°C; ii) conc. HCl, MeOH, RT; iii) **40**, HATU, TEA, DMF, RT.

#### *tert*-butyl (*R*)-(2-(2-(4-(3-amino-6-(2-hydroxyphenyl)pyridazin-4-yl)piperazin-1-yl)ethoxy)-1-(4-(4-methylthiazol-5-yl)phenyl)ethyl)carbamate 38

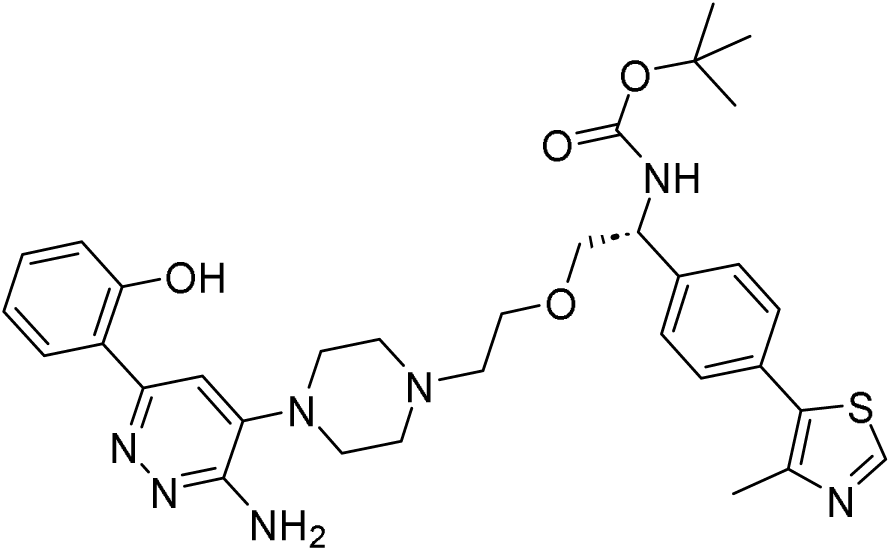

**37** (35 mg, 77 µmol) and **29** (21 mg, 77 µmol, 1.0 eq.) were dissolved in DMF (1000 µL) and DIPEA (66 µL, 0.38 mmol, 5 eq.) was added. The mixture was stirred at 80°C for 14 h. It was diluted with MeCN and water, filtered through a syringe filter and purified by basic HPLC-chromatography to obtain **38** (23 mg, 48% yield).

#### (2*S*,4*R*)-N-((*R*)-2-(2-(4-(3-amino-6-(2-hydroxyphenyl)pyridazin-4-yl)piperazin-1-yl)ethoxy)-1-(4-(4-methylthiazol-5-yl)phenyl)ethyl)-1-((*S*)-2-(1-fluorocyclopropane-1-carboxamido)-3,3-dimethylbutanoyl)-4-hydroxypyrrolidine-2-carboxamide P18

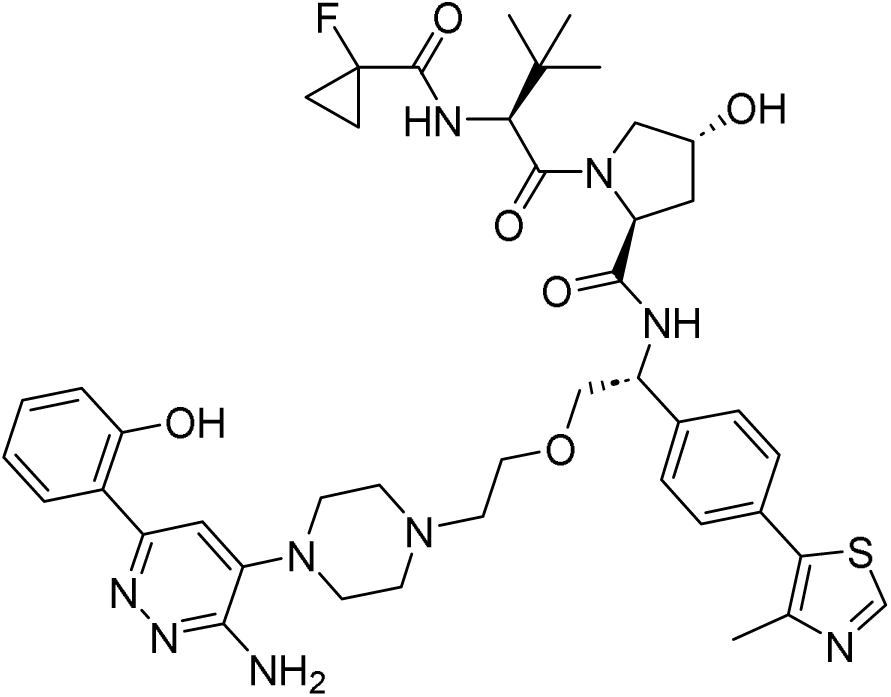

**38** (23 mg, 36 µmol) was taken up in methanol (1000 µL) and conc. aq. HCl (400 µL) was added. The mixture was stirred at RT for 1 h. The solvents were removed under reduced pressure to obtain crude **39**.

**40** (14 mg, 41 µmol, 1.2 eq.) was dissolved in DMF (500 µL) and TEA (21 µL, 0.14 mmol, 4.0 eq.) and HATU (41 mg, 0.11 mmol, 3.0 eq.) were added. The resulting mixture was stirred at RT for 15 minutes, then **39** (19 mg, 36 µmol) was added and stirring was continued at RT for 3 h. The reaction was diluted with water, filtered through a syringe filter and purified by basic HPLC-chromatography to obtain **P18** (17 mg, 56% yield).

##### HRMS (ESI+) *m*/*z*

[M+H] ^+^ calcd for C_43_H_54_FN_9_O_6_S 844.39134; found 844.39862

##### ^1^H NMR (DMSO-d_6_) δ

14.16 (br s, 1H), 8.92 (s, 0.1H rotamer), 8.87 (s, 1H), 8.86 (s, 0.1H rotamer), 8.70 (br d, J=7.9 Hz, 0.1H rotamer), 8.49 (br d, J=7.7 Hz, 0.1H rotamer), 8.46 (d, J=7.9 Hz, 1H), 7.80 (d, J=7.2 Hz, 1H), 7.35-7.42 (range, 5H), 7.32-7.35 (m, 0.3H rotamer), 7.13-7.20 (range, 2H), 6.91 (dd, J=9.0, 2.6 Hz, 0.1H rotamer), 6.79-6.86 (range, 2H), 6.12 (s, 2H), 5.08 (d, J=3.5 Hz, 1H), 4.93-5.00 (m, 1H), 4.84-4.92 (m, 0.1H rotamer), 4.69 (t, J=7.4 Hz, 0.1H rotamer), 4.51 (d, J=9.2 Hz, 1H), 4.47 (t, J=8.3 Hz, 1H), 4.33 (d, J=9.0 Hz, 0.1H rotamer), 4.20-4.25 (m, 1H), 4.16-4.20 (m, 0.1H rotamer), 3.75 (dd, J=10.1, 4.8 Hz, 0.1H rotamer), 3.65 (s, 0.1H rotamer), 3.63 (br d, J=2.8 Hz, 0.1H rotamer), 3.58-3.62 (m, 1H), 3.47-3.57 (range, 5H), 3.44 (t, J=6.1 Hz, 0.1H rotamer), 3.40 (br d, J=12.5 Hz, 0.1H rotamer), 2.88-3.11 (range, 4H), 2.62 (s, 0.1H rotamer), 2.51-2.59 (range, 4H), 2.49 (br t, J=4.7 Hz, 2H), 2.39 (s, 0.3H rotamer), 2.37 (s, 3H), 2.06 (s, 0.28H rotamer), 2.02 (br t, J=10.3 Hz, 1H), 1.79-1.85 (m, 0.1H rotamer), 1.71 (ddd, J=12.9, 8.7, 4.5 Hz, 1H), 1.18-1.35 (m, 2H), 1.08-1.18 (m, 2H), 0.89-0.93 (m, 9H)

##### ^13^C NMR (DMSO-d_6_) δ

171.4, 169.3, 168.5 (d, CF=20 Hz), 159.0, 155.1, 153.6, 152.0, 148.3, 141.2, 140.9, 131.5, 130.6, 130.6, 129.1, 127.9, 126.6, 118.9, 118.3, 117.9, 110.7, 78.6 (d, CF=231 Hz), 73.4, 69.3, 68.8, 59.1, 57.5, 57.1, 57.0, 53.0, 52.6, 49.0, 38.2, 36.6, 26.7, 16.5, 13.4 (d, CF=10 Hz), 13.1 (d, CF=10 Hz)

##### HPLC

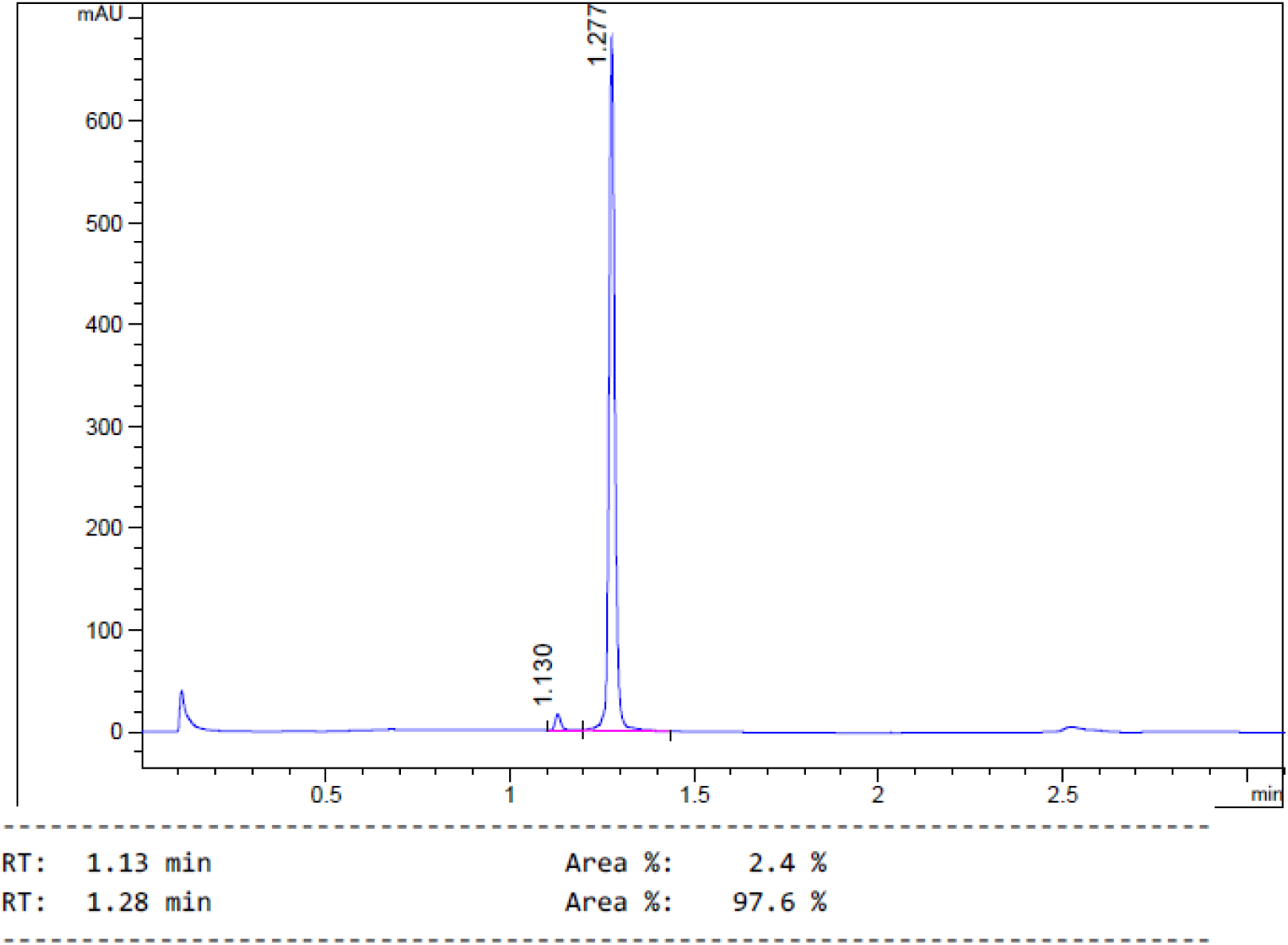

##### 1H and ^13^C NMR

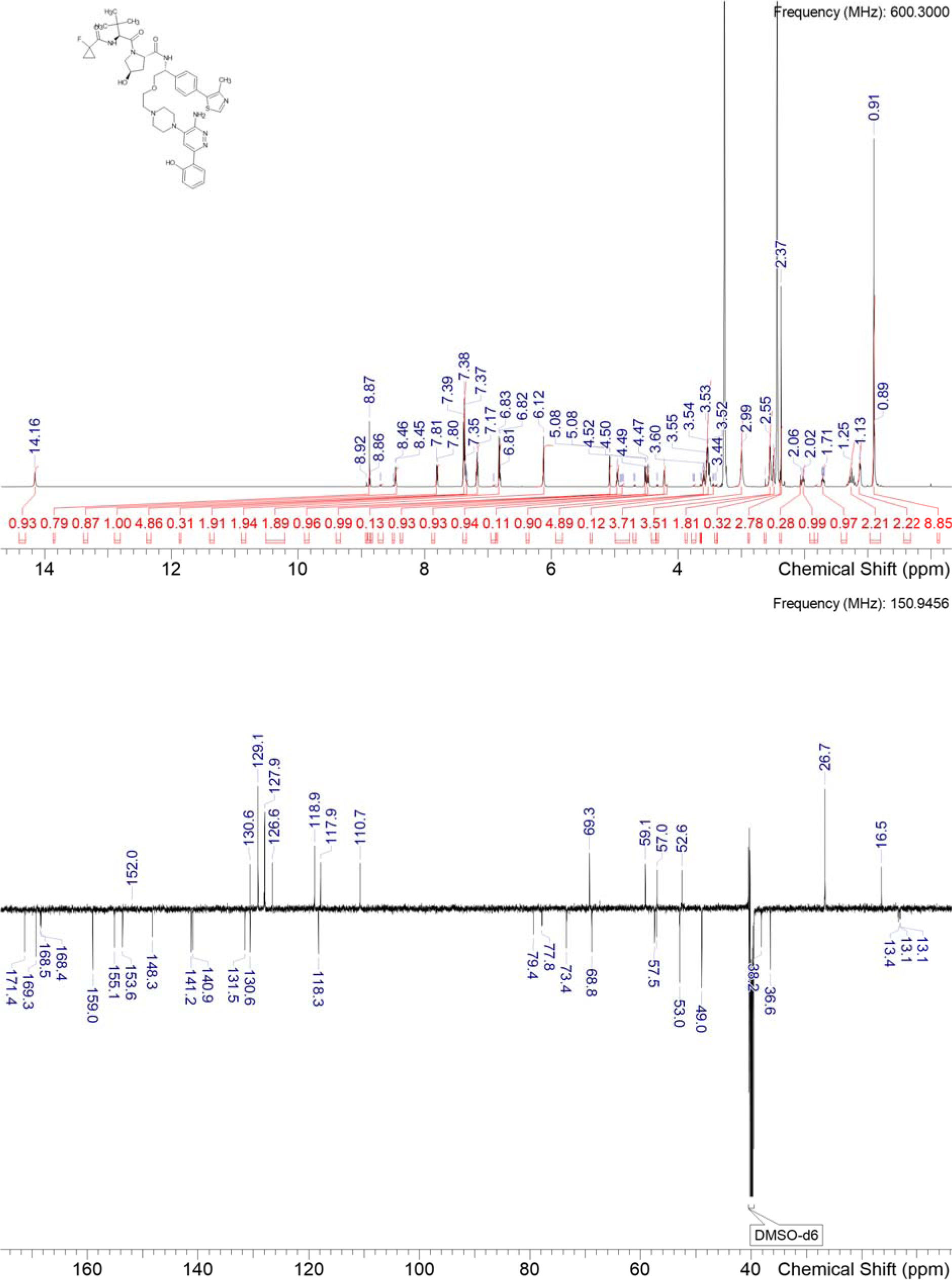

